# Genomic, phenotypic and environmental correlates of speciation in the midwife toads (*Alytes*)

**DOI:** 10.1101/2024.10.24.619835

**Authors:** Johanna Ambu, Spartak N Litvinchuk, Carlos Caballero-Díaz, Alfredo Nicieza, Guillermo Velo-Antón, Helena Gonçalves, Fernando Martínez-Freiría, Helena Martínez-Gil, Juan Francisco Beltrán, David Donaire-Barroso, Axel Hernandez, Tomasz Suchan, Pierre-André Crochet, ĺñigo Martínez-Solano, Christophe Dufresnes

## Abstract

Speciation, i.e., the formation of new species, implies that diverging populations evolve genetic and phenotypic factors that promote reproductive isolation (RI), but the adaptive vs. neutral origin of these factors and their relative contributions across the speciation continuum remain elusive. Here we test which of genomic, bioacoustic, morphological and environmental differentiation best predicts RI across the midwife toads (genus *Alytes*), a diversification of Mediterranean amphibians. Hybrid zone analyses in the *A. obstetricans* complex support that without strong geographic barriers to dispersal, the extent of introgression (which should reflect the strength of RI) covaries with genomic divergence irrespective of other factors. Phenotypic divergence become important later along the continuum, namely between non-admixing species attributed to distinct subgenera. Our results suggest that by putatively causing intrinsic incompatibilities in hybrids, the genetic mutations accumulating randomly between allopatric populations act as the initial trigger of RI, while substantial ecological and behavioral differentiation is a long-term consequence of species divergence that ultimately promotes sympatry. Whereas speciation is usually claimed to be primarily adaptive, our study corroborates recent findings that new species may also be a neutral outcome of gradual phylogeographic divergence, which has practical implications for species delimitation.

## Introduction

Speciation, i.e., the formation of species, is a multifaceted process involving an array of behavioral, ecological, geographic and genetic factors, and understanding how these factors interact through time and space in shaping Earth’s amazing biodiversity is one of the oldest and most fascinating topics in evolutionary biology (Coyne & Orr, 2004; Nosil, 2012; Mérot et al., 2017; Nosil et al., 2017, 2021). What best characterizes a species is also a very practical issue, as it conditions how populations are considered in taxonomy and thereby acknowledged in conservation policies and by society in general (Thomson et al., 2018; Vences et al., 2024). The diversity and complexity of speciation mechanisms is partly why scientists struggle to agree on the way to operationally delimit species (Zachos, 2016; Conix et al., 2023; Dufresnes et al., 2023).

It is generally accepted that species correspond to genealogical lineages that have differentiated following a reduction of gene flow (de Queiroz, 2007), to the point that they may no longer fuse back when given the chance to hybridize during secondary contact (Hillis, 2020; Stankowski & Ravinet, 2021; Dufresnes et al., 2023; Vences et al., 2024). Accordingly, taxonomic concepts and criteria have either brought the emphasis on the process leading to speciation (e.g., reproductive barriers; “biological” species, Mayr, 1942; Coyne & Orr, 2004) or on its perceived outcomes (e.g., phylogenetic divergence: “phylogenetic” species, de Queiroz, 1998; phenotypic divergence: “morphological” species). Both aspects are expected to increase with time, and speciation is accordingly viewed as a gradual phenomenon across the whole animal kingdom (Hedges et al., 2015; Roux et al., 2016; Coughlan & Matute, 2020; but see Ronco & Salzburger, 2021). Jointly documenting patterns of reproductive isolation (RI), genetic divergence, phenotypic and ecological differentiation can thus provide clues on the pace of the speciation clock, and on the mechanisms ticking that clock at different stages of the continuum.

In this respect, two frequently asked questions in speciation and species delimitation research are (1) when does speciation start (Butlin & Faria, 2024)? and (2) is its onset primarily the product of neutral or adaptive processes (Matsubayashi & Yamaguchi, 2022, Black et al., 2024)? The answers to these questions should depend on the mode by which speciation progresses. Ecological speciation, the most accepted *modus operandi* for the evolution of new species, implies that disruptive natural or sexual selection on locally-adapted phenotypes may rapidly promote pre-zygotic or extrinsic post- zygotic barriers. RI driven by ecological, behavioral or morphological differences thus initiates genetic divergence between diversifying populations, starting with genomic regions putatively bearing adaptive polymorphism (Schluter, 2009; Servedio et al., 2011; Nosil, 2012; Arnegard et al., 2014; Butlin & Faria, 2024). On the other hand, intrinsic post-zygotic barriers can arise from genomic incompatibilities such as Dobzhansky-Muller incompatibilities (DMIs) (Muller, 1940; Turelli & Orr, 1995), following the random accumulation and fixation by drift of otherwise neutral mutations as lineages progressively diverge (Orr, 1995; Orr & Turelli, 2001; Matute et al., 2010; Moyle & Nakazato, 2010). Such genetic divergence may thus initiate intrinsic post-zygotic RI, with ecological and behavioral barriers eventually arising later along the continuum (Dufresnes et al., 2021b; Dufresnes & Crochet, 2022). Hence, genetic divergence can either be the cause or the consequence of RI between incipient species, depending on whether speciation is initially neutral or adaptive.

Studying the gray zone of speciation, i.e., the window of divergence during which reproductive barriers are still incomplete (Roux et al., 2016), can be informative to solve this chicken-and-egg situation and disentangle the adaptive and neutral processes contributing to the formation of species. In particular, hybrid zones, i.e., geographic areas where diverging lineages meet and hybridize, offer open world experiments to quantify RI *in statu nascendi*, by documenting patterns of hybridization and gene flow (Hewitt, 1988; Barton & Hewitt, 1989; Harrison, 1993; Payseur, 2010). If speciation starts with pre-zygotic or extrinsic post-zygotic barriers, nascent species should bear diagnostic differences at the behavioral or ecological traits relevant to these barriers, and admixture across hybrid zones should be buffered by their degree of phenotypic or environmental differentiation rather than their genetic distance. On the other hand, if RI primarily results from molecular divergence, admixture across hybrid zones should essentially depend on genetic distances, and nascent species can remain phenotypically similar (i.e., cryptic species). Understanding whether species are first determined by genetic or phenotypic factors can in turn inform taxonomists on the most relevant criteria when delimiting species under integrative taxonomy (Padial et al., 2010).

Moreover, the potential relationships between RI and differentiation at molecular and phenotypic traits can further serve to make predictive inferences on species status when contact zones are not accessible (Dufresnes et al., 2021b, 2023; Vences et al., 2024).

Comparing hybridization patterns, genetic divergence, and phenotypic differentiation in a single comprehensive framework is no small feat. Given the amount of resources needed to analyze just a single hybrid zone, comparative studies usually compile datasets from the literature, sometimes pertaining to unrelated organisms, both to increase sample sizes and provide a broader scope (McEntee et al., 2020; Dufresnes et al., 2021b). Although informative of global trends, such meta-analyses can hardly control for the many biological and methodological factors that may blur the relationships, notably those affecting gene flow across hybrid zones (e.g., dispersal capabilities and opportunities), the accuracy to quantify it (e.g., type and numbers of loci), and the comparability of divergence estimates at molecular and phenotypic traits (e.g., obtained from different sources and approaches). For instance, species age, which is often used as a measure of relative divergence in biological studies (Kumar et al., 2022), can markedly differ between nuclear and mitochondrial trees (e.g., due to cyto-nuclear discordance), between phylogenies constrained with different time calibrations, and may be irrelevant when tree topologies are not robustly supported or when taxa are the product of reticulate evolution (Ambu et al., 2023). Given these challenges, available introgression-divergence correlations relied on ordinal rather than quantitative statistics (e.g., Dufresnes et al., 2019a, 2019b, 2020), or expectedly left a large proportion of variance unexplained (e.g., Morgan-Richards & Wallis, 2003; Arntzen et al., 2014; McEntee et al., 2020; Pulido-Santacruz et al., 2020; Dufresnes et al., 2021b). The best way to alleviate this “background noise” is to reframe quantitative comparisons to a smaller scale, namely in species groups of known life-histories, featuring numerous lineages and hybrid zones, and by inferring introgression and divergence in a single batch of analyses (e.g., Singhal & Moritz, 2013; Pabijan et al., 2017).

Midwife toads of the genus *Alytes* tick all of these boxes. Widely distributed across the Western Mediterranean region, *Alytes* are notorious for their parental care behavior – the male carries the eggs on its back for several weeks until the larvae are ready to hatch (Speybroeck et al., 2016).

This remarkable innovation reduces the reliance on water compared to aquatic egg-laying species and enhances protection during the sensitive spawning and early larval stages (Lange et al., 2022). Hence, midwife toads have colonized a broad spectrum of terrestrial habitats, some species even overcoming landscape features that act as strong dispersal barriers in other amphibians (e.g., the Pyrenean mountains). Following two decades of phylogeographic research (Martínez-Solano et al., 2004; Gonçalves et al., 2007; Maia-Carvalho et al., 2018; Dufresnes & Hernandez, 2021; Lucati et al., 2022), we recently reassessed the evolution of the genus by genomic analyses (Ambu et al., 2023).

These analyses solved long-standing phylogenetic discordances between molecular markers and confirmed ten deeply divergent lineages corresponding to described taxa, presently delimited in six species, and arranged in three major clades given as distinct subgenera (summarized in Fig. 1).

**Fig. 1:**
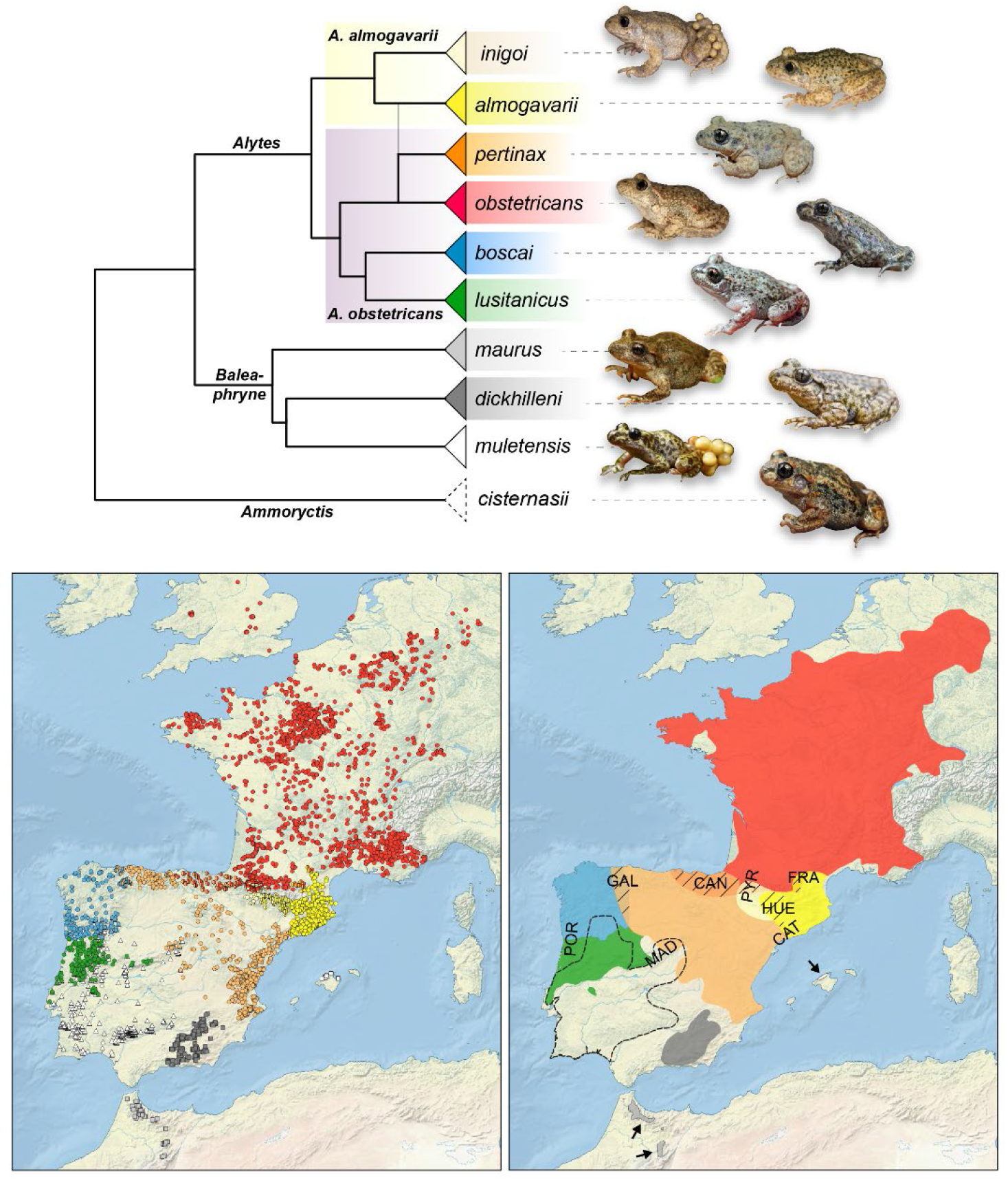
Evolution, distribution and contact zones in the *Alytes* genus. The tree reflects phylogenomic analyses adapted from Ambu et al. (2023); the putative hybrid origin of *A. o. pertinax* is emphasized. The three subgenera (branch labels) and the two species of subgenus *Alytes* / the *A. obstetricans* complex (faded squares) are identified. The left map shows occurrence records assigned to taxa based on our results for the *A. obstetricans* complex (mixed circles: admixed localities). The right map shows taxa distributions based on these records, with emphasis of the analyzed contact zones in the *A. obstetricans* complex. Arrows point to the geographically-restricted *A. maurus* and *A. muletensis*.

The subgenus *Alytes*, often referred to as the *A. obstetricans* complex, is of particular interest as it comprises six Plio-Pleistocene lineages that diversified amid frequent hybridization, even leading to extensive cyto-nuclear discordance and reticulate evolution (Fig. 1). These lineages have continuous distributions and are directly in contact in their areas of parapatry (Maia-Carvalho et al., 2018), with the most diverged ones forming steep hybrid zones consistent with incomplete but advanced stages of speciation (e.g., Dufresnes & Martínez-Solano, 2020; Ambu & Dufresnes, resubmitted). Current taxonomic accounts accordingly divided the complex into two species, *A. obstetricans* and *A. almogavarii* (Speybroeck et al., 2020; Frost, 2024), which respectively include four and two lineages preliminarily attributed to subspecies (Ambu et al., 2024). High intra- and inter- specific variation in habitat use, climatic tolerance, mating calls and morphology have been reported across the genus, but how this variation relates to speciation, local adaptation or even plasticity remains largely unanswered, notably because many populations were misassigned in previous accounts (Dufresnes & Hernandez, 2021; Ambu et al., 2023). Midwife toads thus offer a rare system to study the genetic, ecological and phenotypic determinants of speciation along a continuum of divergence spanning multiple pairs of related taxa, for which a comprehensive phylogeographic framework is now available.

Our study investigates whether RI primarily stems from genomic divergence or phenotypic and ecological differentiation in the *Alytes* genus using a total evidence approach. Specifically, we analyzed population genomics data obtained with double-digest restriction-associated DNA sequencing (ddRAD-seq) for more than 400 individuals, with a particular focus on hybrid zones between eight pairs of lineages in the *A. obstetricans* complex. We then compiled morphometric measurements, audio recordings of mating calls (the primary reproductive cue of anurans, Köhler et. al. 2017), and occurrence datasets for species distribution modelling to compare the external characteristics and environmental envelope of each lineage. If midwife toads initiated speciation by neutral genetic factors, RI between lineages should mostly depend on their genome-average divergence, with nascent species featuring steep hybrid zones irrespective of their phenotypic differentiation. On the other hand, if speciation is initially driven by adaptive factors, such as behavioral or ecological divergence, these should affect admixture between lineages irrespective of their genetic distances.

## Methods

### Lineage distribution and hybrid zone genomics

We analyzed new ddRADseq data for 332 samples (File S1), of which 127 were included in previous studies (Ambu et al., 2023; Ambu & Dufresnes, resubmitted). These samples represent all lineages/species of the genus with a particular effort to cover their sympatric/parapatric ranges along geographic transects. Genomic DNA was isolated from buccal swabs (adults; stored at -20°C) and tail tips (tadpoles; stored in 70% ethanol) using an *ad hoc* salt protocol (Bruford et al., 1992), and checked for yield and integrity on agarose gels. Genomic libraries were prepared following a custom protocol (dx.doi.org/10.17504/protocols.io.kxygx3nzwg8j/v1) adapted from Brelsford et al. (2016), and were sequenced paired-end on an Illumina NextSeq 550, using either the 2×75bp or 2×150bp High-Output kit. Paired-end reads were demultiplexed with STACKS 2.59 (Catchen et al., 2013) and trimmed to 65bp for all samples. We then used the denovo_map.pl pipeline of STACKS for RAD loci construction, assembly, and cataloging (default -m, -n, and -M values). The STACKS catalog contained 540,223 loci, with an average coverage of 17.6 reads (5.4–45.4, SD = 5.9). The module *population* of STACKS was used to export SNP and allele frequency datasets for the downstream analyses detailed below. File S1 highlights the samples used in each analysis (datasets: Ambu & Dufresnes, 2024).

To examine global patterns of genetic structure in the *A. obstetricans* complex, we obtained a matrix of 5,111 unlinked SNPs for 314 samples from 163 localities, by retaining RAD tags sequenced in at least 150 localities (-*p* 150), in at least half of the samples from each locality (-*r* 0.5), and by randomly choosing a single SNP per RAD tag (-*write-random-snp*). The dataset was analyzed in STRUCTURE 2.3.4 (Pritchard et al., 2000) through runs of 200,000 iterations (20,000 burnin) for *K* = 1–10, with a particular emphasis for *K* = 6 to distinguish lineages/subspecies (Ambu et al., 2023). As a complement, we considered the clustering results of a previous ddRAD-seq study in northeastern Spain, namely the nuclear ancestries for *K* = 2 of 111 *almogavarii*/*pertinax* samples (18 localities), inferred from 433 SNPs (Dufresnes & Martínez-Solano, 2020).

Separate analyses were conducted on six contact zones of the *A. obstetricans* complex (CAN, HUE, POR, MAD, GAL, PYR, Fig. 1). In each case, we selected a subset of samples encompassing parental and parapatric localities, and featuring less than 50% of missing data in the global dataset above. A SNP matrix was obtained by retaining the RAD tags sequenced in all localities of the subset (-*p* number of localities), in at least half of the samples from each locality (-*r* 0.5), and by randomly choosing a single SNP per RAD tag (-*write-random-snp*). The matrix was analyzed in a STRUCTURE run (same parameters as above) with *K* = 2 to compute the average population ancestry to each lineage (*Q_pop_*). The geographic extent of gene flow was then quantified by fitting sigmoid clines to *Q_pop_* for populations sampled along the geographic transect. To this end, we used the *R* package *hzar* (Derryberry et al., 2014), choosing the cline model with two parameters (width *w* and center *c*), which allows to compare estimates across the different hybrid zones without overparameterization. We further characterized the heterogeneity of gene flow throughout the genome by fitting sigmoid clines to allele frequency data at lineage-diagnostic SNPs, i.e., with fixed alleles between the pure populations of each lineage. Two additional hybrid zones (FRA and CAT) were incorporated based on previous analyses similarly performed by Dufresnes & Martínez-Solano (2020) and Ambu & Dufresnes (resubmitted).

Lastly, we investigated whether the southern lineages of the *A. obstetricans* complex (*A. o. lusitanicus*, *A. o. pertinax*) still hybridize and admix with other *Alytes* species occurring within or close to their ranges (*A. cisternasii*, *A. dickhilleni*). To this end, we selected two additional subsets of samples from which we obtained two matrices of unlinked SNPs: one restricted to the RAD tags shared between *A. cisternasii*, *A. o. lusitanicus* and *A. o. pertinax* from Central Spain (-*p* 3), present in at least half of the samples in each group (-*r* 0.5), and randomly choosing a single SNP per RAD tag (-*write-random-snp*); and one restricted to the RAD tags shared between *A. dickhilleni* and *A. o. pertinax* from SE-Spain (-*p* 2), present in at least half of the samples in each group (-*r* 0.5), and also randomly choosing a single SNP per RAD tag (-*write-random-snp*). These datasets were analyzed with STRUCTURE (same parameters as above) with *K* = 3 and *K* = 2, respectively.

### Genomic divergence

As a proxy to genomic divergence, net pairwise sequence divergence was computed using MEGA X (Kumar et al., 2018) between the ten *Alytes* species/lineages, based on the concatenated RAD tag sequences used in the final phylogenomic analyses of Ambu et al. (2023). The alignment comprises 278,267 bp (including 13,764 SNPs) and 45 samples (File S1).

### Bioacoustic differentiation

The advertisement call of midwife toads consists of single short high-pitched “whistling” notes, emitted from the ground in often unreachable spots (e.g., rock crevices). To study the diversity of calls, we combined our own call recordings with recordings from online media repositories.

Taxonomic assignments follow our phylogeographic results (see Results), and records of unclear origin or from areas of admixture were not included. Recordings were processed and analyzed in Raven Pro 1.6.1 (theCornellLab©) as in Ambu & Dufresnes (resubmitted) to identify unique individuals and measure four call parameters: the dominant frequency (DF) in kHz; the note duration (ND) in s; the rising time (RT) in s; and the pulse rate (PR) in s^-1^. The final bioacoustic dataset includes 721 notes from 153 individuals (1–9 notes per individual) representative of all taxa (File S2). For each parameter, individual averages were used in the statistical analyses.

To visualize call variation among taxa, we performed two Principal Component Analyses (PCA) on scaled data using the *R* package *FactoMineR* (Lê et al., 2008): one on the entire dataset, and the other restricted to the *A. obstetricans* complex. For comparative purposes, we computed a matrix of multivariate pairwise Euclidian distances between lineages, from which we built a neighbor-joining (NJ) tree (R package *stats*; R Core Team, 2023).

### Morphometric differentiation

We re-analyzed the raw measurements of museum specimens made by Martínez-Gil et al. (2022), reassigned to species and subspecies according to our phylogeographic results. Eight characters were measured (details in Martínez-Gil et al., 2022): snout-vent length (SVL); mouth length (ML); head width (HW); forelimb length (FLL); femur length (FML); tibia length (TBL); foot length (FTL), hindlimb length (HLL). Specimens of unclear origin or originating from areas of admixture were discarded. The final morphometric dataset consists of 211 individuals representing all taxa (File S3).

To compare body shape without the effect of body size, we applied allometric corrections (Chan & Grismer, 2022) with the *R* package *GroupStruct*, using the multispecies method to perform taxon-specific adjustments. As above, variation in body shape was assessed with two PCAs and by computing a matrix and NJ tree of multivariate pairwise Euclidian distances.

### Environmental differentiation

We compared the environmental conditions associated with each lineage’s occurrence based on 35 variables in raster format with 30 arc second resolution (File S4). These included elevation, 19 bioclimatic layers, 11 land cover layers, and 4 landscape layers. A total of 7,829 high precision (<1km) occurrences of *Alytes* were gathered from our own records, museum collections, published data and publicly available databases (Fig. 2; Ambu & Dufresnes, 2024). Records were assigned to taxa following our phylogeographic results and those from areas of admixture were subsequently excluded, resulting in an occurrence dataset of 7,086 localities, of which 5,031 were used after duplicate removal. As above, variations among lineages were assessed by two PCAs and a matrix/NJ tree of multivariate pairwise Euclidian distances. To assess the link between environmental differentiation and geography, we also computed pairwise Euclidian spatial distances among occurrence records.

**Fig. 2:**
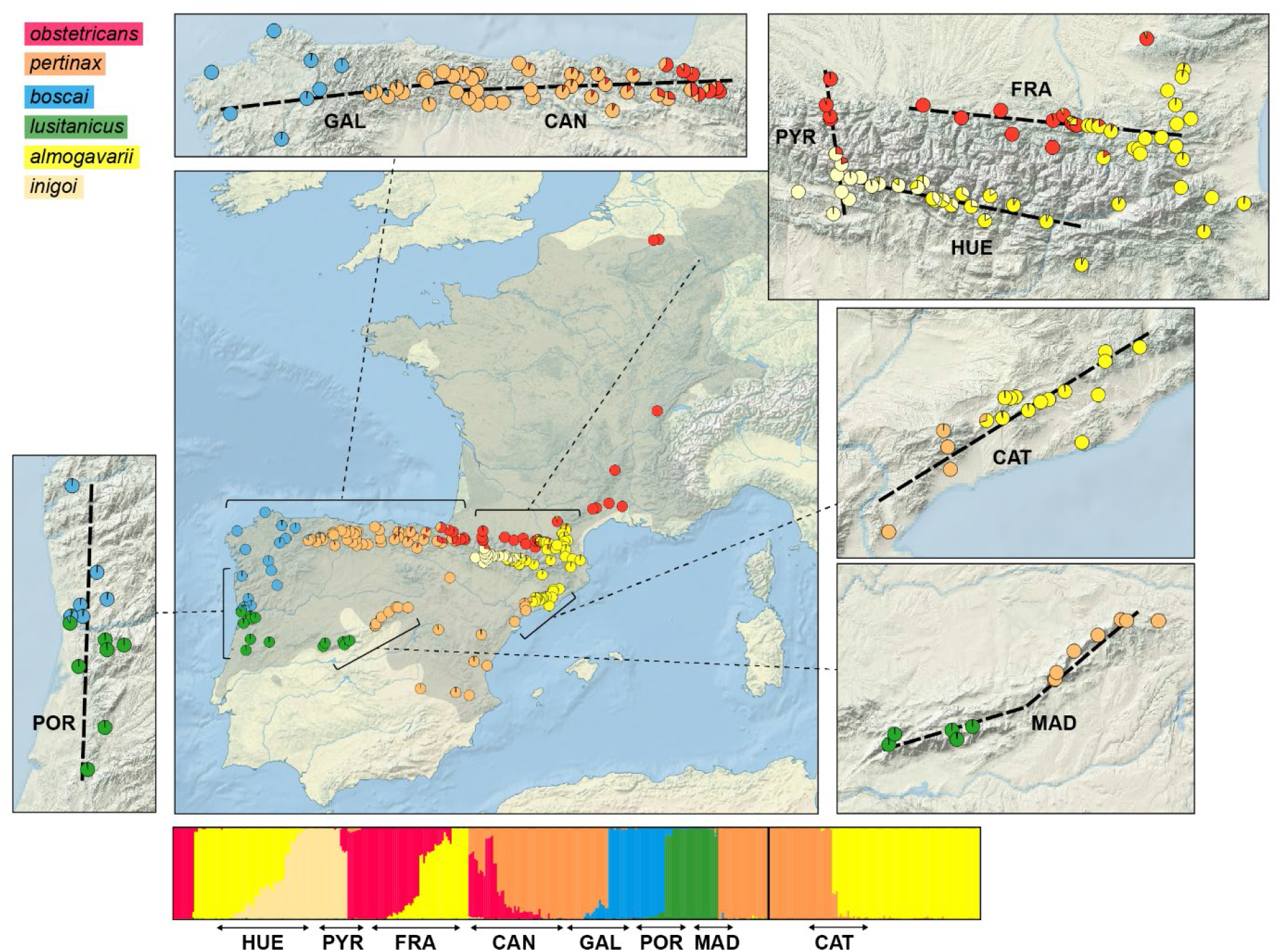
Genetic structure and admixture in the *A. obstetricans* complex. Barplots show individual ancestry as inferred by the clustering analysis of 5,111 genomic SNPs in six groups. Average ancestries are represented on the maps, with insets zooming in on parapatric ranges. Shaded grey: distribution of the *A. obstetricans* complex (see Fig. 1). Dashed lines show the transects used in the cline analyses (Fig. 3–4).

Ecological niche models were reconstructed with MaxEnt 3.4.4 (Phillips et al., 2006), using the WGS84 projection and a mask spanning from 31°N to 56°N and 11°W to 15°E that broadly covers the genus range. For each model, 10 replicates were run, retaining 30% of occurrences for random testing and 20,000 background points. The relative contributions of variables were estimated through a jackknife analysis, and maps of projected distributions were processed with the ClogLog output format (Phillips et al., 2017). Model calibration involved the evaluation of 372 candidate models produced with distinct regularization multipliers (0.5 to 6 at intervals of 0.5) and feature classes (resulting from all combinations of linear, quadratic, product, threshold, and hinge response types). The best model was selected considering statistical significance (partial ROC), predictive power (omission rates E = 5%), and complexity level (AICc), computed with the R package *kuenm* (Cobos et al., 2019), and its performance was assessed using the Area Under the Curve (AUC) and the True Skill Statistic (TSS) of the 10 percentile training omission threshold (Allouche et al., 2006).

Niche overlap between pairs of lineages was quantified using Schoener’s *D* distance (Schoener, 1968) computed in ENMTools (Warren et al., 2021).

### Comparisons across datasets

In the absence of environmental barriers to dispersal, cline width (*w*), which reflects the geographic extent of introgression between two parental genomes across their hybrid zone, offers an ad hoc proxy to compare the degree of RI between pairs of lineages with similar dispersal capacities (Dufresnes et al., 2021a, 2023). To explore which factors best determine RI between lineages of the *A. obstetricans* complex, we related their cline width *w* with their pairwise genomic, bioacoustic, morphological and environmental divergence as computed above. Two comparisons were performed: one on all eight hybrid zones, and one excluding two hybrid zones that appeared primarily mediated by geographic barriers (see Results). We further assessed how phenotypic and environmental divergence covaried with genomic divergence, within the *A. obstetricans* complex, and across the whole genus. Finally, we examined whether the contact zones of the *A. obstetricans* complex match environmental transitions for the lineages involved, by overlaying, for each transect, the genome average cline with the occurrence probabilities of the sampled populations predicted by the ecological niche models.

## Results

### Lineage distribution and hybrid zone analyses

Clustering analyses of 314 individuals (163 localities) genotyped at 5,111 SNPs in the *A. obstetricans* complex suggested up to six genetic groups (File S5), corresponding to the six lineages identified by Ambu et al. (2023) and attributed to six taxa: *A. a. almogavarii*, *A. a. inigoi*, *A. o. obstetricans*, *A. o. pertinax*, *A. o. boscai*, and *A. o. lusitanicus*. By combining our results with the ancestry estimates obtained from a previous ddRAD-seq study in NE-Spain (Dufresnes & Martínez-Solano, 2020), their distributions could be accurately inferred based on 425 individuals from 181 localities (Fig. 2).

The clustering analyses revealed the location of eight areas of presumed parapatry where individuals received intermediate ancestry estimates, suggestive of genetic admixture (Fig. 2). By fitting clines on the average genome ancestry (*Q_pop_*) along geographic transects, we obtained various cline widths *w*, characteristic of wide (>50km: CAN, HUE), intermediate (30–50km: GAL), steep (10–30km: CAT, FRA, PYR) and very steep (≤10km: POR, MAD) hybrid zones (Table 1, Fig. 3). Locus-specific clines confirmed these patterns (Table 1, Fig. 4), with their median *w* similar to the *Q_pop_ w* (File S6). The locus-specific analyses further documented high variation in cline parameters throughout the genome for the wide hybrid zones (CAN, HUE), while this variation was generally reduced for the steep hybrid zones, where most loci featured clines resembling the genome average (Fig. 4, File S6).

**Fig. 3:**
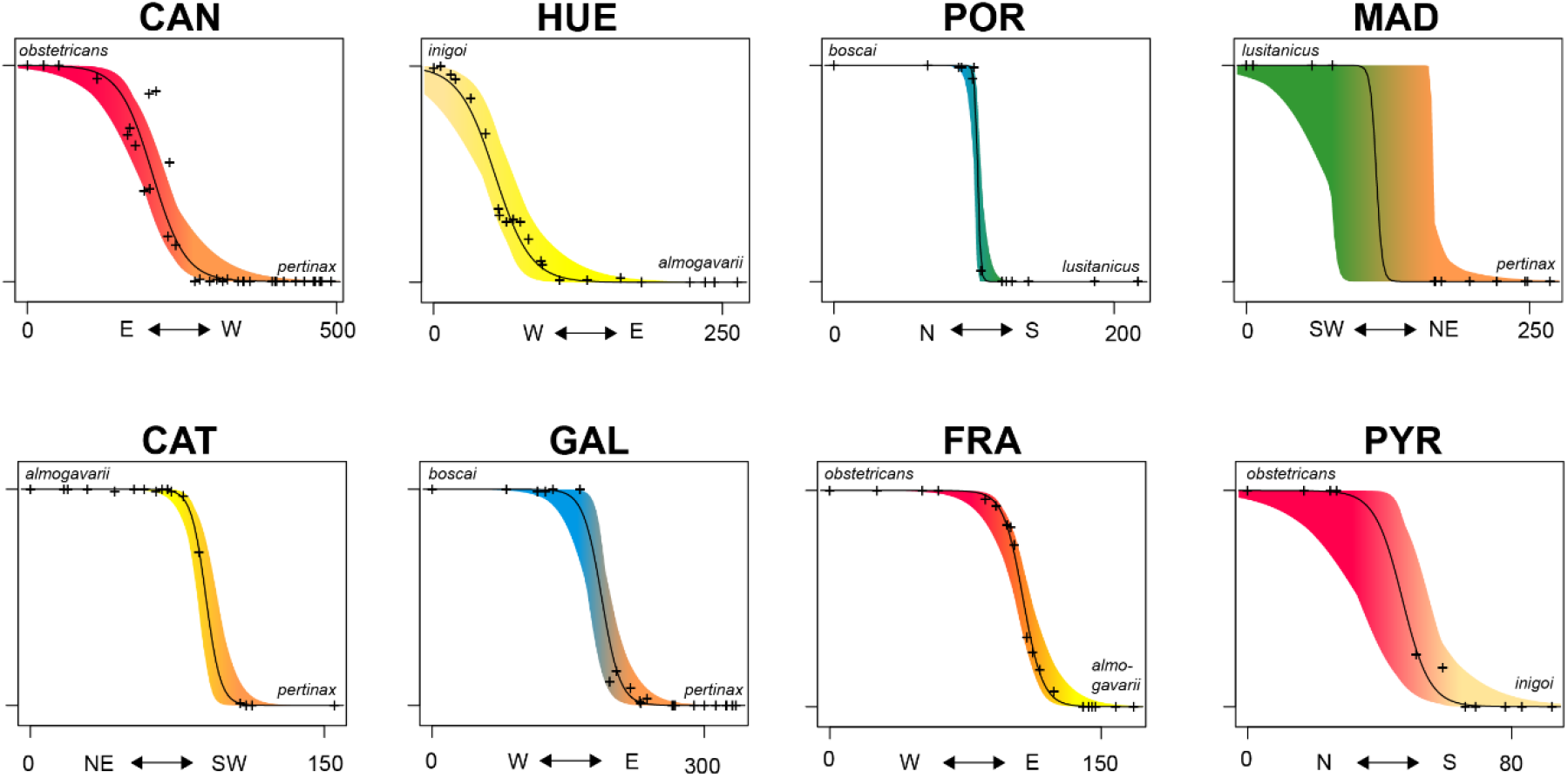
Geographic clines of genome-average ancestries fitted along transects for eight pairs of lineages in the *A. obstetricans* complex. Lines show the average clines, and colored areas show their 95% confidence intervals. Color gradients illustrate the lineages involved (see Fig. 1). Note that POR and MAD feature extrinsic barriers (see Fig. 2), which likely explains their steep transitions. Distances are in km.

**Fig. 4:**
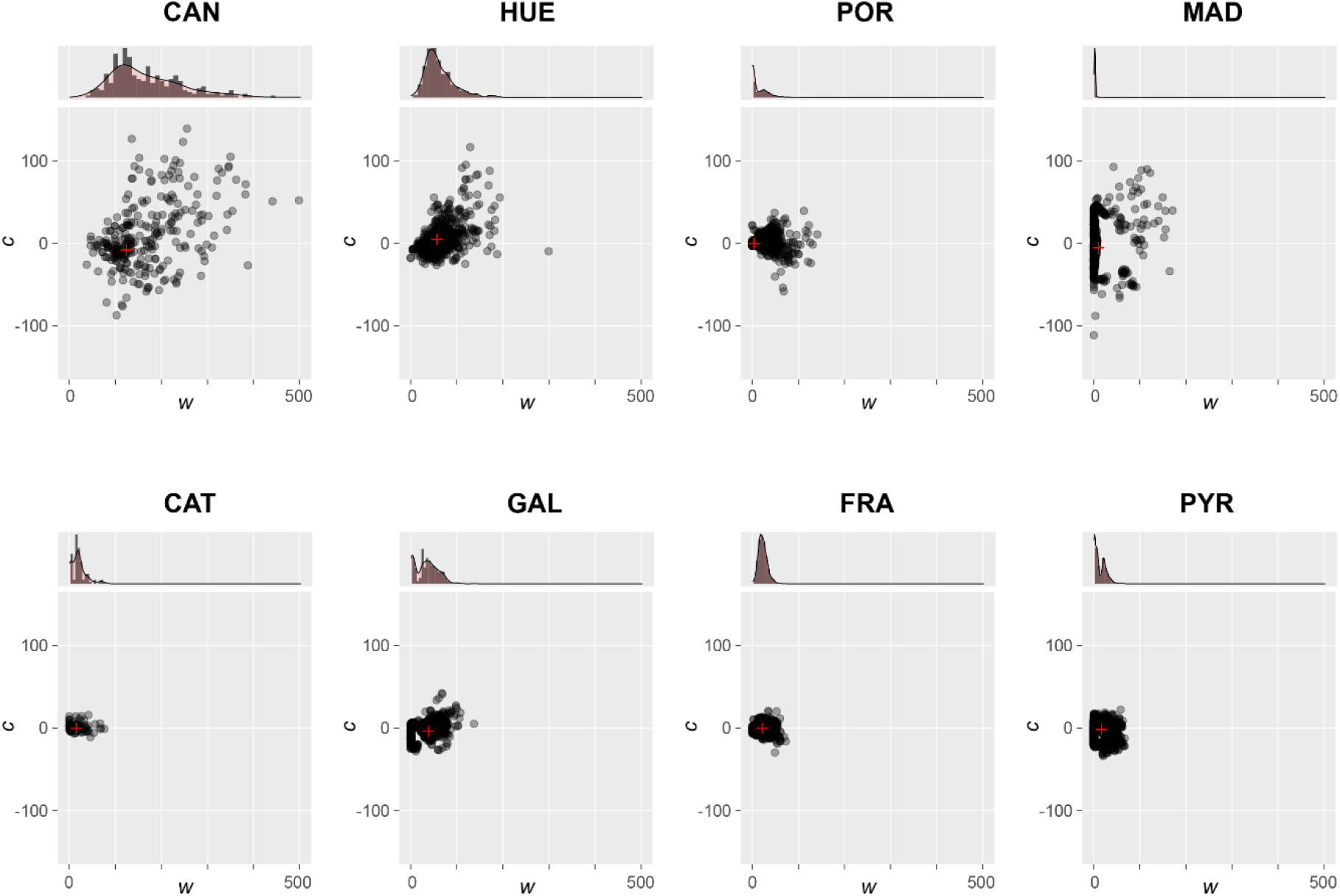
Cline widths (*w*) and center (*c*) computed separately for each diagnostic SNP along the eight transects analyzed in the *A. obstetricans* complex. Center estimates correspond to deviation from the genome average center. Red crosses show the mitochondrial clines. Histograms show estimate distributions (red overlay: density) for *w*. All units are km. Contact zones are arranged according to the phylogenetic distances of the involved lineages.

**Table 1:**
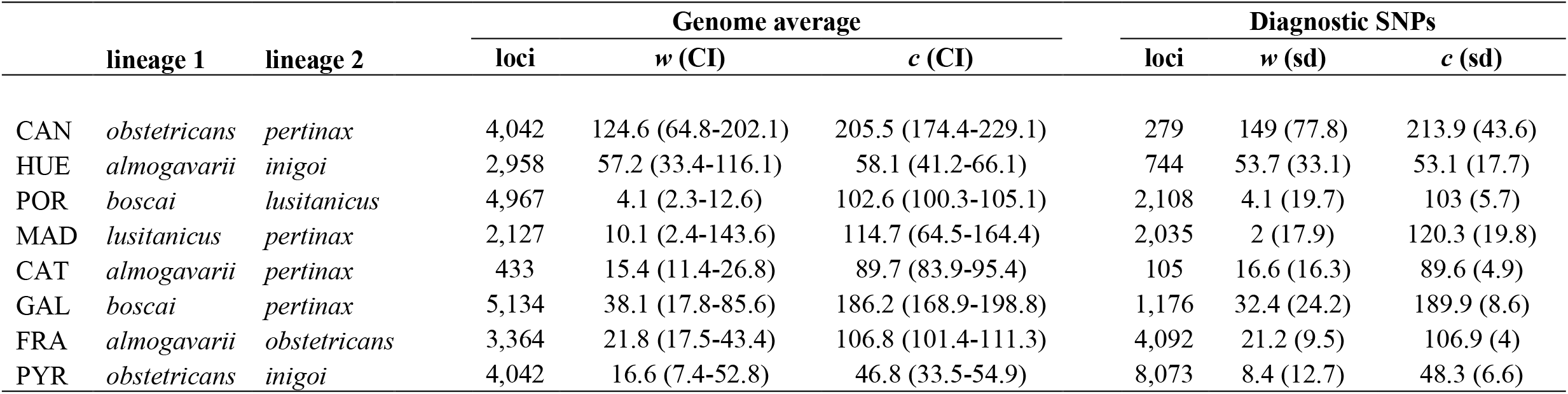
Summary statistics for the cline analyses in the eight hybrid zones surveyed; *w*: width, *c*: center; CI: 95% confidence interval; sd: standard deviation.

Two of the steepest contact zones, POR and MAD, featured a distinct pattern, namely very narrow clines at individual markers (*w* < 5km), yet with striking outliers, i.e., *w* > 100km (Fig. 4, File S6). A close inspection of their transition suggests current geographic isolation. In POR, the predicted center of the hybrid zone corresponds to the Douro River, the highest flow river of the Iberian Peninsula. In our transect area, the Douro River consists of a >250m wide brackish estuary (surrounded by the Porto urban area), which effectively separates *A. o. boscai* (north) and *A. o. lusitanicus* (south). In MAD, the predicted center of the hybrid zone falls within a ∼75km sampling gap between the Sierra de Gredos (inhabited by *A. o. lusitanicus*) and the Sierra de Guadarrama (inhabited by *A. o. pertinax*), which substantially reduces the interpretation of cline width estimates – the confidence interval of the genome average *w* accordingly spans over more than 100km (Table 1, Fig. 3). Importantly, this gap likely represents a real absence of *A. obstetricans*, since only *A. cisternasii* was reported from the area (Fig. 1). Therefore, while the faint traces of admixture (e.g., the few wide clines, Fig. 4) indicate that secondary contacts have probably existed, these two pairs of lineages are now effectively allopatric. In contrast, the other six transitions span areas where midwife toads are continuously distributed (Fig. 1) and with shared ancestries in parapatric populations suggestive of habitat connectivity (Fig. 2).

Finally, we did not detect recent admixture between the *A. obstetricans* complex and parapatric/sympatric midwife toad species, including where these are in close proximity (Fig. 5). Clustering analyses of *A. cisternasii*, *A. o. lusitanicus* and *A. o. pertinax* with *K* = 3, and of *A. o. pertinax* and *A. dickhilleni* with *K* = 2 distinguished these taxa and no sample showed mixed ancestry (Fig. 5).

**Fig. 5:**
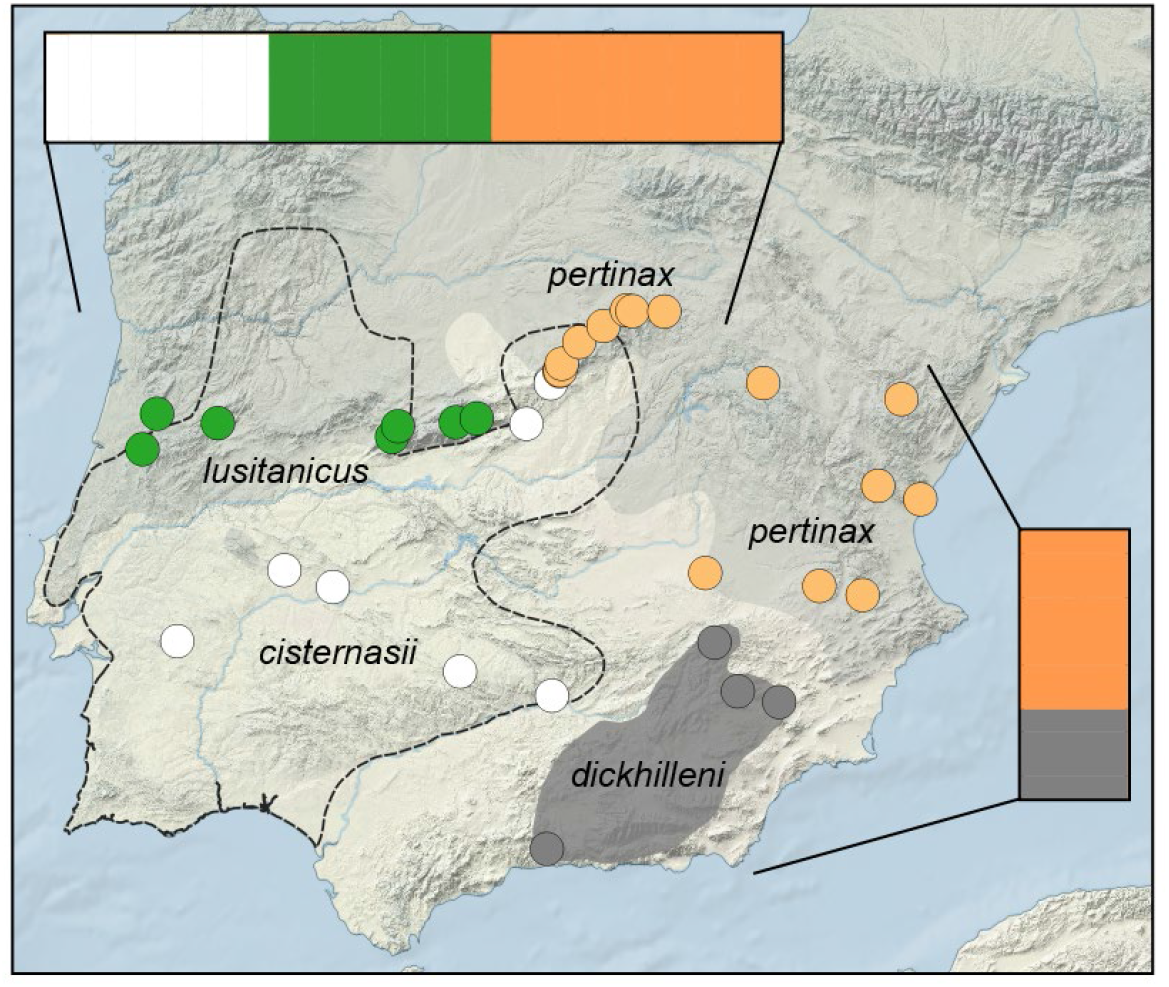
Admixture analyses between species from distinct clades / subgenera in their areas of sympatry / near-parapatry. The map shows sample locations and the barplots show individual ancestry from two separate analyses: (1) between *A. cisternasii* (subgenus *Ammoryctis*) and *A. obstetricans* from central (*pertinax*) and western Iberia (*lusitanicus*) (subgenus *Alytes*); (2) between *A. dickhilleni* (subgenus *Baleaphryne*) and *A. obstetricans* (*pertinax*) from southeastern Iberia (subgenus *Alytes*).

### Genomic divergence

Based on 278,267 bp (including 13,764 SNPs), net pairwise divergence spanned 1.4–9.1‰ across the whole genus, including 1.4‰ –4.1‰ within the *A. obstetricans* complex (subgenus *Alytes*), 3.7–5.7‰ among *A. maurus*, *A. muletensis*, *A. dickhilleni* (subgenus *Baleaphryne*), and >3.9‰ between *A. cisternasii* (subgenus *Ammoryctis*) and all other lineages (File S7).

### Bioacoustic differentiation

Call parameters, measured from four variables in 153 individuals, generally differed between the *A. obstetricans* complex and other species (Fig. 6A, File S8; with some overlap), except the Moroccan *A. maurus*. On the PCA based on the whole dataset, the Majorcan *A. muletensis* was partly distinguished by axis 1, which was influenced by variables DF and PR. The Iberian *A. cisternasii* and *A. dickhilleni* were partly distinguished by axis 2, which was influenced by variables RT and ND. All six lineages of the *A. obstetricans* complex share generally similar call characteristics and are grouped together in the NJ tree of multivariate Euclidian distances (Fig. 6A).

**Fig. 6:**
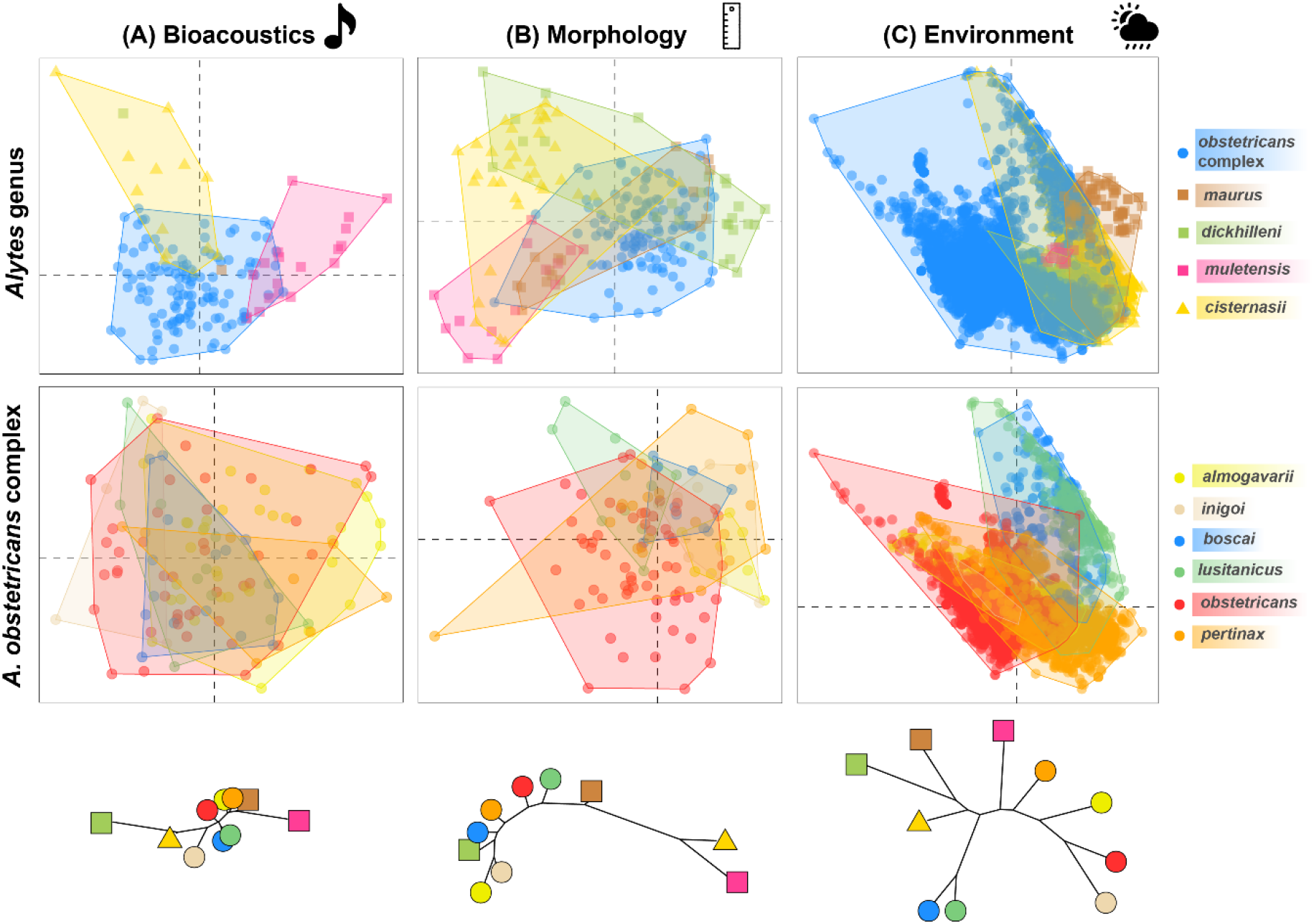
Phenotypic variation in *Alytes*. First dimensions of the PCAs built on bioacoustic (A), morphological (B), and environmental variables (C), distinguishing species from the three main clades (top panels) and the six lineages of the *A. obstetricans* complex (middle panels), and NJ trees of Euclidian distances (bottom). The dots on the PCAs represent individuals (A–B) or localities (C).

### Morphometric differentiation

Body shape, measured from seven variables (adjusted by body size) in 211 individuals, also suggested species-specific differences (Fig. 6B, File S9). On the PCA based on the whole dataset, axes 1 and 2 distinguished most individuals of *A. muletensis*, *A. cisternasii* and *A. dickhilleni* from the *A. obstetricans* complex and *A. maurus*, which both largely overlap. The PCA restricted to the *A*. *obstetricans* complex and the NJ tree of Euclidian distances also revealed noticeable patterns among sister lineages, including overlap and resemblance between *almogavarii* and *inigoi*, little overlap between *boscai* and *lusitanicus*, and high variability in the widespread *obstetricans* and *pertinax* (Fig. 6B).

### Environmental differentiation

The environmental conditions of 7,087 localities, summarized by 35 variables, were the most variable for the *A. obstetricans* complex, which largely overlap with other species except *A. maurus* (Fig. 6C). The PCAs broadly distinguished two climatic spaces, one corresponding to localities of *A. obstetricans* (*boscai* and *lusitanicus*) and *A. cisternasii* from W-Iberia, and the other corresponding to the rest of the ranges (Fig. 6C).

The ecological niche models built for each of the ten *Alytes* lineages overall match their current distribution (Fig. 7). The average AUC and TSS evaluations spanned 0.81–1.00 and 0.50-0.99, indicating high predictive power. Performance metrics for parameter settings are provided in File S4. Several bioclimatic (Bio2, Bio4, Bio6, Bio9, Bio11, Bio14, Bio15, Bio18, Bio19), landcover (barren land, global land cover, % of mixed forest, % of shrubs), and topographic (elevation and terrain ruggedness index) variables had relatively high contributions (˃10%) in all models. Models had low niche overlap overall according to Schoener’s *D* (*D* < 0.25, File S10), except for the three W-Iberian lineages *A. o. boscai*, *A. o. lusitanicus* and *A. cisternasii* (*D* = 0.35–0.47).

**Fig. 7:**
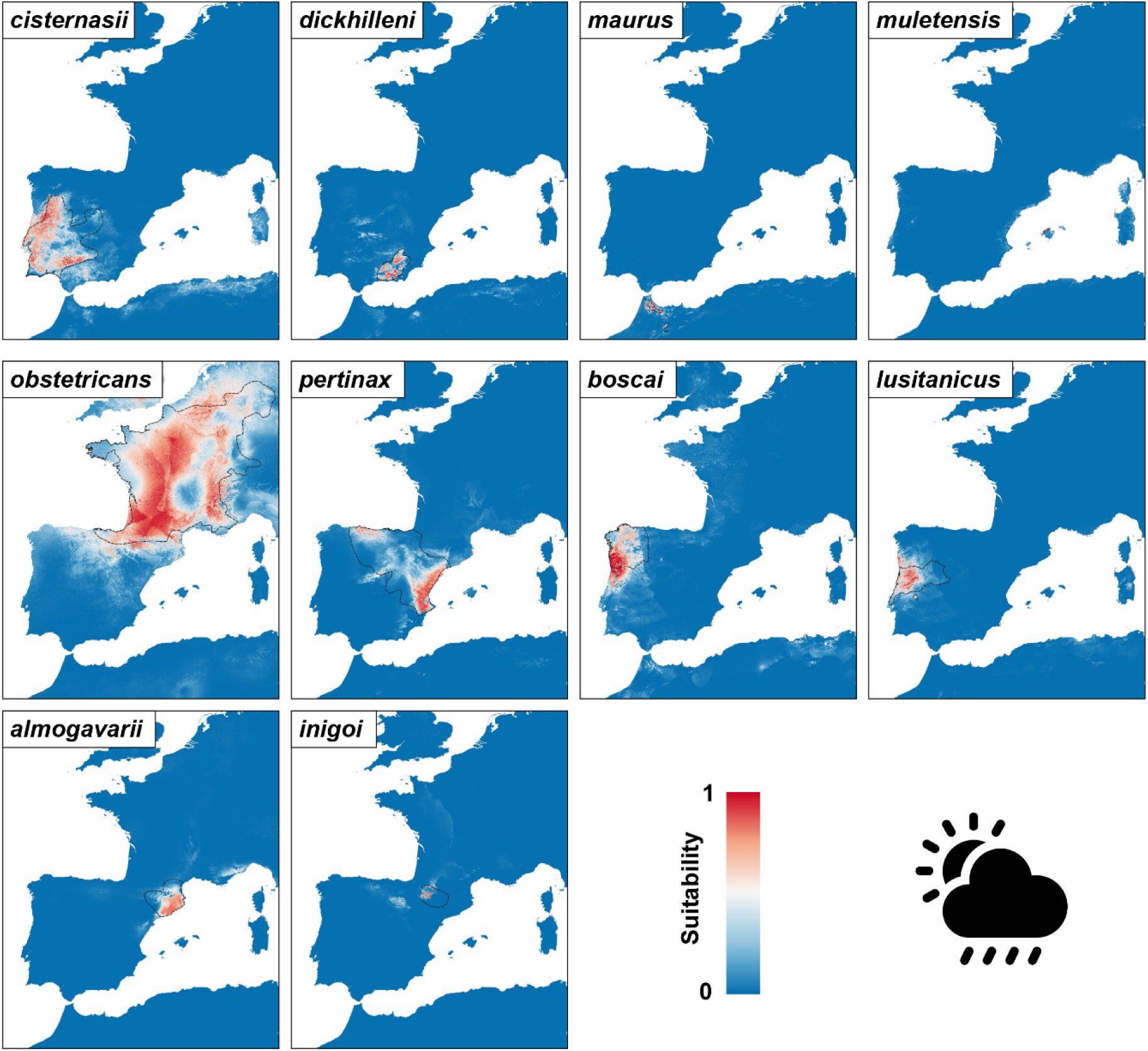
Projected distributions of each *Alytes* taxon based on climatic niche modelling. The same gradient of suitability is used for comparisons. Dashed lineages show current distributions (Fig. 1).

The pairwise Euclidian distances of environmental conditions (File S11) were not related to the niche dissimilarities computed from the reconstructed models (as 1-*D*) (File S12). However, they were significantly related to the spatial Euclidian distances (File S13), as expected given the strong spatial autocorrelation of environmental variables at regional scales (Journé et al., 2020).

### Comparisons across datasets

In the *A. obstetricans* complex, neither genomic, bioacoustic, morphological or environmental differentiation significantly predict cline width *w* when all eight pairs of lineages are considered (linear regression on log-transformed data; Table 2, Fig. 8). When excluding the two transitions that appear mediated by strong geographic barriers (POR and MAD), a negative relationship between genomic divergence and cline width becomes evident, with as much as 84% of variance explained (linear regression, *F* = 21.2, *P* = 0.01), while the other variables still show no effect (*P* > 0.20 in all cases, Table 2, Fig. 8).

**Fig. 8.**
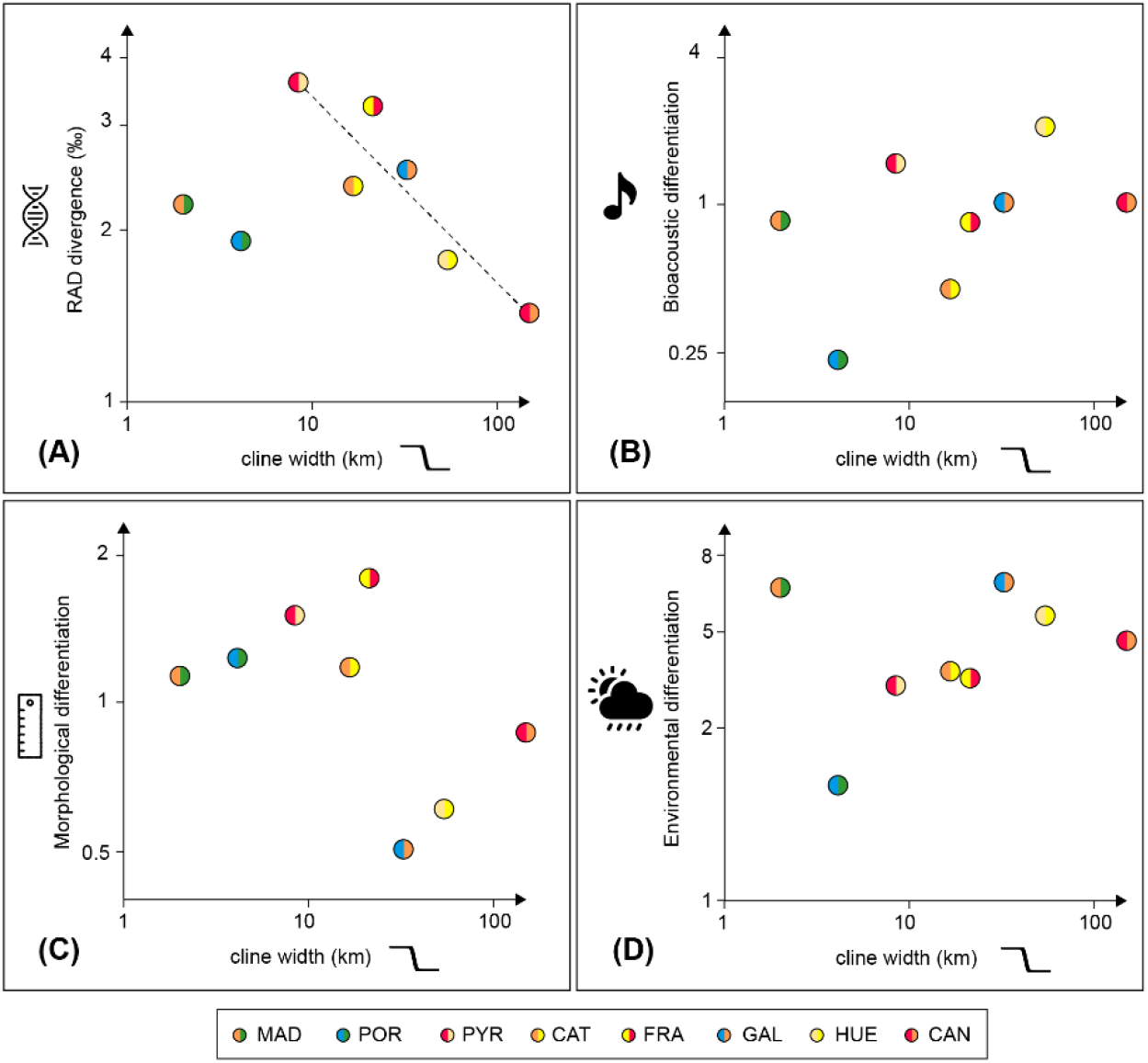
Relationship between cline width and genomic divergence (A), bioacoustic (B), morphological (C), and environmental differentiation (D) in the *A. obstetricans* complex. Axes are log-scaled. Only a single correlation was significant, namely between RAD divergence and cline width for the six lineage pairs currently in contact (*P* < 0.05; dashed line). Colors reflect the lineages involved in the corresponding contact zones.

**Table 2:** Statistical tests of the relationship between hybrid zone width (*w*) and genomic, bioacoustic, morphometric and environmental divergence (linear regressions; R^2^: proportion of variance explained; dof: degrees of freedom; F: F-statistic; *P* = p-value), and between the genomic divergence and the latter three (Mantel tests; *r*: correlation coefficient; *P*: p-value). Bold values indicate significance (*P* < 0.05).

| | cline width ($w$) | | | | | | | | genomic divergence | | | |
| --- | --- | --- | --- | --- | --- | --- | --- | --- | --- | --- | --- | --- |
|  | 8 hybrid zones |  |  |  | 6 hybrid zones |  |  |  | *A. obstetricans*<br>complex |  | *Alytes* genus |  |
| | $R^2$ | dof | F | $P$ | $R^2$ | dof | F | $P$ | $r$ | $P$ | $r$ | $P$ |
| genomic | 0.15 | 6 | 1.0 | 0.35 | **0.84** | **4** | **21.2** | **0.01** | - | - | - | - |
| bioacoustic | 0.20 | 6 | 1.5 | 0.26 | 0.04 | 4 | 0.1 | 0.72 | 0.11 | 0.35 | **0.47** | **0.04** |
| morphometric | 0.23 | 6 | 1.8 | 0.23 | 0.30 | 4 | 1.7 | 0.26 | 0.38 | 0.08 | 0.14 | 0.21 |
| environmental | 0.08 | 6 | 0.5 | 0.49 | 0.28 | 4 | 1.5 | 0.28 | 0.39 | 0.12 | **0.43** | **0.01** |

Pairwise genomic divergence does not co-vary with pairwise Euclidian distances at any phenotypic variable in the *A. obstetricans* complex (Mantel tests, Table 2, Fig. 9). At the level of the entire genus, however, bioacoustic and environmental differentiation significantly increase with pairwise genomic divergence (*P* < 0.05; Table 2, Fig. 9).

**Fig 9.**
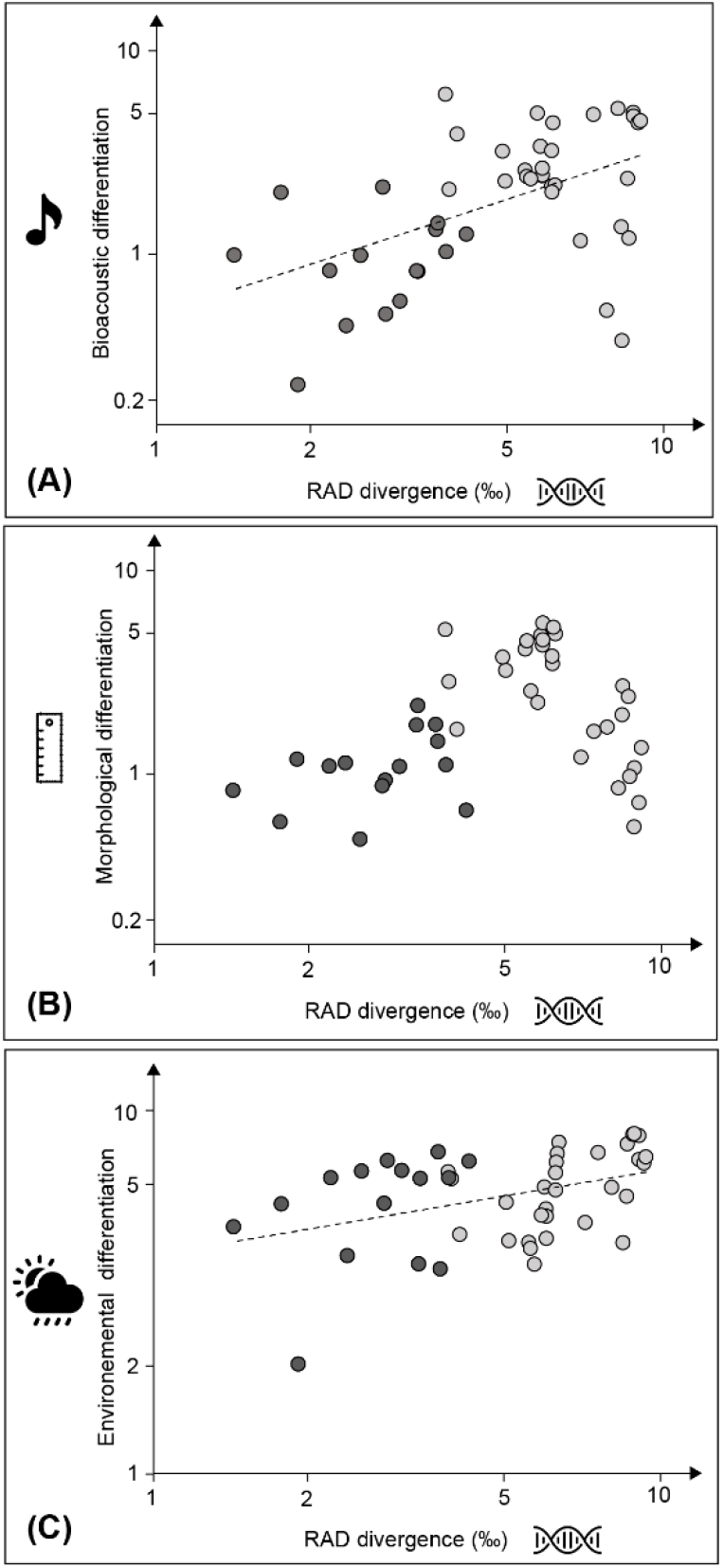
Relationship between genomic divergence and bioacoustic (A), morphological (B), and environmental differentiation (C). Significant links (*P* < 0.05) according to Mantel tests are represented by dash lines. Dark grey dots: pairwise comparisons within the *A. obstetricans* complex; light grey dots: other comparisons.

The contact zones of the *A. obstetricans* complex did not correspond to abrupt environmental transitions for the lineages involved (File S14), with the exception of *inigoi* in PYR. The occurrence probabilities obtained with the ecological niche models either remain similar on both sides of the hybrid zones, or progressively decrease along the transects from one lineage to the other, but without following the genetic transitions (File S14).

## Discussion

### Diverge first, differentiate later

Considering only the pairs of lineages that are currently in contact, the gradual decrease of gene flow with genomic rather than phenotypic divergence in the *A. obstetricans* complex suggests a primary role for genetic factors in initiating speciation. In the steepest hybrid zones (with *w* < 30km), which involve the most diverged lineages, some loci show no admixture at all and may represent barrier loci, i.e., loci linked to the genes causing reproductive barriers (see also Dufresnes & Martínez-Solano, 2020; Ambu & Dufresnes, resubmitted). In contrast, the whole genome seems to introgress freely in the widest hybrid zones (*w* > 50km), which accordingly involve the least diverged lineages. These results largely corroborate the mass of genes model for the buildup of RI (Dufresnes et al., 2021a), which predicts that nascent species become incompatible due to the accumulation of multiple small-effect mutations that additively reduce intrinsic hybrid fitness, such as DMIs (Orr, 1995; Orr & Turelli, 2001; Coyne & Orr, 2004). After a certain degree of divergence, DMIs are numerous enough to substantially affect hybrid viability and fertility (the “snowball effect”, Matute et al., 2010; Moyle & Nakazato, 2010), which eventually restricts gene flow and effectively prevents lineage fusion despite recurrent opportunities for introgressive hybridization.

Our analyses further suggests that the mass of genes model does not necessarily involve conspicuous ecological adaptations or behavioral signals that lineages would have potentially evolved independently during their time in allopatry, at least based on the traits considered here. The steepness of the hybrid zones is independent from the degree of morphological, bioacoustic and environmental differentiation of the lineages involved – which are accordingly not linked to their genetic divergence. Despite high phenotypic variability (see also Arntzen & García-París, 1995), the six lineages of the *A. o. obstetricans* complex remain broadly similar and no external diagnostic criteria could be established to discriminate them (Ambu et al., 2024). In parallel, the strong transgression of ecological niches, which reflect the climatically distinct regions occupied by the lineages (see also Reino et al., 2017; Rodríguez-Rodríguez et al., 2020), are irrespective of their divergence and do not presume of their fundamental climatic tolerance. In particular, the niches are not more different between reproductively (partly) incompatible than compatible lineages, and the phylogeographic transitions do not follow abrupt shifts in environmental suitability. The projected distributions illustrate that some steep hybrid zones are located in areas of generally suitable conditions (e.g., *obstetricans*/*inigoi*, *obstetricans*/*almogavarii*) while the widest hybrid zones were retrieved as climatically suboptimal (e.g., *obstetricans*/*pertinax*, *almogavarii*/*inigoi*) for at least one of the lineages involved.

If RI gradually evolves by the mass of genes model with little contribution by extrinsic factors, it is tempting to assume that the speciation in midwife toads is a neutral, long-term consequence of population isolation resulting from biogeographic processes. Nevertheless, whether the candidate barrier loci detected in the steepest hybrid zones are under selection in the parental species was not assessed, and reciprocally, other phenotypic traits that can potentially play an adaptive role in amphibian speciation may have been overlooked, such as coloration, olfaction, physiology, habitat use, or breeding phenology (Wollenberg Valero et al., 2019). For instance, color patterns were shown to greatly vary at the local scale in the *A. obstetricans* complex, perhaps to enhance camouflage (Polo-Cavia et al., 2016), and fixed differences have been suspected between some subspecies. This variation has not been investigated, however, and its importance for sexual selection in midwife toads is virtually unknown. It is further possible that local population differentiation at range margins, notably in habitat use and advertisement calls, may contribute some pre-mating isolation, notably through reinforcement (Vences & Wake, 2007; Wollenberg Valero et al., 2019). Reinforcement has been invoked to explain a reduction of the bioacoustic variation in the parapatric populations of *A. obstetricans* and *A. almogavarii* compared to their allopatric populations (Ambu & Dufresnes, resubmitted).

As exemplified by classical crossing experiments in anurans (Malone & Fontenot, 2008), as well as comparative studies of hybrid zones in vertebrates (e.g., Singhal & Moritz, 2013; Dufresnes et al., 2019a, 2021b; Pulido-Santacruz et al., 2020), the gradual relationship between the degree of RI and genetic divergence is probably a general rule across animals. Yet this relationship remains inherently difficult to establish when comparing datasets across different organisms, and where the proxy used to infer RI (here, the geographic extent of admixture summarized by *w*) is influenced by external factors. For instance, global meta-analyses mainly emphasized the effect of dispersal on *w* (McEntee et al., 2020), which is expected if both vagile and slow-moving species are included, in turn reducing the informativeness of hybrid zone inferences to quantify variations in the strength of RI. By comparing only organisms of broadly similar dispersal rates, the effect of divergence time on *w* becomes clearer (e.g., Dufresnes et al., 2021b). Here, the focus on a single species complex (where all lineages *a priori* share the same dispersal abilities), the exclusion of currently disconnected transitions (where *w* reflects geographic rather than reproductive barriers), and the estimation of genetic divergence based on the same set of genomic markers (rather than using indirect estimates like divergence time, or single gene genetic distances) might all have contributed to the neat relationship retrieved. Accordingly, genus-specific assessments in Australian skinks also reported remarkably strong correlations between lineage divergence and the amount of gene flow at range margins (Singhal & Moritz, 2013).

Although the grey zone of *Alytes* speciation appears mediated by genetic rather than phenotypic divergence, the latter does become substantial outside the grey zone, namely between deeply-diverged species from different subgenera, which are now completely isolated. Members of subgenus *Alytes*, *Ammoryctis* and *Baleaphryne* show no trace of hybridization despite local sympatry (*A. obstetricans*/*cisternasii*) or near-parapatry (*A. obstetricans*/*dickhilleni*), which parallels their greater genomic divergence, but also their more pronounced differences in shape, calls and environmental niches compared to the younger hybridizing lineages of the *A. obstetricans* complex. The only exception is *A. maurus* which was accordingly allocated to the subgenus *Alytes* before the molecular era (Martínez-Solano et al., 2004). In addition to raw genetic divergence, the species differences in body shape are potentially driven by adaptations to different habitats such as soil substrates (e.g., distinct dwelling behaviors, Arntzen & García-París, 1995), which could explain the absence of a linear relationship between molecular and morphological differentiation. In contrast, the gradual increase of bioacoustic differentiation emphasizes that anuran vocalizations can contain phylogenetic information at the macro-evolutionary scale (Robillard et al., 2006; Goicoechea et al., 2010). Finally, the significant link between genetic distances and niche differences probably reflects the geographic proximities and thus the climatic affinities of lineages found within each subgenus, namely the Mediterranean habits of *A. cisternasii* (*Ammoryctis*), the southern mountainous environments of *Baleaphryne* members (*A. maurus*, *A. dickhilleni* and *A. muletensis*), and the northern, globally more temperate climates experienced by the *A. obstetricans* complex (Rodríguez-Rodríguez et al., 2020; Donaire-Barroso et al., 2022; IUCN Red List 2024).

In turn, the higher phenotypic and ecological differentiation observed between the *Alytes* subgenera might be key to the establishment of sympatry. The co-existence of related species implies efficient pre-mating barriers to limit unfruitful hybridization and avoid wasting reproductive efforts, together with ecological differences to reduce niche overlap and escape competitive exclusion. If these attributes evolve gradually with genetic divergence, then sympatry should be the hallmark of deeply-diverged species that have first speciated in allopatry (Wollenberg Valero et al., 2019; Rasolonjatovo et al., 2020; Dufresnes et al., 2021b). Accordingly, here only the most diverged species, *A. cisternasii*, is sympatric with other midwife toads in a substantial part of its range, and its old divergence (≥13 My; Ambu et al., 2023) roughly corresponds to the divergence of other sympatric species of Palearctic anurans (e.g., 20 My in *Hyla* tree frogs, Dufresnes et al., 2020; 16 My in *Bufotes* green toads, Dufresnes et al., 2019; 13 My in *Pelophylax* water frogs; Dufresnes et al., 2024). Sympatry may then further boost phenotypic divergence, by a combination of divergent selection and selection for assortative mating (de Solan et al., 2022).

The ideas that genetic divergence first generates intrinsic isolation in allo-parapatry, and that external differences later evolve and allow sympatry, find roots in the early speciation literature. For instance, Haldane (1922) and Dobzhansky (1936) emphasized the important role of genetic factors in driving RI, while Mayr (1942)’s biological species consist of very divergent forms that are capable of spatial overlap (see also Mallet, 2008). In their seminal book, Coyne & Orr (2004) compromised with these aspects by considering that RI does not need to be complete to achieve speciation. Darwin himself viewed species as the long-term product of natural selection within populations (Darwin, 1859), with the adaptive radiation of Darwin’s finches – which he saw as “varieties” rather than species – being his most famous example. His postulates undoubtedly influenced the next 150 years of speciation studies and have led to the widespread assumption that adaptive factors are the likely source of much of Earth’s biodiversity (Ronco & Salzburger, 2021). Consequently, most of today’s naturalists see species as populations that are necessarily distinct ecologically, morphologically and/or behaviorally. While there is no doubt that local adaptation continuously shapes the genomic and phenotypic variation of nascent species (Butlin & Faria, 2024) and is behind the diversification of many vertebrates (e.g., birds, Cooney et al., 2017; fishes, Seehausen, 2006; but also amphibians and reptiles, Blackburn et al., 2013; Wollenberg Valero et al., 2019), its ubiquitous contribution to reproductive barriers in early stages of speciation is mitigated by three common empirical observations: (1) the existence of externally similar yet reproductively isolated species (“cryptic species”; Bickford et al., 2006) – implying that speciation does not always require phenotypic divergence, at least noticeable by the human eye; (2) the high phenotypic variation at key reproductive and ecological traits among conspecific populations (e.g., Dufresnes et al., 2022; Mulder et al., 2022) – implying that such adaptive innovations do not always lead to speciation; (3) the emergence of sympatry at the very end of the speciation continuum (e.g., Rasolonjatovo et al., 2020) – implying that pre-mating barriers appear after speciation is already complete. Moreover, adaptive radiation is supposed to be a non-gradual process (Ronco & Salzburger, 2021), which conflicts with the gradual nature of speciation retrieved in many vertebrate species groups (e.g., Singhal & Moritz, 2013; Pulido-Santacruz et al., 2020; Dufresnes et al., 2021b). Conversely, allopatric speciation might also be extremely (radiation-like) rapid without divergent selection, in case of repeated bottlenecks and small population sizes (Black et al. 2024). Disentangling under which circumstances speciation is an adaptive or a neutral outcome of population divergence thus deserves in-depth investigations, notably by comparing the mode and tempo of speciation between species groups that differ in their demographic history, evolutionary age, and potential for ecological diversification, e.g., with a high diversity of developmental trajectories (Bright et al., 2016) and ecological niches (Price et al., 2014; Ezard & Purvis, 2016).

### Species delimitation with and without hybrid zones

Patterns of admixture in hybrid zones are key to the taxonomic delimitation of phylogeographic lineages in the gray zone of speciation, at least under the biological species concept (Mayr, 1942), as they can inform on whether hybridizing lineages are likely to remain distinct over evolutionary times (Dufresnes et al., 2021b, 2023; Chambers et al., 2023). In this respect, narrow hybrid zones featuring strong barrier loci are typically maintained by selection against hybrids strong enough to prevent the lineages to fuse back (“speciation reversal”, Kearns et al., 2018). In European anurans, Dufresnes et al. (2021b) proposed two criteria to distinguish species and subspecies: (1) the genome average cline width *w*, noting that species lineages usually admix over less than 30km; (2) the heterogeneity of introgression across the genome, noting that species lineages usually feature loci with very steep clines (barrier loci). Applying these criteria necessitates proper transect sampling of the hybrid zones, and because *w* is determined by the balance between dispersal and selection against introgression (Barton & Gale, 1993), species with smaller dispersal distance than the average European anurans will exhibit steeper clines for the same strength of counterselection against hybridization.

The present findings emphasize the limits of the approach with irregular sampling and when effective barriers to dispersal occur in the middle of the hybrid zones. Without a critical interpretation, the very low average *w* in POR and MAD would suggest to oversplit *A. obstetricans* in three species (*A. boscai*, *A. lusitanicus*, and *A. obstetricans*/*pertinax*). However, in POR (*boscai*/*lusitanicus*), the Douro estuary is expected to significantly restrict gene flow, thus the low *w* does not necessarily indicate reproductive isolation. Accordingly, the river putatively mediates the lineage distribution in the newt *Triturus marmoratus* (Kazilas et al., 2024), as do other riverine systems in other Iberian amphibians like fire salamanders (Figueiredo-Vázquez et al., 2021). In MAD (*lusitanicus*/*pertinax*), the few wide clines suggest gene flow, and what is now a distributional gap may thus have been a hybrid zone. With the present sampling, the clines of loci that never introgressed beyond the edges of the gap are inferred with widths tending towards zero, thus mimicking barrier loci – even though these loci may have admixed within the gap. The issue is also characteristic of GAL (*boscai*/*pertinax*), where the two delimitation criteria mentioned above provide contradictory outcomes: the average extent of admixture (*w* ∼ 38km) is wide given the substantial number of very steep clines. This can be explained by the ∼35km sampling gap in the hybrid zone center of GAL, where clines as wide as 15km would theoretically fit even if assuming negligible admixture in the closest sampled populations (computed from the sigmoid formula; Barton & Gale, 1993). Hence, the current sampling of GAL does not truly allow to appreciate the presence of barrier loci and the delimitation of the two lineages it involves.

In the six other hybrid zones, the neat inverse relationship between genetic divergence and introgression indirectly implies that regional landscapes are generally less important than post-zygotic incompatibilities in mediating gene flow. That said, we cannot rule out that the topographic and hydrographic complexity of the study areas affected contemporary dispersal at local transitions, as seen in e.g., *Salamandra* (Velo-Antón et al., 2021). For instance, the hybrid zone centers of FRA (*obstetricans*/*almogavarii*) and PYR (*obstetricans*/*inigoi*) broadly match the Ariège river and the Portalet mountain pass, respectively – but both are eventually overcome by migrants, which is not surprising given the ecological, notably the altitudinal tolerance of the species. Comparatively large rivers and high mountain ranges are found elsewhere across the vast range of the *A. obstetricans* complex, where they did not substantially disconnect hybridizing lineages (e.g., CAN, *pertinax*/*obstetricans*).

Considering these points, here we can delimit species and subspecies for the six lineages of the *A. obstetricans* complex in taxonomic accounts. The ancestry estimates based on hundreds/thousands of SNPs further provide a fine-scale picture of their respective distribution ranges – which previously remained unclear based on mitochondrial and microsatellite markers (Gonçalves et al., 2015; Maia-Carvalho et al, 2018). *Alytes obstetricans* and *A. almogavarii* always feature narrow hybrid zones with putative barrier loci, namely between *almogavarii* and *pertinax* (CAT; Dufresnes & Martínez-Solano, 2020), *almogavarii* and *obstetricans* (FRA; Ambu & Dufresnes, resubmitted), and *obstetricans* and *inigoi* (PYR; this study). Hybrid zones involving the same incipient species may sometimes differ in their extent of admixture depending on intrinsic factors, such as the intraspecific lineages involved, as well as extrinsic factors, such as the age of the contact zone (Dufresnes et al., 2020a), dispersal opportunities (McEntee et al., 2020) or demographic dynamics (van Riemsdijk et al., 2023). Narrow transitions across several replicate transects are thus the hallmark of stable reproductive barriers (e.g., Dufresnes et al., 2021a). Therefore, our study corroborates the species status of *A. almogavarii*, as currently acknowledged (Speybroeck et al., 2020) although not widely accepted (e.g., Lucati et al., 2022). Reciprocally, the wide hybrid zones between the shallowest lineages confirm the subspecies status of *A. a. inigoi* and *A. o. pertinax*. Finally, the sister taxa *boscai* and *lusitanicus* are more complicated to rank. The intermediate patterns of admixture of *boscai* and *pertinax* in GAL emphasize the difficulty to classify lineages that fall in the middle of the gray zone of speciation. We here continue to treat them as subspecies of *A. obstetricans*, given that the genome average cline of the *boscai*/*pertinax* hybrid zone is overall slightly wider (∼38km) than the 30km threshold, and that the few wide clines in the *lusitanicus*/*pertinax* transect suggest that the former hybrid zone may have been potentially loose. These assessments may be revised in the future pending new hybrid zone analyses based on additional sampling.

When hybrid zones do not exist (or cannot be sampled), genetic or phenotypic divergence may help to rank evolutionary lineages in groups where reproductive isolation correlates with such divergence, affording species status to lineages that are as divergent as valid species in the same groups (Schweizer et al., 2023). Because of their comparability and universality in evolutionary studies, divergence times, that can to some extent be approximated by the percentage of divergence at mitochondrial genes, can be used as a proxy to define operational thresholds for species delimitation (Dufresnes et al., 2021b; Dufresnes & Litvinchuk, 2022). Here, the hybrid-zone validated species *A. obstetricans* and *A. almogavarii* supposedly diverged ∼3.9 Mya, while the hybrid-zone validated subspecies *A. a. inigoi* and *A. o. pertinax* originated <2.5 Mya (Ambu et al., 2023). This temporal window can thus assist the ranking of older or younger *Alytes* lineages left to discover as new species or subspecies, respectively. Because with sufficient divergence, any lineage may eventually become distinguishable externally, candidate species may still be identified based on substantial morphological and behavioral differentiation (e.g., Dubois, 2023). However, if speciation is primarily a matter of genomic divergence, comprehensive molecular analyses alone should suffice to justify species splitting or lumping. Rather than directly contributing to species delimitation decisions in integrative taxonomy (Padial et al., 2010), phenotypic analyses shall be relevant for finding diagnostic criteria to substantiate the description of new species and subspecies in compliance with nomenclatural rules (Braby et al., 2024).

Finally, it should be noted that species ages obtained from time-calibrated phylogenies and divergence estimates obtained from clonal sequences (e.g., mtDNA) may not be adequate to rank taxa of reticulate origin, since these taxa do not only result from tree-like evolutionary processes. Raw genomic divergence, which should covary with the average number of mutations involved in RI under the mass-of-genes model, might thus be more relevant, especially when hybrid taxa are included in the comparison – here the subspecies *A. o. pertinax* exhibits ancestry from both *A. o. obstetricans* and *A. almogavarii* (Ambu et al., 2023). In this respect, we note the similar percentages of sequence divergence at ddRAD-seq loci in our study compared to previous analyses on other anuran genera (Dufresnes et al., 2021b), where ∼3‰ always corresponds to a mid-point threshold between species and subspecies. Such comparability is not necessarily expected given that these analyses rely on different sets of markers, which polymorphism depends on methodological aspects such as bioinformatic stringency and sample scheme. Future investigations should explore the universality of divergence thresholds applicable across genomic datasets and species groups, in the framework of integrating next-generation sequencing approaches in taxonomy.

## Supplementary Material

File S1: List of the samples used in the genetic analyses; File S2: Bioacoustic data; File S3: Morphological data; File S4: Variables used in the climatic models and their contributions; File S5: Clustering analyses in *A. obstetricans* with K = 2–6; File S6: Relationship between genome average and individual cline width estimates; File S7: Pairwise genomic divergence; File S8: Pairwise bioacoustic differentiation; File S9: Pairwise morphological differentiation; File S10: Niche overlap between models; File S11: Pairwise environmental differentiation; File S12: Relationship between environmental differentiation and niche overlap; File S13: Relationship between environmental differentiation and spatial distances; File S14: Occurrence probabilities along the contact zone transects.

## Data Availability Statement

raw ddRAD sequence reads were uploaded on NCBI SRA under BioProject PRJNA949685; SNP, allele frequency, occurrence, and environmental datasets are available on Zenodo (Ambu & Dufresnes, 2024). Bioacoustic and morphological datasets are available as supplementary material (Files S2–S3).

## Acknowledgements

Sampling was conducted under collecting permits issued by DREAL Occitanie (DREAL-OCC-2023-s-05), the Community of Madrid and Sierra de Guadarrama National Park (10/184709.9/22, 10/035516.9/22, 10/370877.9/22), the Picos de Europa National Park (CO/09/05/2008 and CO/09/0571/2009), the governments of Cantabria (CAP/2021/038), Castilla y León (EP/CYL/389/2007, EP/LE/428/2010, EP/P/428/2010, EP/CYL/31/2010, EP/P/426/2010 and AUES/CyL/222/2022), Vizcaya (AM-16-2022), Álava (EXP-22-13-18), Guipúzcoa (2022-FAUNA- 206225527), Navarra (76E/2022), Aragón (INAGA/500201/24/2022/02013), Cataluña (SF/0042/22), the Xunta de Galicia (EB-016/2018), and the Principality of Asturias (2010/000371) and PA/356/2022/2455), as well as authorizations for handling with wildlife issued by the Community of Madrid (10/142121.9/20 to CCD) and Andalucía (EXP-000117 to IMS). We further warmly thank M. Vences (TU-Braunschweig) for laboratory access and support.

