## Supplementary material for "Genomic, phenotypic and environmental correlates of speciation in the midwife toads (*Alytes*)": SI

Ambu et al.

### Table of content


| | |
|---|---|
| **File S1:** List of the samples used in the genetic analyses | 2 |
| **File S2:** Bioacoustic data | 14 |
| **File S3:** Morphological data | 19 |
| **File S4:** Variables used in the climatic models and their contributions | 26 |
| **File S5:** Clustering analyses in the *A. obstetricans* complex with K = 2–6 | 28 |
| **File S6:** Relationship between genome average and individual cline width estimates | 29 |
| **File S7:** Pairwise genomic divergence | 30 |
| **File S8:** Pairwise bioacoustic differentiation | 31 |
| **File S9:** Pairwise morphological differentiation | 32 |
| **File S10:** Niche overlap between models | 33 |
| **File S11:** Pairwise environmental differentiation | 34 |
| **File S12:** Relationship between environmental differentiation and niche overlap | 35 |
| **File S13:** Relationship between environmental differentiation and spatial distances | 36 |
| **File S14:** Occurrence probabilities along the contact zone transects | 37 |

**File S1**: List of samples and inclusion in the contact zone analyses as follows. s: selected for the ancestry analyses; r: used as parental reference to find lineage diagnostic SNPs; t: used in cline analyses along transects. O-C: *A. obstetricans*/*cisternasii* analysis; O-D: *A. obstetricans*/*dickhilleni* analysis. BEV and BEV.T samples are housed in the collection of the Biogeography and Ecology team at the CEFE lab in Montpellier (EPHE, CNRS).

| Loc. | Sample | Y | X | Taxon | Source | GAL | POR | CAN | FRA | MAD | PYR | HUE | O-C | O-D |
|---|---|---|---|---|---|---|---|---|---|---|---|---|---|---|
| 1 | R262 | 50.8520 | 5.9720 | obstetricans | 1 | | | sr | sr | | sr | | | |
| 2 | R213 | 50.8380 | 5.7750 | obstetricans | 1 | | | sr | sr | | sr | | | |
| 3 | JEUR01 | 46.3700 | 5.7159 | obstetricans | 2 | | | sr | sr | | sr | | | |
| 4 | BEV.13848 | 44.8170 | 4.1780 | obstetricans | 1 | | | sr | sr | | sr | | | |
| 5 | BEV.14198 | 43.8712 | 4.4004 | obstetricans | 1 | | | sr | sr | | sr | | | |
| 6 | BEV.14199 | 43.9557 | 3.9451 | obstetricans | 1 | | | sr | sr | | sr | | | |
| 7 | LAR01 | 43.8326 | 3.5048 | obstetricans | 1 | | | sr | sr | | sr | | | |
| 7 | LAR03 | 43.8326 | 3.5048 | obstetricans | 1 | | | sr | sr | | sr | | | |
| 8 | BEV.T3119 | 43.8090 | 3.3789 | obstetricans | 1 | | | sr | sr | | sr | | | |
| 8 | BEV.T3120 | 43.8090 | 3.3789 | obstetricans | 1 | | | sr | sr | | sr | | | |
| 9 | DOUR01 | 43.4770 | 2.1475 | admixed | 2 | | | | | | | | | |
| 10 | VGLY10 | 43.2838 | 2.4430 | almogavarii | 2 | | | | s | | | | | |
| 11 | BOUG02 | 43.2407 | 2.4306 | almogavarii | 2 | | | | s | | | | | |
| 12 | COUF01 | 43.1574 | 2.3119 | almogavarii | 2 | | | | sr | | | sr | | |
| 12 | COUF02 | 43.1574 | 2.3119 | almogavarii | 2 | | | | sr | | | sr | | |
| 13 | GREF12 | 43.0688 | 2.3743 | almogavarii | 2 | | | | sr | | | sr | | |
| 13 | GREF13 | 43.0688 | 2.3743 | almogavarii | 2 | | | | sr | | | sr | | |
| 14 | SALZ01 | 42.9834 | 2.4970 | almogavarii | 2 | | | | sr | | | sr | | |
| 14 | SALZ02 | 42.9834 | 2.4970 | almogavarii | 2 | | | | sr | | | sr | | |
| 15 | ARQ04 | 42.9524 | 2.3836 | almogavarii | 2 | | | | sr | | | sr | | |
| 16 | JUL01 | 42.8691 | 2.2570 | almogavarii | 2 | | | | srt | | | sr | | |
| 16 | JUL04 | 42.8691 | 2.2570 | almogavarii | 2 | | | | srt | | | sr | | |
| 16 | JUL08 | 42.8691 | 2.2570 | almogavarii | 2 | | | | srt | | | sr | | |
| 17 | FEN02 | 42.8146 | 2.3735 | almogavarii | 2 | | | | srt | | | sr | | |

| | | | | | | | |
|---|---|---|---|---|---|---|---|
| 17 | FEN03 | 42.8146 | 2.3735 | almogavarii | 2 | srt | sr |
| 18 | SOU01 | 42.7331 | 2.4265 | almogavarii | 2 | sr | sr |
| 18 | SOU02 | 42.7331 | 2.4265 | almogavarii | 2 | sr | sr |
| 19 | FI01 | 42.5609 | 2.4097 | almogavarii | 2 | sr | srt |
| 19 | FI02 | 42.5609 | 2.4097 | almogavarii | 2 | sr | srt |
| 20 | ALM347 | 42.4986 | 2.6571 | almogavarii | 1 | sr | srt |
| 21 | MNCN81015 | 42.4624 | 2.9150 | almogavarii | 1 | sr | srt |
| 22 | ALM293 | 42.2885 | 2.5892 | almogavarii | 1 | sr | srt |
| 23 | TJ01 | 42.4590 | 1.9058 | admixed | 1 | | st |
| 24 | ALM408 | 42.0859 | 1.6079 | admixed | 1 | | st |
| 25 | MNCN81003 | 42.3517 | 1.3288 | admixed | 3 | | st |
| 26 | MNCN81001 | 42.4522 | 1.0645 | admixed | 1 | | st |
| 26 | MNCN81001b | 42.4522 | 1.0645 | admixed | 3 | | st |
| 26 | MNCN81002 | 42.4522 | 1.0645 | admixed | 3 | | st |
| 27 | MNCN80999 | 42.3565 | 0.8340 | admixed | 3 | | st |
| 27 | MNCN81000 | 42.3565 | 0.8340 | admixed | 3 | | st |
| 28 | RIB01 | 42.5093 | 0.8768 | admixed | 3 | | st |
| 28 | RIB02 | 42.5093 | 0.8768 | admixed | 3 | | st |
| 28 | RIB03 | 42.5093 | 0.8768 | admixed | 3 | | st |
| 29 | SARRO01 | 42.4437 | 0.7241 | admixed | 3 | | st |
| 29 | SARRO02 | 42.4437 | 0.7241 | admixed | 3 | | st |
| 29 | SARRO03 | 42.4437 | 0.7241 | admixed | 3 | | st |
| 30 | MNCN80998 | 42.5221 | 0.6520 | admixed | 1 | | st |
| 31 | ABE01 | 42.4516 | 0.5647 | admixed | 3 | | st |
| 31 | ABE02 | 42.4516 | 0.5647 | admixed | 3 | | st |
| 31 | ABE03 | 42.4516 | 0.5647 | admixed | 3 | | st |
| 32 | GAB01 | 42.4842 | 0.4955 | admixed | 3 | | st |
| 32 | GAB02 | 42.4842 | 0.4955 | admixed | 3 | | st |
| 32 | GAB03 | 42.4842 | 0.4955 | admixed | 3 | | st |
| 33 | SEI01 | 42.4804 | 0.4294 | admixed | 3 | | st |
| 33 | SEI02 | 42.4804 | 0.4294 | admixed | 3 | | st |

| | | | | | | | |
|---|---|---|---|---|---|---|---|
| 34 | BARB01 | 42.4928 | 0.4139 | admixed | 3 | | st |
| 34 | BARB02 | 42.4928 | 0.4139 | admixed | 3 | | st |
| 35 | GIT01 | 42.5911 | 0.3366 | admixed | 3 | | |
| 35 | GIT02 | 42.5911 | 0.3366 | admixed | 3 | | |
| 36 | SARA01 | 42.5607 | 0.2914 | admixed | 3 | | st |
| 36 | SARA02 | 42.5607 | 0.2914 | admixed | 3 | | st |
| 36 | SARA03 | 42.5607 | 0.2914 | admixed | 3 | | st |
| 37 | ESCU01 | 42.5730 | 0.1392 | admixed | 3 | | st |
| 37 | ESCU02 | 42.5730 | 0.1392 | admixed | 3 | | st |
| 37 | INIG01/MNCN50762 | 42.5730 | 0.1392 | admixed | 1 | | st |
| 38 | BUIS01 | 42.5818 | -0.0175 | admixed | 3 | | st |
| 38 | BUIS03 | 42.5818 | -0.0175 | admixed | 3 | | st |
| 38 | BUIS04 | 42.5818 | -0.0175 | admixed | 3 | | st |
| 39 | MNCN81034 | 42.5688 | -0.0772 | inigoi | 1 | | st |
| 40 | SORR01 | 42.6237 | -0.1683 | inigoi | 3 | sr | st |
| 40 | SORR02 | 42.6237 | -0.1683 | inigoi | 3 | sr | st |
| 41 | ABJ01 | 42.3988 | -0.3850 | inigoi | 3 | srt | sr |
| 41 | ABJ02 | 42.3988 | -0.3850 | inigoi | 3 | srt | sr |
| 41 | ABJ03 | 42.3988 | -0.3850 | inigoi | 3 | srt | sr |
| 41 | MNCN81004 | 42.3988 | -0.3850 | inigoi | 1 | srt | sr |
| 42 | MNCN81008 | 42.5335 | -0.6690 | inigoi | 1 | sr | sr |
| 43 | SOBA01 | 42.4879 | -0.2601 | inigoi | 3 | srt | sr |
| 43 | SOBA06/MNCN50838 | 42.4879 | -0.2601 | inigoi | 3 | srt | sr |
| 43 | SOBA06b/MNCN50838 | 42.4879 | -0.2601 | inigoi | 3 | srt | sr |
| 44 | JAV03 | 42.5345 | -0.3204 | inigoi | 3 | st | sr |
| 44 | JAV04 | 42.5345 | -0.3204 | inigoi | 3 | st | sr |
| 44 | JAV05 | 42.5345 | -0.3204 | inigoi | 3 | st | sr |
| 45 | YES01 | 42.6175 | -0.2483 | inigoi | 3 | st | srt |
| 45 | YES02 | 42.6175 | -0.2483 | inigoi | 3 | st | srt |
| 46 | BDS01 | 42.6390 | -0.3574 | inigoi | 3 | st | |
| 46 | BDS02 | 42.6390 | -0.3574 | inigoi | 3 | st | |

| | | | | | | | | |
|---|---|---|---|---|---|---|---|---|
| 46 | BDS06 | 42.6390 | -0.3574 | inigoi | 3 | | | st |
| 46 | BDS11 | 42.6390 | -0.3574 | inigoi | 3 | | | st |
| 47 | GORG01 | 42.7047 | -0.3214 | admixed | 3 | | | st |
| 48 | FORM04 | 42.7681 | -0.3664 | admixed | 3 | | | st |
| 48 | FORM05 | 42.7681 | -0.3664 | admixed | 3 | | | st |
| 48 | FORM06 | 42.7681 | -0.3664 | admixed | 3 | | | st |
| 49 | ASS01 | 42.9827 | -0.4103 | obstetricans | 3 | | | st |
| 50 | LOUV01 | 42.9996 | -0.4137 | obstetricans | 3 | | | st |
| 51 | BENOU04 | 43.0690 | -0.4445 | obstetricans | 3 | | | st |
| 51 | BENOU05 | 43.0690 | -0.4445 | obstetricans | 3 | | | st |
| 52 | GAN01 | 43.2284 | -0.4111 | obstetricans | 1 | st | | srt |
| 52 | GAN22_01 | 43.2284 | -0.4111 | obstetricans | 3 | st | | srt |
| 52 | GAN22_02 | 43.2284 | -0.4111 | obstetricans | 3 | st | | srt |
| 53 | AVE01 | 43.0665 | 0.3378 | obstetricans | 2 | srt | srt | sr |
| 53 | AVE06 | 43.0665 | 0.3378 | obstetricans | 2 | srt | srt | sr |
| 53 | AVE07 | 43.0665 | 0.3378 | obstetricans | 2 | srt | srt | sr |
| 54 | GAL01 | 42.9897 | 0.6472 | obstetricans | 2 | srt | srt | sr |
| 54 | GAL04 | 42.9897 | 0.6472 | obstetricans | 2 | srt | srt | sr |
| 54 | GAL05 | 42.9897 | 0.6472 | obstetricans | 2 | srt | srt | sr |
| 55 | CHAC01 | 43.0351 | 0.9658 | obstetricans | 2 | srt | srt | sr |
| 55 | CHAC02 | 43.0351 | 0.9658 | obstetricans | 2 | srt | srt | sr |
| 55 | CHAC04 | 43.0351 | 0.9658 | obstetricans | 2 | srt | srt | sr |
| 55 | CHAC05 | 43.0351 | 0.9658 | obstetricans | 2 | srt | srt | sr |
| 56 | BET01 | 42.8892 | 1.0565 | obstetricans | 2 | | st | |
| 56 | BET02 | 42.8892 | 1.0565 | obstetricans | 2 | | st | |
| 57 | MNCN81020 | 42.8073 | 1.3826 | obstetricans | 1 | | s | |
| 57 | MNCN81021 | 42.8073 | 1.3826 | obstetricans | 1 | | | |
| 58 | SDS01 | 42.9726 | 1.3885 | obstetricans | 2 | | st | |
| 58 | SDS02 | 42.9726 | 1.3885 | obstetricans | 2 | | st | |
| 58 | SDS03 | 42.9726 | 1.3885 | obstetricans | 2 | | st | |
| 59 | VIE01 | 43.0046 | 1.4722 | admixed | 2 | | st | |

| | | | | | | |
|---|---|---|---|---|---|---|
| 59 | VIE02 | 43.0046 | 1.4722 | admixed | 2 | st |
| 59 | VIE03 | 43.0046 | 1.4722 | admixed | 2 | |
| 59 | VIE04 | 43.0046 | 1.4722 | admixed | 2 | st |
| 59 | VIE05 | 43.0046 | 1.4722 | admixed | 2 | st |
| 59 | VIE06 | 43.0046 | 1.4722 | admixed | 2 | st |
| 60 | RECO01 | 42.9574 | 1.5387 | admixed | 2 | st |
| 60 | RECO03 | 42.9574 | 1.5387 | admixed | 2 | st |
| 61 | BLAN01 | 42.9479 | 1.5582 | admixed | 2 | st |
| 61 | BLAN02 | 42.9479 | 1.5582 | admixed | 2 | st |
| 61 | BLAN03 | 42.9479 | 1.5582 | admixed | 2 | st |
| 62 | BECQ01 | 42.9450 | 1.5789 | admixed | 2 | st |
| 62 | BECQ02 | 42.9450 | 1.5789 | admixed | 2 | st |
| 62 | BECQ03 | 42.9450 | 1.5789 | admixed | 2 | st |
| 63 | CARA01 | 42.9419 | 1.6712 | admixed | 2 | st |
| 63 | CARA02 | 42.9419 | 1.6712 | admixed | 2 | st |
| 63 | CARA03 | 42.9419 | 1.6712 | admixed | 2 | st |
| 64 | CIR01 | 42.9416 | 1.7046 | admixed | 2 | st |
| 64 | CIR02 | 42.9416 | 1.7046 | admixed | 2 | st |
| 64 | CIR03 | 42.9416 | 1.7046 | admixed | 2 | st |
| 64 | CIR04 | 42.9416 | 1.7046 | admixed | 2 | st |
| 64 | CIR05 | 42.9416 | 1.7046 | admixed | 2 | st |
| 65 | ROQ01 | 42.9355 | 1.7553 | admixed | 2 | st |
| 65 | ROQ02 | 42.9355 | 1.7553 | admixed | 2 | st |
| 65 | ROQ05 | 42.9355 | 1.7553 | admixed | 2 | st |
| 65 | ROQ06 | 42.9355 | 1.7553 | admixed | 2 | st |
| 65 | ROQ07 | 42.9355 | 1.7553 | admixed | 2 | st |
| 66 | BEN01 | 42.9061 | 1.8506 | admixed | 2 | st |
| 66 | BEN03 | 42.9061 | 1.8506 | admixed | 2 | st |
| 67 | PEC02 | 42.7442 | 1.7867 | admixed | 2 | s |
| 67 | PEC03 | 42.7442 | 1.7867 | admixed | 2 | s |
| 68 | MAL01 | 42.8020 | 2.0331 | almogavarii | 2 | st |

| | | | | | | | |
|---|---|---|---|---|---|---|---|
| 68 | MAL04 | 42.8020 | 2.0331 | almogavarii | 2 | | st |
| 68 | MAL05 | 42.8020 | 2.0331 | almogavarii | 2 | | st |
| 69 | ROD04 | 42.7984 | 2.0697 | almogavarii | 2 | | st |
| 70 | FONT01 | 42.7687 | 2.0844 | almogavarii | 2 | | st |
| 71 | COU01 | 42.8627 | 2.1254 | almogavarii | 2 | | st |
| 71 | COU02 | 42.8627 | 2.1254 | almogavarii | 2 | | st |
| 71 | COU04 | 42.8627 | 2.1254 | almogavarii | 2 | | st |
| 71 | COU05 | 42.8627 | 2.1254 | almogavarii | 2 | | st |
| 72 | BEV.14083 | 43.0063 | -1.0281 | admixed | 1 | st | |
| 73 | MNCN68405 | 43.0457 | -1.0736 | admixed | 3 | st | |
| 73 | MNCN68406 | 43.0457 | -1.0736 | admixed | 3 | st | |
| 74 | MNCN150803 | 43.0126 | -1.1862 | admixed | 3 | st | |
| 74 | MNCN150804 | 43.0126 | -1.1862 | admixed | 3 | st | |
| 75 | MNCN150798 | 42.9724 | -1.3516 | admixed | 3 | st | |
| 75 | MNCN150799 | 42.9724 | -1.3516 | admixed | 3 | st | |
| 76 | MNCN150796 | 43.0277 | -1.4648 | admixed | 3 | st | |
| 76 | MNCN150797 | 43.0277 | -1.4648 | admixed | 3 | st | |
| 77 | MNCN150791 | 43.2055 | -1.4455 | admixed | 3 | st | |
| 77 | MNCN150792 | 43.2055 | -1.4455 | admixed | 3 | st | |
| 78 | MNCN150780 | 43.2485 | -1.5820 | admixed | 3 | st | |
| 78 | MNCN150781 | 43.2485 | -1.5820 | admixed | 3 | st | |
| 79 | MNCN150775 | 43.3210 | -1.8550 | admixed | 3 | st | |
| 79 | MNCN150776 | 43.3210 | -1.8550 | admixed | 3 | st | |
| 80 | MNCN150749 | 42.9227 | -1.8187 | admixed | 3 | st | |
| 80 | MNCN150750 | 42.9227 | -1.8187 | admixed | 3 | st | |
| 81 | MNCN150742 | 42.9645 | -1.9795 | admixed | 3 | st | |
| 81 | MNCN150743 | 42.9645 | -1.9795 | admixed | 3 | st | |
| 82 | MNCN150727 | 43.1918 | -2.3628 | admixed | 3 | st | |
| 82 | MNCN150728 | 43.1918 | -2.3628 | admixed | 3 | st | |
| 83 | MNCN150723 | 42.9976 | -2.4525 | admixed | 3 | st | |
| 83 | MNCN150724 | 42.9976 | -2.4525 | admixed | 3 | st | |

| | | | | | | | |
|---|---|---|---|---|---|---|---|
| 84 | MNCN150770 | 42.7810 | -2.6750 | admixed | 3 | | st |
| 85 | MNCN67930 | 43.0875 | -2.7921 | admixed | 3 | | st |
| 85 | MNCN67932 | 43.0875 | -2.7921 | admixed | 3 | | st |
| 85 | MNCN67933 | 43.0875 | -2.7921 | admixed | 3 | | st |
| 86 | MNCN150707 | 43.2083 | -2.9105 | admixed | 3 | | st |
| 86 | MNCN150708 | 43.2083 | -2.9105 | admixed | 3 | | st |
| 86 | MNCN150709 | 43.2083 | -2.9105 | admixed | 3 | | st |
| 87 | MNCN150696 | 42.9411 | -3.0007 | admixed | 3 | | st |
| 88 | MNCN150691 | 43.1648 | -3.2288 | admixed | 3 | | st |
| 89 | PANDO01 | 43.2082 | -3.3187 | admixed | 1 | | st |
| 89 | PANDO03 | 43.2082 | -3.3187 | admixed | 3 | | st |
| 89 | PANDO04 | 43.2082 | -3.3187 | admixed | 3 | | |
| 90 | MNCN150686 | 43.0670 | -3.3188 | pertinax | 3 | | st |
| 91 | MNCN150675 | 43.0003 | -3.4519 | admixed | 3 | | |
| 91 | MNCN150676 | 43.0003 | -3.4519 | admixed | 3 | | st |
| 92 | MNCN150660 | 42.9370 | -3.9001 | pertinax | 3 | | |
| 92 | MNCN150661 | 42.9370 | -3.9001 | pertinax | 3 | | st |
| 92 | MNCN150662 | 42.9370 | -3.9001 | pertinax | 3 | | st |
| 93 | BEV.12756 | 43.0298 | -3.9454 | pertinax | 1 | | st |
| 94 | GVA9189 | 43.2595 | -3.9813 | admixed | 3 | | st |
| 95 | MNCN69013 | 43.3386 | -4.1339 | pertinax | 1 | | st |
| 96 | GVA6523 | 42.8753 | -4.3636 | pertinax | 3 | | st |
| 97 | GVA6526 | 42.8822 | -4.5478 | pertinax | 3 | | st |
| 98 | MNCN150654 | 42.8511 | -4.7473 | pertinax | 3 | | st |
| 99 | AO31 | 43.0129 | -4.7639 | pertinax | 3 | | |
| 100 | MNCN150648 | 42.9273 | -4.8872 | pertinax | 3 | sr | srt |
| 101 | MNCN150644 | 43.0013 | -5.0777 | pertinax | 3 | sr | srt |
| 102 | AO81 | 43.2102 | -4.7196 | pertinax | 3 | srt | srt |
| 103 | AO121 | 43.2064 | -4.7671 | pertinax | 3 | srt | srt |
| 104 | AO41 | 43.1652 | -4.8528 | pertinax | 3 | srt | srt |
| 105 | AO51 | 43.1508 | -4.8480 | pertinax | 3 | srt | srt |

| | | | | | | | | |
|---|---|---|---|---|---|---|---|---|
| 106 | AO61 | 43.2227 | -4.9921 | pertinax | 3 | srt | | sr |
| 107 | AO171 | 43.1854 | -5.1405 | pertinax | 3 | srt | | sr |
| 108 | AO251 | 43.2392 | -5.2927 | pertinax | 3 | srt | | sr |
| 109 | AO231 | 43.1863 | -5.5642 | pertinax | 3 | srt | | sr |
| 110 | AO241 | 43.1508 | -5.5840 | pertinax | 3 | srt | | sr |
| 110 | AO242 | 43.1508 | -5.5840 | pertinax | 3 | srt | | sr |
| 111 | AO221 | 43.1190 | -5.5404 | pertinax | 3 | srt | | sr |
| 112 | MNCN150630 | 42.8536 | -5.4957 | pertinax | 3 | | | |
| 113 | AO261 | 42.9903 | -5.9054 | admixed | 3 | st | | |
| 113 | AO262 | 42.9903 | -5.9054 | admixed | 3 | st | | |
| 114 | MNCN150625 | 43.0603 | -6.0021 | pertinax | 3 | st | | |
| 115 | MNCN150620 | 43.0387 | -6.0052 | admixed | 3 | st | | |
| 116 | MNCN150615 | 42.9435 | -6.1356 | admixed | 3 | st | | |
| 116 | MNCN150616 | 42.9435 | -6.1356 | admixed | 3 | st | | |
| 116 | MNCN150617 | 42.9435 | -6.1356 | admixed | 3 | st | | |
| 117 | MNCN150609 | 43.0060 | -6.3269 | admixed | 3 | st | | |
| 117 | MNCN150610 | 43.0060 | -6.3269 | admixed | 3 | st | | |
| 118 | MNCN150604 | 42.9838 | -6.4078 | admixed | 3 | st | | |
| 118 | MNCN150605 | 42.9838 | -6.4078 | admixed | 3 | st | | |
| 118 | MNCN150606 | 42.9838 | -6.4078 | admixed | 3 | st | | |
| 119 | ILNO01 | 43.3100 | -6.8685 | boscai | 3 | st | | |
| 119 | ILNO02 | 43.3100 | -6.8685 | boscai | 3 | st | | |
| 119 | ILNO03 | 43.3100 | -6.8685 | boscai | 3 | st | | |
| 119 | ILNO04 | 43.3100 | -6.8685 | boscai | 3 | st | | |
| 120 | BALEI01 | 43.0328 | -7.1934 | boscai | 3 | srt | sr | |
| 120 | BALEI02 | 43.0328 | -7.1934 | boscai | 3 | srt | sr | |
| 120 | BALEI03 | 43.0328 | -7.1934 | boscai | 3 | srt | sr | |
| 121 | MNCN81866 | 42.9239 | -7.3889 | boscai | 3 | srt | sr | |
| 122 | PAST01 | 43.3707 | -7.3441 | boscai | 3 | srt | sr | |
| 122 | PAST02 | 43.3707 | -7.3441 | boscai | 3 | srt | sr | |
| 123 | MVTIS33289 | 43.7200 | -7.9400 | boscai | 3 | srt | sr | |

| | | | | | | | |
|---|---|---|---|---|---|---|---|
| 124 | GVA9126 | 43.2316 | -8.8767 | boscai | 3 | srt | sr |
| 125 | GVA8927 | 42.7450 | -8.5560 | boscai | 3 | | |
| 126 | MNCN68115 | 42.0500 | -8.6333 | boscai | 1 | sr | srt |
| 127 | CASG01 | 42.4466 | -7.7499 | boscai | 3 | sr | sr |
| 127 | CASG06 | 42.4466 | -7.7499 | boscai | 3 | sr | sr |
| 128 | VILA01 | 42.1982 | -7.6392 | boscai | 3 | sr | sr |
| 129 | MNCN68105 | 41.8103 | -7.4221 | boscai | 1 | sr | sr |
| 130 | BRIT01 | 41.4583 | -8.3765 | boscai | 3 | | st |
| 130 | BRIT02 | 41.4583 | -8.3765 | boscai | 3 | | st |
| 130 | BRIT03 | 41.4583 | -8.3765 | boscai | 3 | | st |
| 131 | LOUS02 | 41.2744 | -8.2746 | boscai | 3 | | st |
| 132 | ALF01 | 41.2351 | -8.5100 | boscai | 3 | | st |
| 132 | ALF02 | 41.2351 | -8.5100 | boscai | 3 | | st |
| 133 | VALAN01 | 41.1573 | -8.4838 | boscai | 3 | | st |
| 133 | VALAN02 | 41.1573 | -8.4838 | boscai | 3 | | st |
| 133 | VALAN03 | 41.1573 | -8.4838 | boscai | 3 | | st |
| 134 | PRT01 | 41.1593 | -8.5890 | admixed | 3 | | st |
| 134 | PRT02 | 41.1593 | -8.5890 | admixed | 3 | | st |
| 134 | PRT03 | 41.1593 | -8.5890 | admixed | 3 | | st |
| 135 | VNG01 | 41.1112 | -8.5947 | admixed | 3 | | st |
| 135 | VNG02 | 41.1112 | -8.5947 | admixed | 3 | | st |
| 135 | VNG05 | 41.1112 | -8.5947 | admixed | 3 | | st |
| 135 | VNG06 | 41.1112 | -8.5947 | admixed | 3 | | st |
| 136 | REAL01 | 40.9974 | -8.2754 | lusitanicus | 3 | | st |
| 136 | REAL02 | 40.9974 | -8.2754 | lusitanicus | 3 | | st |
| 136 | REAL03 | 40.9974 | -8.2754 | lusitanicus | 3 | | st |
| 137 | AROU01 | 40.9296 | -8.2562 | lusitanicus | 3 | | st |
| 137 | AROU02 | 40.9296 | -8.2562 | lusitanicus | 3 | | st |
| 138 | GRI01 | 40.9614 | -8.1058 | lusitanicus | 3 | | st |
| 139 | UL01 | 40.8148 | -8.4990 | lusitanicus | 3 | | |
| 139 | UL02 | 40.8148 | -8.4990 | lusitanicus | 3 | | st |

| | | | | | | | | |
|---|---|---|---|---|---|---|---|---|
| 139 | UL04 | 40.8148 | -8.4990 | lusitanicus | 3 | st | | |
| 140 | RDM01 | 40.3982 | -8.2424 | lusitanicus | 3 | srt | sr | s |
| 140 | RDM02 | 40.3982 | -8.2424 | lusitanicus | 3 | srt | sr | s |
| 140 | RDM03 | 40.3982 | -8.2424 | lusitanicus | 3 | srt | sr | s |
| 141 | MNCN80425 | 40.1106 | -8.3785 | lusitanicus | 1 | srt | sr | s |
| 142 | MNCN68622 | 40.3225 | -7.6028 | lusitanicus | 1 | sr | sr | s |
| 143 | MNCN80888 | 40.2115 | -5.7781 | lusitanicus | 3 | sr | srt | s |
| 144 | MNCN68632 | 40.2975 | -5.7192 | lusitanicus | 1 | | | |
| 144 | MNCN68633 | 40.2975 | -5.7192 | lusitanicus | 3 | sr | srt | s |
| 144 | MNCN68634 | 40.2975 | -5.7192 | lusitanicus | 3 | sr | srt | s |
| 145 | MNCN80419 | 40.3302 | -5.1158 | lusitanicus | 3 | | | |
| 145 | MNCN80420 | 40.3302 | -5.1158 | lusitanicus | 3 | | st | s |
| 146 | AREN01 | 40.2541 | -5.0607 | lusitanicus | 3 | | | |
| 146 | AREN04/MNCN50839 | 40.2541 | -5.0607 | lusitanicus | 3 | | | |
| 146 | AREN04b/MNCN50839 | 40.2541 | -5.0607 | lusitanicus | 3 | | | |
| 147 | MNCN119092 | 40.3606 | -4.8992 | lusitanicus | 1 | | st | s |
| 148 | MNCN150755 | 40.7444 | -4.0451 | pertinax | 3 | | st | s |
| 149 | MNCN150756 | 40.7497 | -4.0377 | pertinax | 3 | | st | s |
| 149 | MNCN150757 | 40.7497 | -4.0377 | pertinax | 3 | | | |
| 149 | MNCN150758 | 40.7497 | -4.0377 | pertinax | 3 | | st | s |
| 150 | MNCN67899 | 40.8036 | -4.0233 | pertinax | 3 | | | |
| 150 | MNCN67900 | 40.8036 | -4.0233 | pertinax | 3 | | st | s |
| 150 | MNCN67903 | 40.8036 | -4.0233 | pertinax | 3 | | st | s |
| 151 | MNCN67867 | 40.9793 | -3.8441 | pertinax | 3 | | st | s |
| 151 | MNCN67869 | 40.9793 | -3.8441 | pertinax | 3 | | | |
| 152 | MNCN150761 | 41.1100 | -3.5931 | pertinax | 3 | | st | s |
| 152 | MNCN150762 | 41.1100 | -3.5931 | pertinax | 3 | | st | s |
| 153 | MNCN67815 | 41.2367 | -3.3498 | pertinax | 3 | | | |
| 153 | MNCN67817 | 41.2367 | -3.3498 | pertinax | 3 | | srt | s |
| 154 | MNCN67820 | 41.2297 | -3.2921 | pertinax | 3 | | srt | s |
| 154 | MNCN67821 | 41.2297 | -3.2921 | pertinax | 3 | | srt | s |

| | | | | | | | | |
|---|---|---|---|---|---|---|---|---|
| 155 | MNCN67939 | 41.2273 | -2.9558 | pertinax | 3 | srt | s | |
| 155 | MNCN67940 | 41.2273 | -2.9558 | pertinax | 3 | srt | s | |
| 156 | MNCN68466 | 42.0058 | -1.5365 | pertinax | 1 | sr | | |
| 157 | MNCN67426 | 40.6473 | -1.9073 | pertinax | 1 | sr | | s |
| 158 | MNCN67537 | 40.5163 | -0.4594 | pertinax | 1 | sr | | s |
| 159 | MNCN67523 | 39.8149 | -0.6997 | pertinax | 1 | sr | | s |
| 160 | BEV9964 | 39.7054 | -0.2598 | pertinax | 1 | sr | | s |
| 161 | MNCN67454 | 38.9290 | -0.8550 | pertinax | 3 | sr | | s |
| 162 | MNCN67430 | 39.0029 | -1.3042 | pertinax | 3 | sr | | s |
| 163 | MNCN150821 | 39.1078 | -2.4869 | pertinax | 3 | sr | | s |
| 163 | MNCN150823 | 39.1078 | -2.4869 | pertinax | 3 | sr | | s |
| - | MNCN68027 | 38.3092 | -5.0004 | cisternasii | 1 | | s | |
| - | MNCN150769 | 40.6445 | -4.1340 | cisternasii | 3 | | s | |
| - | MNCN150766 | 40.6578 | -4.1345 | cisternasii | 3 | | s | |
| - | MNCN150768 | 40.6445 | -4.1340 | cisternasii | 3 | | s | |
| - | MNCN81858 | 38.9946 | -6.3408 | cisternasii | 1 | | s | |
| - | MNCN68046 | 38.5554 | -7.9276 | cisternasii | 1 | | s | |
| - | MNCN118912 | 40.317 | -4.3790 | cisternasii | 1 | | s | |
| - | BEV12834 | 38.1140 | -4.0476 | cisternasii | 1 | | s | |
| - | MNCN150767 | 40.6445 | -4.1340 | cisternasii | 3 | | s | |
| - | MNCN80787 | 39.1300 | -6.8500 | cisternasii | 1 | | s | |
| - | MNCN81899 | 36.8721 | -4.0643 | dickhilleni | 1 | | | s |
| - | MNCN103299 | 38.0621 | -1.7090 | dickhilleni | 1 | | | s |
| - | MNCN66282 | 38.1491 | -2.1356 | dickhilleni | 1 | | | s |
| - | MNCN68006 | 38.5451 | -2.3826 | dickhilleni | 1 | | | s |
| - | BEV.T4781 | 34.0733 | -4.1314 | maurus | 1 | | | |
| - | AmaurAH | 35.5203 | -5.3410 | maurus | 1 | | | |
| - | MNCN68108 | - | - | muletensis | 1 | | | |
| - | MVTIS3217 | - | - | muletensis | 1 | | | |

[1] Ambu et al. 2023; [2] Ambu & Dufresnes resubmitted; [3] this study

**File S2**: Bioacoustic measurements.

| Individual | Taxon | Y | X | notes | DF | ND | RT | PR | Source |
|---|---|---|---|---|---|---|---|---|---|
| ARQ11 | *almogavarii* | 42.95242 | 2.38355 | 6 | 1550.4 | 0.091 | 0.012 | 1527 | own record |
| BEN04 | *almogavarii* | 42.90614 | 1.85062 | 6 | 1378.1 | 0.086 | 0.013 | 1372 | own record |
| BEN05 | *almogavarii* | 42.90614 | 1.85062 | 5 | 1550.4 | 0.086 | 0.007 | 1522 | own record |
| BEN06 | *almogavarii* | 42.90614 | 1.85062 | 5 | 1610.7 | 0.083 | 0.011 | 1598 | own record |
| BOUG11 | *almogavarii* | 43.24066 | 2.430638 | 2 | 1312.0 | 0.096 | 0.012 | 1325 | own record |
| CARCAS01 | *almogavarii* | 43.20625 | 2.362223 | 6 | 1464.0 | 0.085 | 0.013 | 1440 | own record |
| COU21_01 | *almogavarii* | 42.86187 | 2.125124 | 5 | 1292.0 | 0.105 | 0.007 | 1250 | own record |
| COU21_02 | *almogavarii* | 42.86187 | 2.125124 | 6 | 1292.0 | 0.092 | 0.007 | 1290 | own record |
| COU21_03 | *almogavarii* | 42.86187 | 2.125124 | 6 | 1378.0 | 0.085 | 0.013 | 1310 | own record |
| COU21_04 | *almogavarii* | 42.86187 | 2.125124 | 6 | 1206.0 | 0.078 | 0.009 | 1197 | own record |
| COU21_05 | *almogavarii* | 42.86187 | 2.125124 | 2 | 1378.0 | 0.074 | 0.010 | 1279 | own record |
| COU21_06 | *almogavarii* | 42.86187 | 2.125124 | 3 | 1378.0 | 0.085 | 0.010 | 1303 | own record |
| GREF12 | *almogavarii* | 43.06815 | 2.375177 | 6 | 1464.0 | 0.084 | 0.011 | 1488 | own record |
| GREF13 | *almogavarii* | 43.06815 | 2.375177 | 8 | 1551.0 | 0.076 | 0.010 | 1561 | own record |
| GREF14 | *almogavarii* | 43.06815 | 2.375177 | 6 | 1636.0 | 0.090 | 0.009 | 1609 | own record |
| GREF15 | *almogavarii* | 43.06815 | 2.375177 | 6 | 1464.0 | 0.100 | 0.010 | 1432 | own record |
| SOU21_01 | *almogavarii* | 42.73314 | 2.426531 | 6 | 1378.0 | 0.073 | 0.004 | 1342 | own record |
| VGLY01 | *almogavarii* | 43.28379 | 2.442997 | 6 | 1464.0 | 0.136 | 0.008 | 1424 | own record |
| VGLY09 | *almogavarii* | 43.28379 | 2.442997 | 6 | 1378.0 | 0.092 | 0.007 | 1408 | own record |
| VGLY10 | *almogavarii* | 43.28379 | 2.442997 | 6 | 1292.0 | 0.096 | 0.008 | 1268 | own record |
| VGLY11 | *almogavarii* | 43.28379 | 2.442997 | 6 | 1378.0 | 0.112 | 0.009 | 1341 | own record |
| VGLY12 | *almogavarii* | 43.28379 | 2.442997 | 6 | 1550.0 | 0.08 | 0.005 | 1539 | own record |
| ALM01 | *almogavarii* | 41.39382 | 1.914294 | 5 | 1464.2 | 0.095 | 0.010 | 1463 | iNaturalist |
| ALM02 | *almogavarii* | 42.95264 | 2.556986 | 5 | 1359.4 | 0.090 | 0.009 | 1330 | iNaturalist |
| ALM03 | *almogavarii* | 41.36628 | 2.151931 | 4 | 1292.0 | 0.121 | 0.013 | 1290 | iNaturalist |
| ALM04 | *almogavarii* | 41.48305 | 2.074496 | 4 | 1550.4 | 0.080 | 0.009 | 1530 | iNaturalist |
| ALM05 | *almogavarii* | 42.6656 | 2.651315 | 3 | 1378.1 | 0.087 | 0.008 | 1392 | iNaturalist |
| ALM06 | *almogavarii* | 41.48296 | 2.07393 | 6 | 1622.2 | 0.075 | 0.008 | 1605 | iNaturalist |
| ALM07 | *almogavarii* | 41.48296 | 2.07393 | 6 | 1636.5 | 0.084 | 0.008 | 1622 | iNaturalist |

| | | | | | | | | | |
|---|---|---|---|---|---|---|---|---|---|
| ALM08 | *almogavarii* | 41.46925 | 2.096489 | 5 | 1335.1 | 0.11 | 0.01 | 1329 | iNaturalist |
| ALM09 | *almogavarii* | 41.46925 | 2.096489 | 5 | 1335.1 | 0.115 | 0.016 | 1298 | iNaturalist |
| ALM10 | *almogavarii* | 41.45159 | 2.111267 | 3 | 1292.0 | 0.110 | 0.007 | 1288 | iNaturalist |
| ALM11 | *almogavarii* | 41.4606 | 2.036483 | 6 | 1356.6 | 0.104 | 0.010 | 1343 | iNaturalist |
| ALM12 | *almogavarii* | 41.36719 | 2.164661 | 4 | 1453.1 | 0.071 | 0.006 | 1465 | iNaturalist |
| ALM13 | *almogavarii* | 41.26035 | 1.439437 | 2 | 1335.1 | 0.107 | 0.009 | 1315 | iNaturalist |
| ALM14 | *almogavarii* | 41.36809 | 2.158263 | 3 | 1406.2 | 0.098 | 0.008 | 1429 | iNaturalist |
| ALM15 | *almogavarii* | 41.36809 | 2.158263 | 2 | 1359.4 | 0.094 | 0.007 | 1356 | iNaturalist |
| GORG01 | *inigoi* | 42.70471 | -0.32141 | 6 | 1076.7 | 0.107 | 0.009 | 1071 | own record |
| PIED13 | *inigoi* | 42.6968 | -0.35358 | 6 | 1248.9 | 0.096 | 0.006 | 1230 | own record |
| SOBA02 | *inigoi* | 42.48787 | -0.26006 | 6 | 1327.9 | 0.094 | 0.019 | 1311 | own record |
| SOBA03 | *inigoi* | 42.48787 | -0.26006 | 4 | 1205.9 | 0.097 | 0.016 | 1195 | own record |
| INIG03 | *inigoi* | 42.59656 | 0.128031 | 5 | 1326.5 | 0.124 | 0.016 | 1306 | iNaturalist |
| BOSC01 | *boscai* | 41.50472 | -8.23778 | 6 | 1227.4 | 0.081 | 0.011 | 1221 | iNaturalist |
| BOSC02 | *boscai* | 41.50472 | -8.23778 | 6 | 1248.9 | 0.081 | 0.013 | 1223 | iNaturalist |
| BOSC03 | *boscai* | 41.63311 | -7.09152 | 5 | 1180.0 | 0.074 | 0.009 | 1171 | iNaturalist |
| BOSC04 | *boscai* | 41.21059 | -8.79549 | 5 | 1312.5 | 0.086 | 0.008 | 1306 | iNaturalist |
| BOSC05 | *boscai* | 41.54366 | -7.64882 | 4 | 1265.6 | 0.088 | 0.008 | 1240 | iNaturalist |
| BOSC06 | *boscai* | 41.54366 | -7.64882 | 2 | 1429.7 | 0.076 | 0.008 | 1421 | iNaturalist |
| BOSC07 | *boscai* | 41.57166 | -7.82232 | 6 | 1406.2 | 0.073 | 0.006 | 1386 | iNaturalist |
| BOSC08 | *boscai* | 41.57166 | -7.82232 | 6 | 1359.4 | 0.084 | 0.011 | 1367 | iNaturalist |
| BOSC09 | *boscai* | 42.87661 | -8.5503 | 2 | 1292.0 | 0.101 | 0.012 | 1294 | iNaturalist |
| BOSC10 | *boscai* | 43.27999 | -8.21986 | 4 | 1359.4 | 0.099 | 0.009 | 1355 | iNaturalist |
| BOSC11 | *boscai* | 41.61534 | -6.40788 | 3 | 1312.5 | 0.091 | 0.017 | 1290 | iNaturalist |
| BOSC12 | *boscai* | 41.61534 | -6.40788 | 2 | 1265.6 | 0.085 | 0.018 | 1282 | iNaturalist |
| BOSC13 | *boscai* | 42.19239 | -7.79827 | 5 | 1218.7 | 0.09 | 0.009 | 1203 | iNaturalist |
| BOSC14 | *boscai* | 41.80926 | -8.41928 | 4 | 1406.2 | 0.084 | 0.011 | 1395 | iNaturalist |
| BOSC15 | *boscai* | 41.55788 | -8.66422 | 3 | 1359.4 | 0.076 | 0.007 | 1338 | iNaturalist |
| LUSI01 | *lusitanicus* | 41.11117 | -8.59451 | 6 | 1284.8 | 0.094 | 0.014 | 1279 | iNaturalist |
| LUSI02 | *lusitanicus* | 39.93565 | -8.18846 | 2 | 1227.4 | 0.078 | 0.007 | 1230 | iNaturalist |
| LUSI03 | *lusitanicus* | 40.30123 | -7.82208 | 2 | 1336.0 | 0.088 | 0.010 | 1318 | iNaturalist |
| LUSI04 | *lusitanicus* | 40.40187 | -7.53715 | 1 | 1265.6 | 0.112 | 0.018 | 1295 | iNaturalist |
| LUSI05 | *lusitanicus* | 40.40187 | -7.53715 | 2 | 1218.8 | 0.095 | 0.015 | 1211 | iNaturalist |
| LUSI06 | *lusitanicus* | 40.29729 | -7.23282 | 4 | 1421.2 | 0.069 | 0.010 | 1399 | iNaturalist |

| | | | | | | | | | |
|---|---|---|---|---|---|---|---|---|---|
| LUSI07 | *lusitanicus* | 40.29761 | -7.23285 | 1 | 1312.5 | 0.086 | 0.01 | 1302 | iNaturalist |
| LUSI08 | *lusitanicus* | 40.29761 | -7.23285 | 2 | 1406.2 | 0.069 | 0.008 | 1391 | iNaturalist |
| LUSI09 | *lusitanicus* | 41.07673 | -8.5993 | 5 | 1464.3 | 0.074 | 0.006 | 1462 | iNaturalist |
| LUSI10 | *lusitanicus* | 40.12008 | -8.23988 | 3 | 1335.1 | 0.091 | 0.008 | 1321 | iNaturalist |
| LUSI11 | *lusitanicus* | 40.12008 | -8.23988 | 3 | 1378.1 | 0.106 | 0.009 | 1361 | iNaturalist |
| AEB06 | *obstetricans* | 42.89698 | 1.041388 | 6 | 1270.5 | 0.08 | 0.008 | 1254 | own record |
| AGO05 | *obstetricans* | 43.01215 | 1.498627 | 6 | 1162.8 | 0.068 | 0.009 | 1174 | own record |
| BET05 | *obstetricans* | 42.88917 | 1.05645 | 6 | 1119.7 | 0.081 | 0.008 | 1130 | own record |
| BET06 | *obstetricans* | 42.88917 | 1.05645 | 6 | 1119.7 | 0.081 | 0.012 | 1121 | own record |
| BET07 | *obstetricans* | 42.88917 | 1.05645 | 3 | 1292.0 | 0.081 | 0.009 | 1283 | own record |
| DSA03 | *obstetricans* | 43.01689 | 1.339885 | 6 | 1335.1 | 0.084 | 0.009 | 1329 | own record |
| DSA04 | *obstetricans* | 43.01689 | 1.339885 | 6 | 1507.3 | 0.086 | 0.008 | 1505 | own record |
| DSA05 | *obstetricans* | 43.01689 | 1.339885 | 6 | 1507.3 | 0.088 | 0.010 | 1481 | own record |
| GAN22_01 | *obstetricans* | 43.22841 | -0.41109 | 6 | 1636.5 | 0.095 | 0.010 | 1636 | own record |
| GAN22_02 | *obstetricans* | 43.22841 | -0.41109 | 6 | 1636.5 | 0.097 | 0.009 | 1649 | own record |
| GAN22_03 | *obstetricans* | 43.22841 | -0.41109 | 6 | 1378.1 | 0.088 | 0.008 | 1387 | own record |
| JEUR05 | *obstetricans* | 46.36995 | 5.715916 | 9 | 1277.6 | 0.089 | 0.008 | 1243 | own record |
| JEUR06 | *obstetricans* | 46.36995 | 5.715916 | 6 | 1378.1 | 0.104 | 0.011 | 1350 | own record |
| LAR01 | *obstetricans* | 43.83256 | 3.504805 | 4 | 1206.0 | 0.133 | 0.012 | 1208 | own record |
| LAR02 | *obstetricans* | 43.83256 | 3.504805 | 4 | 1206.0 | 0.127 | 0.007 | 1213 | own record |
| LAR03 | *obstetricans* | 43.83256 | 3.504805 | 5 | 1206.0 | 0.129 | 0.007 | 1173 | own record |
| LAR04 | *obstetricans* | 43.83256 | 3.504805 | 5 | 1292.0 | 0.121 | 0.006 | 1299 | own record |
| MOS01 | *obstetricans* | 46.53463 | 6.366401 | 6 | 1378.0 | 0.104 | 0.003 | 1329 | own record |
| MOULIS01 | *obstetricans* | 42.9607 | 1.091897 | 6 | 1248.9 | 0.076 | 0.010 | 1256 | own record |
| SALV03 | *obstetricans* | 43.56664 | 2.738563 | 6 | 1292.0 | 0.116 | 0.007 | 1262 | own record |
| SALV04 | *obstetricans* | 43.56664 | 2.738563 | 6 | 1292.0 | 0.141 | 0.006 | 1260 | own record |
| VIE22_01 | *obstetricans* | 43.00461 | 1.472231 | 6 | 1292.0 | 0.106 | 0.012 | 1274 | own record |
| VIE22_02 | *obstetricans* | 43.00461 | 1.472231 | 6 | 1292.0 | 0.088 | 0.008 | 1271 | own record |
| VIE23_01 | *obstetricans* | 43.00461 | 1.472231 | 6 | 1335.1 | 0.067 | 0.006 | 1333 | own record |
| VIE23_02 | *obstetricans* | 43.00461 | 1.472231 | 6 | 1205.9 | 0.082 | 0.009 | 1207 | own record |
| VIE23_03 | *obstetricans* | 43.00461 | 1.472231 | 6 | 1248.9 | 0.086 | 0.010 | 1244 | own record |
| VIE23_04 | *obstetricans* | 43.00461 | 1.472231 | 6 | 1421.2 | 0.079 | 0.005 | 1398 | own record |
| OBST01 | *obstetricans* | 49.20608 | -0.69152 | 6 | 1349.4 | 0.090 | 0.017 | 1335 | iNaturalist |
| OBST02 | *obstetricans* | 48.23039 | 4.714629 | 4 | 1312.5 | 0.109 | 0.013 | 1324 | iNaturalist |

| | | | | | | | | | |
|---|---|---|---|---|---|---|---|---|---|
| OBST03 | *obstetricans* | 44.81792 | -0.21202 | 5 | 1335.1 | 0.114 | 0.016 | 1316 | iNaturalist |
| OBST04 | *obstetricans* | 44.81792 | -0.21202 | 5 | 1197.3 | 0.106 | 0.010 | 1191 | iNaturalist |
| OBST05 | *obstetricans* | 46.5758 | 0.333777 | 6 | 1312.5 | 0.072 | 0.009 | 1317 | iNaturalist |
| OBST06 | *obstetricans* | 46.5758 | 0.333777 | 5 | 1406.2 | 0.075 | 0.011 | 1409 | iNaturalist |
| OBST07 | *obstetricans* | 49.84226 | 6.319592 | 6 | 1205.9 | 0.134 | 0.008 | 1213 | iNaturalist |
| OBST08 | *obstetricans* | 43.57528 | 1.374659 | 5 | 1350.0 | 0.103 | 0.009 | 1338 | iNaturalist |
| OBST09 | *obstetricans* | 43.57528 | 1.374659 | 5 | 1396.8 | 0.095 | 0.007 | 1402 | iNaturalist |
| OBST10 | *obstetricans* | 43.57528 | 1.374659 | 4 | 1347.7 | 0.106 | 0.009 | 1346 | iNaturalist |
| OBST11 | *obstetricans* | 43.60098 | 1.348861 | 6 | 1550.4 | 0.098 | 0.011 | 1543 | iNaturalist |
| OBST12 | *obstetricans* | 49.92628 | 6.331533 | 5 | 1274.8 | 0.094 | 0.007 | 1253 | iNaturalist |
| OBST13 | *obstetricans* | 48.63678 | -2.06269 | 1 | 1464.3 | 0.107 | 0.011 | 1439 | iNaturalist |
| OBST14 | *obstetricans* | 43.04004 | 1.135093 | 3 | 1205.9 | 0.113 | 0.010 | 1213 | iNaturalist |
| OBST15 | *obstetricans* | 44.59828 | 1.24362 | 6 | 1205.9 | 0.109 | 0.012 | 1204 | iNaturalist |
| OBST16 | *obstetricans* | 44.59828 | 1.24362 | 6 | 1378.1 | 0.095 | 0.014 | 1354 | iNaturalist |
| PERT01 | *pertinax* | 43.39448 | -4.19378 | 4 | 1546.9 | 0.093 | 0.007 | 1555 | iNaturalist |
| PERT02 | *pertinax* | 43.37532 | -4.5629 | 6 | 1601.5 | 0.068 | 0.006 | 1595 | iNaturalist |
| PERT03 | *pertinax* | 43.37532 | -4.5629 | 5 | 1453.1 | 0.063 | 0.007 | 1457 | iNaturalist |
| PERT04 | *pertinax* | 43.37532 | -4.5629 | 6 | 1359.4 | 0.062 | 0.007 | 1346 | iNaturalist |
| PERT05 | *pertinax* | 43.37532 | -4.5629 | 6 | 1453.1 | 0.073 | 0.009 | 1458 | iNaturalist |
| PERT06 | *pertinax* | 43.33429 | -3.55064 | 4 | 1265.6 | 0.108 | 0.011 | 1247 | iNaturalist |
| PERT07 | *pertinax* | 43.33429 | -3.55064 | 6 | 1218.7 | 0.086 | 0.015 | 1187 | iNaturalist |
| PERT08 | *pertinax* | 43.33429 | -3.55064 | 6 | 1406.2 | 0.088 | 0.011 | 1353 | iNaturalist |
| MUL01 | *muletensis* | 39.7987 | 2.7598 | 6 | 1679.6 | 0.082 | 0.005 | 1655 | own record |
| MUL02 | *muletensis* | 39.7987 | 2.7598 | 1 | 1464.3 | 0.069 | 0.006 | 1464 | own record |
| MUL03 | *muletensis* | 39.7987 | 2.7598 | 2 | 1507.3 | 0.067 | 0.006 | 1511 | own record |
| MUL04 | *muletensis* | 39.7987 | 2.7598 | 5 | 1550.4 | 0.071 | 0.006 | 1551 | own record |
| MUL05 | *muletensis* | 39.7987 | 2.7598 | 5 | 1593.5 | 0.076 | 0.006 | 1587 | own record |
| MUL06 | *muletensis* | 39.7987 | 2.7598 | 6 | 1593.5 | 0.069 | 0.006 | 1574 | own record |
| MUL07 | *muletensis* | 39.7987 | 2.7598 | 6 | 1636.5 | 0.092 | 0.006 | 1601 | own record |
| MUL08 | *muletensis* | 39.7987 | 2.7598 | 3 | 1794.4 | 0.082 | 0.004 | 1780 | own record |
| MUL09 | *muletensis* | 39.7987 | 2.7598 | 2 | 1464.3 | 0.083 | 0.004 | 1442 | own record |
| MUL10 | *muletensis* | 39.7987 | 2.7598 | 3 | 1636.5 | 0.068 | 0.005 | 1581 | own record |
| MUL11 | *muletensis* | 39.89394 | 3.086522 | 2 | 1734.4 | 0.114 | 0.009 | 1767 | iNaturalist |
| MUL12 | *muletensis* | 39.7987 | 2.7598 | 6 | 1894.9 | 0.077 | 0.007 | 1843 | AmphibiaWeb |

| | | | | | | | | | |
|---|---|---|---|---|---|---|---|---|---|
| MUL13 | *muletensis* | 39.7987 | 2.7598 | 3 | 1981.1 | 0.077 | 0.006 | 2004 | AmphibiaWeb |
| MUL14 | *muletensis* | 39.7987 | 2.7598 | 4 | 1765.7 | 0.068 | 0.005 | 1774 | AmphibiaWeb |
| MUL15 | *muletensis* | 39.7987 | 2.7598 | 5 | 1851.9 | 0.078 | 0.006 | 1834 | AmphibiaWeb |
| MUL16 | *muletensis* | 39.7987 | 2.7598 | 2 | 1808.8 | 0.069 | 0.006 | 1818 | AmphibiaWeb |
| MUL17 | *muletensis* | 39.7987 | 2.7598 | 1 | 1593.5 | 0.082 | 0.008 | 1573 | Youtube |
| MUL18 | *muletensis* | 39.7987 | 2.7598 | 3 | 1550.4 | 0.08 | 0.008 | 1527 | Youtube |
| MUL19 | *muletensis* | 39.7987 | 2.7598 | 2 | 1636.5 | 0.089 | 0.007 | 1617 | Youtube |
| DICK01 | *dickhilleni* | 37.89277 | -2.89338 | 3 | 1335.1 | 0.225 | 0.011 | 1323 | AmphibiaWeb |
| MAUR01 | *maurus* | 35.54145 | -5.38597 | 2 | 1464.3 | 0.090 | 0.009 | 1439 | own record |
| CIS01 | *cisternasii* | 38.114 | -4.0476 | 6 | 1312.5 | 0.169 | 0.011 | 1279 | own record |
| CIS02 | *cisternasii* | - | - | 6 | 1292.0 | 0.132 | 0.008 | 1289 | Youtube |
| CIS03 | *cisternasii* | - | - | 6 | 1464.3 | 0.109 | 0.008 | 1442 | Youtube |
| CIS04 | *cisternasii* | 38.16948 | -5.16182 | 6 | 1378.1 | 0.160 | 0.013 | 1394 | Soundscape |
| CIS05 | *cisternasii* | 37.89999 | -4.85 | 5 | 1455.7 | 0.105 | 0.010 | 1431 | Youtube |
| CIS06 | *cisternasii* | 37.89999 | -4.85 | 6 | 1335.1 | 0.108 | 0.012 | 1322 | Youtube |
| CIS07 | *cisternasii* | 37.4271 | -7.88254 | 6 | 1218.7 | 0.230 | 0.019 | 1226 | iNaturalist |
| CIS08 | *cisternasii* | 37.4271 | -7.88254 | 6 | 1312.5 | 0.188 | 0.012 | 1306 | iNaturalist |
| CIS09 | *cisternasii* | 37.25957 | -7.53834 | 6 | 1359.4 | 0.119 | 0.013 | 1335 | iNaturalist |
| CIS10 | *cisternasii* | 40.67489 | -3.73583 | 4 | 1359.4 | 0.123 | 0.006 | 1340 | iNaturalist |
| CIS11 | *cisternasii* | 38.9197 | -6.34659 | 3 | 1464.3 | 0.186 | 0.014 | 1508 | AmphibiaWeb |
| CIS12 | *cisternasii* | 38.9197 | -6.34659 | 3 | 1464.3 | 0.176 | 0.009 | 1430 | AmphibiaWeb |
| CIS13 | *cisternasii* | 38.9197 | -6.34659 | 5 | 1498.7 | 0.157 | 0.009 | 1505 | AmphibiaWeb |

**File S3**: Morphological measurements (in mm).

| Voucher | Taxon | SVL | ML | HW | FLL | FML | TBL | FTL | HLL |
|---|---|---|---|---|---|---|---|---|---|
| MNCN.24083 | *almogavarii* | 31.79 | 14.47 | 14.03 | 20.46 | 15.33 | 14.34 | 22.92 | 46.78 |
| MNCN.24084 | *almogavarii* | 33.77 | 14.73 | 13.28 | 22.17 | 15.16 | 14.53 | 23.90 | 48.22 |
| MNCN.24085 | *almogavarii* | 43.36 | 16.64 | 17.03 | 22.94 | 18.57 | 18.18 | 28.56 | 59.66 |
| MNCN.2584 | *almogavarii* | 39.96 | 14.70 | 15.80 | 21.47 | 15.37 | 17.12 | 25.66 | 51.58 |
| MNCN.2585 | *almogavarii* | 35.77 | 14.50 | 14.55 | 22.64 | 17.40 | 16.47 | 26.54 | 53.78 |
| MNCN.2587 | *almogavarii* | 33.32 | 13.42 | 14.53 | 20.05 | 15.33 | 15.61 | 22.33 | 50.50 |
| MNCN.2588 | *almogavarii* | 39.41 | 14.87 | 16.18 | 23.40 | 17.80 | 18.92 | 25.45 | 54.17 |
| MNCN.26318 | *almogavarii* | 37.69 | 14.09 | 15.06 | 21.87 | 16.27 | 17.30 | 21.25 | 53.26 |
| MNCN.9808 | *inigoi* | 34.70 | 14.42 | 13.80 | 18.58 | 13.06 | 15.61 | 21.78 | 44.49 |
| MNCN.9811 | *inigoi* | 42.98 | 14.97 | 16.64 | 21.69 | 17.71 | 16.01 | 24.53 | 53.06 |
| MNCN.9813 | *inigoi* | 40.13 | 15.02 | 15.66 | 25.27 | 17.26 | 16.81 | 27.20 | 56.88 |
| MNCN.9815 | *inigoi* | 39.53 | 13.78 | 15.24 | 21.53 | 17.74 | 17.65 | 25.19 | 51.45 |
| MNCN.9816 | *inigoi* | 39.89 | 15.60 | 16.90 | 21.78 | 16.97 | 16.45 | 23.80 | 49.75 |
| MNCN.9817 | *inigoi* | 35.51 | 12.63 | 15.02 | 19.37 | 13.95 | 14.89 | 21.71 | 46.10 |
| MNCN.9818 | *inigoi* | 38.42 | 13.59 | 14.98 | 20.81 | 14.00 | 15.69 | 25.24 | 51.52 |
| MNCN.9820 | *inigoi* | 39.52 | 15.55 | 16.69 | 20.39 | 16.32 | 18.01 | 28.15 | 52.38 |
| MNCN.9821 | *inigoi* | 39.59 | 14.43 | 15.80 | 20.98 | 13.82 | 16.79 | 24.96 | 49.50 |
| MNCN.9825 | *inigoi* | 37.43 | 13.62 | 14.72 | 18.77 | 17.82 | 15.98 | 24.51 | 50.14 |
| MNCN.11268 | *boscai* | 39.52 | 13.63 | 15.14 | 20.24 | 15.16 | 15.49 | 23.56 | 49.00 |
| MNCN.155 | *boscai* | 39.30 | 14.57 | 16.54 | 19.56 | 15.90 | 14.02 | 24.74 | 49.05 |
| MNCN.41751 | *boscai* | 32.97 | 13.33 | 14.50 | 20.49 | 13.64 | 15.44 | 23.37 | 46.00 |
| NHM.1972.1531 | *boscai* | 39.13 | 14.24 | 15.80 | 20.70 | 16.00 | 17.20 | 21.71 | 49.80 |
| NHM.1972.1532 | *boscai* | 38.55 | 14.50 | 15.86 | 19.16 | 15.09 | 16.52 | 22.34 | 46.10 |
| NHM.1972.1534 | *boscai* | 40.11 | 15.68 | 16.58 | 20.71 | 17.81 | 16.80 | 25.60 | 54.35 |
| NHM.1972.1536 | *boscai* | 33.51 | 13.84 | 14.40 | 17.94 | 14.34 | 14.00 | 20.95 | 45.88 |
| NHM.1972.1538 | *boscai* | 37.60 | 14.17 | 15.10 | 19.09 | 16.71 | 15.57 | 22.81 | 48.75 |
| NHM.1972.1539 | *boscai* | 37.90 | 14.20 | 16.46 | 19.30 | 16.00 | 16.50 | 23.67 | 48.90 |
| NHM.1972.1541 | *boscai* | 38.50 | 13.94 | 16.46 | 21.65 | 16.56 | 16.36 | 24.60 | 50.00 |

| | | | | | | | | | |
|---|---|---|---|---|---|---|---|---|---|
| MNCN.11369 | *lusitanicus* | 33.67 | 13.47 | 14.82 | 19.51 | 15.51 | 13.84 | 21.44 | 46.76 |
| MNCN.13133 | *lusitanicus* | 41.27 | 14.57 | 16.60 | 22.60 | 15.64 | 16.58 | 25.70 | 53.28 |
| MNCN.13134 | *lusitanicus* | 37.60 | 13.51 | 15.55 | 22.72 | 15.24 | 15.67 | 24.43 | 50.90 |
| MNCN.26241 | *lusitanicus* | 44.80 | 16.50 | 16.84 | 21.54 | 13.00 | 17.54 | 25.60 | 53.75 |
| MNCN.314 | *lusitanicus* | 28.50 | 11.79 | 14.03 | 14.37 | 11.36 | 11.56 | 17.66 | 36.31 |
| MNCN.315 | *lusitanicus* | 33.99 | 13.41 | 17.06 | 19.99 | 14.39 | 15.21 | 23.38 | 47.42 |
| MNCN.316 | *lusitanicus* | 30.45 | 13.54 | 14.11 | 17.26 | 14.03 | 13.26 | 21.07 | 42.71 |
| MNCN.317 | *lusitanicus* | 32.33 | 13.64 | 15.13 | 19.07 | 15.09 | 14.08 | 18.39 | 41.66 |
| MNCN.318 | *lusitanicus* | 30.84 | 12.77 | 13.01 | 16.50 | 13.56 | 13.25 | 19.61 | 41.80 |
| MNCN.319 | *lusitanicus* | 32.40 | 13.80 | 15.19 | 17.26 | 14.48 | 13.04 | 20.25 | 42.64 |
| MNHN.1900.123 | *lusitanicus* | 36.02 | 14.02 | 15.90 | 20.50 | 15.24 | 16.25 | 23.45 | 48.67 |
| MNCN.16845 | *obstetricans* | 37.31 | 14.99 | 15.00 | 20.55 | 15.55 | 15.84 | 23.55 | 49.14 |
| MNCN.16846 | *obstetricans* | 35.08 | 13.42 | 14.29 | 21.06 | 15.50 | 15.83 | 25.22 | 50.90 |
| MNCN.16847 | *obstetricans* | 32.07 | 11.82 | 12.68 | 17.60 | 14.02 | 13.50 | 22.20 | 43.80 |
| MNCN.16849 | *obstetricans* | 36.98 | 14.77 | 15.57 | 21.30 | 16.00 | 16.15 | 24.50 | 50.43 |
| MNCN.16850 | *obstetricans* | 34.94 | 13.29 | 15.12 | 19.87 | 14.39 | 15.30 | 23.10 | 46.44 |
| MNCN.26266 | *obstetricans* | 32.36 | 12.91 | 13.30 | 19.68 | 14.27 | 14.63 | 22.58 | 45.20 |
| MNCN.44134 | *obstetricans* | 32.41 | 13.06 | 12.07 | 18.71 | 14.75 | 15.27 | 22.01 | 47.40 |
| MNCN.44313 | *obstetricans* | 31.67 | 12.03 | 13.81 | 21.68 | 13.62 | 14.13 | 23.31 | 47.39 |
| MNCN.44315 | *obstetricans* | 33.22 | 12.33 | 13.25 | 19.18 | 13.93 | 13.57 | 21.32 | 46.45 |
| MNCN.44316 | *obstetricans* | 28.89 | 10.83 | 11.56 | 17.51 | 12.09 | 12.15 | 19.45 | 39.69 |
| MNCN.44318 | *obstetricans* | 27.83 | 10.20 | 9.95 | 15.60 | 12.41 | 12.07 | 18.03 | 39.24 |
| MNCN.44320 | *obstetricans* | 30.30 | 12.86 | 13.80 | 18.64 | 14.04 | 14.91 | 20.50 | 44.51 |
| MNCN.44321 | *obstetricans* | 29.48 | 10.62 | 10.79 | 17.76 | 13.09 | 12.93 | 19.19 | 40.65 |
| MNCN.44325 | *obstetricans* | 31.95 | 12.51 | 13.09 | 18.59 | 14.12 | 14.72 | 19.20 | 43.07 |
| MNCN.44326 | *obstetricans* | 37.75 | 13.47 | 14.21 | 22.87 | 15.49 | 16.30 | 26.23 | 54.20 |
| MNCN.44327 | *obstetricans* | 35.02 | 13.53 | 14.35 | 20.53 | 14.70 | 15.65 | 24.52 | 48.72 |
| MNHN.1908.0216 | *obstetricans* | 32.64 | 11.61 | 12.78 | 21.55 | 15.51 | 14.89 | 22.25 | 47.95 |
| MNHN.1980.1766 | *obstetricans* | 37.71 | 14.96 | 17.10 | 20.94 | 18.39 | 17.81 | 25.84 | 54.97 |
| MNHN.1981.689 | *obstetricans* | 32.78 | 11.80 | 13.27 | 19.48 | 14.94 | 14.38 | 22.08 | 45.97 |
| MNHN.1981.693 | *obstetricans* | 28.91 | 9.89 | 10.37 | 17.73 | 12.59 | 11.75 | 17.50 | 39.68 |
| MNHN.1993.5182 | *obstetricans* | 31.80 | 11.45 | 14.22 | 17.83 | 11.12 | 13.00 | 19.15 | 38.07 |

| | | | | | | | | | |
|---|---|---|---|---|---|---|---|---|---|
| MNHN.1998.0119 | *obstetricans* | 33.19 | 11.46 | 12.84 | 18.84 | 13.51 | 13.12 | 20.71 | 42.10 |
| MNHN.1998.0120 | *obstetricans* | 38.42 | 14.12 | 16.54 | 23.70 | 16.32 | 15.69 | 25.89 | 50.81 |
| MNHN.1998.0121 | *obstetricans* | 35.42 | 12.95 | 14.54 | 19.95 | 14.71 | 14.29 | 23.05 | 48.36 |
| MNHN.2000.0770 | *obstetricans* | 39.43 | 14.60 | 16.50 | 20.30 | 17.21 | 18.18 | 27.10 | 56.05 |
| MNHN.472.1 | *obstetricans* | 34.30 | 13.38 | 15.20 | 20.66 | 17.12 | 16.49 | 23.86 | 50.28 |
| MNHN.472.3 | *obstetricans* | 32.73 | 11.03 | 14.67 | 21.65 | 15.51 | 15.17 | 23.12 | 49.95 |
| NHM.1983.659 | *obstetricans* | 36.36 | 13.07 | 14.62 | 18.57 | 15.50 | 14.50 | 21.37 | 48.12 |
| NHM.1983.660. | *obstetricans* | 24.92 | 9.82 | 10.87 | 14.11 | 10.58 | 9.95 | 16.91 | 33.37 |
| NHM.1983.661 | *obstetricans* | 29.30 | 10.50 | 12.60 | 19.50 | 13.60 | 14.03 | 20.56 | 41.83 |
| NHM.1983.662 | *obstetricans* | 31.64 | 14.06 | 13.33 | 19.02 | 14.55 | 14.24 | 19.20 | 45.60 |
| NHM.1983.663 | *obstetricans* | 29.57 | 11.20 | 13.16 | 16.74 | 13.57 | 13.57 | 20.00 | 41.92 |
| NHM.1983.664 | *obstetricans* | 31.74 | 12.76 | 12.63 | 13.53 | 14.51 | 14.16 | 20.20 | 45.09 |
| NHM.1983.665 | *obstetricans* | 31.03 | 12.00 | 13.51 | 15.75 | 13.97 | 13.76 | 21.75 | 44.42 |
| NHM.1983.667 | *obstetricans* | 24.90 | 9.71 | 10.60 | 14.05 | 10.30 | 10.23 | 16.16 | 30.28 |
| NHM.1983.671 | *obstetricans* | 41.37 | 14.78 | 16.17 | 19.73 | 17.23 | 15.97 | 23.97 | 51.60 |
| NHM.1983.673 | *obstetricans* | 32.00 | 12.10 | 13.29 | 16.50 | 13.10 | 14.13 | 21.85 | 41.63 |
| NHM.1983.674 | *obstetricans* | 32.86 | 12.08 | 13.03 | 17.13 | 13.46 | 12.80 | 20.50 | 42.63 |
| NHM.1983.675 | *obstetricans* | 30.56 | 11.48 | 13.82 | 18.10 | 13.67 | 14.45 | 20.85 | 41.51 |
| NHM.1983.676 | *obstetricans* | 29.96 | 11.53 | 13.58 | 17.51 | 12.56 | 12.74 | 20.20 | 40.90 |
| NHM.1983.677 | *obstetricans* | 29.95 | 11.11 | 11.97 | 16.31 | 12.37 | 12.00 | 19.64 | 38.65 |
| NHM.1983.913 | *obstetricans* | 39.14 | 16.22 | 15.60 | 21.52 | 16.56 | 16.57 | 25.88 | 52.70 |
| NHM.1983.915 | *obstetricans* | 33.21 | 13.40 | 13.64 | 18.70 | 14.15 | 13.97 | 22.50 | 46.60 |
| ZFMK.23797 | *obstetricans* | 40.51 | 14.11 | 16.51 | 22.90 | 17.65 | 16.89 | 25.46 | 52.25 |
| ZFMK.23798 | *obstetricans* | 39.32 | 13.24 | 15.19 | 20.03 | 16.00 | 14.83 | 22.93 | 47.96 |
| ZFMK.23799 | *obstetricans* | 40.77 | 14.46 | 15.71 | 22.87 | 17.20 | 16.62 | 22.59 | 51.87 |
| ZFMK.23800 | *obstetricans* | 37.76 | 13.97 | 14.89 | 21.85 | 16.21 | 15.87 | 24.15 | 49.88 |
| ZFMK.23802 | *obstetricans* | 40.68 | 14.28 | 16.90 | 22.63 | 18.09 | 18.74 | 27.54 | 56.80 |
| ZFMK.23803 | *obstetricans* | 42.36 | 14.40 | 17.10 | 24.73 | 16.96 | 17.53 | 24.25 | 53.62 |
| ZFMK.23804 | *obstetricans* | 37.47 | 13.81 | 14.92 | 21.83 | 16.72 | 16.54 | 24.45 | 51.30 |
| ZFMK.27796 | *obstetricans* | 40.93 | 14.35 | 16.56 | 21.33 | 18.49 | 16.45 | 26.19 | 55.54 |
| ZFMK.30171 | *obstetricans* | 40.40 | 15.50 | 17.35 | 25.84 | 19.10 | 18.30 | 27.44 | 58.26 |
| ZFMK.30832 | *obstetricans* | 40.74 | 14.45 | 16.18 | 21.10 | 16.87 | 17.00 | 24.68 | 52.46 |

| | | | | | | | | | |
|---|---|---|---|---|---|---|---|---|---|
| ZFMK.30833 | *obstetricans* | 39.74 | 14.07 | 16.34 | 24.20 | 18.06 | 16.88 | 26.46 | 56.48 |
| ZFMK.42601 | *obstetricans* | 36.51 | 12.53 | 13.64 | 16.70 | 13.05 | 13.84 | 19.83 | 40.55 |
| ZFMK.45794 | *obstetricans* | 46.41 | 16.10 | 19.18 | 25.94 | 20.13 | 20.34 | 27.72 | 61.50 |
| ZFMK.47440 | *obstetricans* | 35.35 | 14.08 | 14.58 | 18.96 | 15.87 | 14.98 | 22.68 | 47.32 |
| ZFMK.61941 | *obstetricans* | 41.68 | 16.03 | 17.44 | 19.09 | 16.28 | 18.93 | 26.80 | 55.30 |
| ZFMK.63600 | *obstetricans* | 41.11 | 15.40 | 16.56 | 24.08 | 18.49 | 19.35 | 26.44 | 56.37 |
| ZFMK.72431 | *obstetricans* | 37.92 | 13.65 | 15.31 | 19.49 | 15.66 | 15.60 | 23.03 | 48.56 |
| ZFMK.99010 | *obstetricans* | 41.02 | 16.01 | 18.44 | 23.42 | 18.60 | 17.15 | 27.18 | 55.00 |
| ZFMK.99011 | *obstetricans* | 43.51 | 15.04 | 17.11 | 21.83 | 16.24 | 17.66 | 27.64 | 53.45 |
| MNCN.147 | *pertinax* | 32.42 | 11.77 | 12.80 | 16.92 | 12.30 | 12.52 | 18.88 | 39.15 |
| MNCN.180 | *pertinax* | 37.32 | 15.42 | 17.61 | 19.43 | 17.30 | 16.00 | 25.67 | 53.87 |
| MNCN.183 | *pertinax* | 35.11 | 14.07 | 16.48 | 19.08 | 16.63 | 15.18 | 22.24 | 50.63 |
| MNCN.184 | *pertinax* | 38.06 | 14.00 | 17.41 | 22.29 | 18.90 | 18.14 | 25.32 | 54.90 |
| MNCN.185 | *pertinax* | 41.11 | 15.11 | 16.96 | 19.58 | 16.75 | 16.96 | 24.87 | 52.30 |
| MNCN.210 | *pertinax* | 32.41 | 12.37 | 13.88 | 16.44 | 15.01 | 13.86 | 21.51 | 45.09 |
| MNCN.24087 | *pertinax* | 39.84 | 17.75 | 16.66 | 20.81 | 16.86 | 17.01 | 25.00 | 50.71 |
| MNCN.2604 | *pertinax* | 28.79 | 11.23 | 10.88 | 15.44 | 12.45 | 11.25 | 16.40 | 36.70 |
| MNCN.26123 | *pertinax* | 25.98 | 10.87 | 10.58 | 14.39 | 9.60 | 10.24 | 16.51 | 32.85 |
| MNCN.26169 | *pertinax* | 39.67 | 15.41 | 15.46 | 20.53 | 17.09 | 16.10 | 23.47 | 49.54 |
| MNCN.26170 | *pertinax* | 43.02 | 15.53 | 16.60 | 22.85 | 16.91 | 17.80 | 27.03 | 55.99 |
| MNCN.26171 | *pertinax* | 43.94 | 17.52 | 15.22 | 21.99 | 16.22 | 19.03 | 27.12 | 55.86 |
| MNCN.26172 | *pertinax* | 40.39 | 15.26 | 15.09 | 22.31 | 16.68 | 15.86 | 23.30 | 51.92 |
| MNCN.26173 | *pertinax* | 40.03 | 14.87 | 15.28 | 19.91 | 15.93 | 16.98 | 20.92 | 52.15 |
| MNCN.26174 | *pertinax* | 36.36 | 15.32 | 14.63 | 21.81 | 16.68 | 15.48 | 23.17 | 50.17 |
| MNCN.26175 | *pertinax* | 40.71 | 14.66 | 16.10 | 23.33 | 18.00 | 17.24 | 26.70 | 57.21 |
| MNCN.26176 | *pertinax* | 36.12 | 14.71 | 14.20 | 20.20 | 15.71 | 15.50 | 23.25 | 47.30 |
| MNCN.755 | *pertinax* | 37.25 | 15.50 | 16.42 | 22.25 | 17.52 | 15.82 | 25.83 | 51.97 |
| MNCN.756 | *pertinax* | 39.73 | 14.88 | 16.65 | 23.23 | 18.70 | 17.71 | 26.26 | 54.94 |
| MNCN.21653 | *maurus* | 41.94 | 14.88 | 15.64 | 23.02 | 16.73 | 16.61 | 25.85 | 52.52 |
| MNCN.21654 | *maurus* | 26.77 | 9.85 | 10.70 | 15.95 | 10.70 | 11.67 | 16.96 | 31.37 |
| MNHN.1908.111 | *maurus* | 29.75 | 11.05 | 12.48 | 15.85 | 11.04 | 11.53 | 18.43 | 35.33 |
| MNHN.1981.668 | *maurus* | 37.04 | 12.15 | 14.82 | 20.59 | 15.57 | 15.11 | 22.72 | 49.35 |

| | | | | | | | | | |
|---|---|---|---|---|---|---|---|---|---|
| MNHN.1981.669 | *maurus* | 36.25 | 13.09 | 14.26 | 20.33 | 15.32 | 14.90 | 21.87 | 46.75 |
| MNHN.1981.670 | *maurus* | 32.44 | 12.45 | 13.38 | 18.85 | 13.64 | 13.50 | 20.21 | 41.56 |
| MNHN.1981.673 | *maurus* | 34.27 | 11.40 | 13.68 | 19.23 | 14.48 | 14.39 | 21.75 | 43.81 |
| MNHN.1981.674 | *maurus* | 31.86 | 11.15 | 12.37 | 18.57 | 14.16 | 12.87 | 20.37 | 40.90 |
| MNHN.1981.675 | *maurus* | 31.74 | 9.77 | 12.37 | 17.79 | 11.53 | 13.49 | 18.95 | 38.94 |
| MNHN.1981.678 | *maurus* | 34.57 | 11.65 | 12.77 | 17.11 | 14.68 | 14.64 | 20.33 | 43.49 |
| MNHN.1981.680 | *maurus* | 32.10 | 11.98 | 12.52 | 17.70 | 11.80 | 13.53 | 20.77 | 40.43 |
| MNHN.1981.681 | *maurus* | 30.41 | 10.24 | 10.78 | 16.53 | 12.07 | 12.55 | 17.14 | 40.11 |
| MNHN.1981.682 | *maurus* | 32.65 | 11.27 | 11.62 | 17.06 | 13.08 | 13.14 | 19.61 | 37.60 |
| MNHN.1990.249 | *maurus* | 26.61 | 11.16 | 12.01 | 16.93 | 9.97 | 11.90 | 17.18 | 34.37 |
| MNHN.1994.1890 | *maurus* | 29.13 | 10.33 | 11.73 | 16.82 | 11.56 | 12.05 | 17.84 | 37.07 |
| MNHN.6892 | *maurus* | 31.45 | 12.02 | 12.10 | 18.71 | 13.66 | 13.45 | 17.44 | 39.37 |
| MNCN.24036 | *muletensis* | 22.85 | 8.12 | 8.80 | 14.77 | 9.23 | 9.82 | 14.40 | 30.04 |
| MNCN.24051 | *muletensis* | 23.69 | 8.76 | 9.55 | 13.76 | 8.55 | 9.12 | 14.48 | 28.59 |
| MNCN.24057 | *muletensis* | 24.14 | 9.01 | 9.07 | 13.57 | 10.11 | 10.98 | 13.88 | 32.44 |
| MNCN.24059 | *muletensis* | 20.76 | 8.36 | 9.02 | 13.67 | 9.14 | 9.59 | 12.79 | 29.01 |
| MNCN.42410 | *muletensis* | 27.27 | 11.17 | 12.18 | 15.53 | 12.41 | 12.85 | 19.72 | 41.62 |
| MNCN.42411 | *muletensis* | 30.25 | 11.50 | 13.18 | 20.43 | 11.00 | 13.41 | 19.95 | 40.77 |
| MNCN.42412 | *muletensis* | 28.38 | 10.10 | 11.40 | 18.98 | 13.03 | 14.29 | 19.12 | 39.60 |
| MNCN.42413 | *muletensis* | 27.10 | 10.88 | 10.61 | 18.76 | 11.11 | 12.47 | 18.11 | 37.32 |
| MNCN.46014 | *muletensis* | 31.68 | 11.96 | 12.59 | 22.05 | 14.83 | 14.95 | 21.20 | 43.01 |
| MNCN.46015 | *muletensis* | 27.90 | 10.63 | 10.69 | 17.58 | 13.40 | 12.93 | 17.70 | 38.19 |
| MNCN.46016 | *muletensis* | 29.27 | 11.70 | 11.80 | 16.32 | 13.00 | 12.98 | 20.15 | 41.55 |
| MNCN.46017 | *muletensis* | 32.09 | 11.55 | 12.20 | 20.36 | 16.34 | 14.11 | 19.96 | 43.05 |
| MNCN.46018 | *muletensis* | 27.18 | 10.47 | 11.14 | 16.53 | 11.55 | 12.49 | 17.87 | 39.16 |
| ZFMK.59084 | *muletensis* | 32.20 | 11.76 | 12.21 | 19.62 | 12.03 | 13.89 | 20.24 | 43.13 |
| MNCN.14017 | *dickhilleni* | 38.61 | 13.41 | 15.02 | 19.87 | 15.51 | 16.11 | 22.38 | 50.38 |
| MNCN.16682 | *dickhilleni* | 46.96 | 17.05 | 17.66 | 24.43 | 21.22 | 19.25 | 30.05 | 63.72 |
| MNCN.16683 | *dickhilleni* | 42.68 | 15.30 | 17.70 | 21.25 | 16.75 | 17.46 | 26.02 | 55.02 |
| MNCN.16684 | *dickhilleni* | 42.88 | 17.00 | 17.34 | 21.21 | 17.08 | 18.61 | 26.65 | 56.03 |
| MNCN.16685 | *dickhilleni* | 45.22 | 15.00 | 16.50 | 25.42 | 20.67 | 19.15 | 26.90 | 57.11 |
| MNCN.16739 | *dickhilleni* | 38.29 | 14.26 | 14.63 | 22.41 | 16.86 | 17.41 | 26.54 | 55.77 |

| | | | | | | | | | |
|---|---|---|---|---|---|---|---|---|---|
| MNCN.16740 | *dickhilleni* | 40.23 | 13.83 | 15.58 | 23.21 | 18.14 | 18.06 | 26.00 | 55.81 |
| MNCN.16741 | *dickhilleni* | 37.65 | 13.12 | 14.05 | 20.09 | 17.07 | 10.10 | 24.44 | 52.10 |
| MNCN.16742 | *dickhilleni* | 41.70 | 15.06 | 17.11 | 20.57 | 18.45 | 18.10 | 25.26 | 57.47 |
| MNCN.20765 | *dickhilleni* | 47.86 | 14.79 | 18.26 | 23.46 | 17.45 | 18.00 | 26.31 | 53.47 |
| MNCN.20767 | *dickhilleni* | 47.62 | 15.05 | 17.72 | 17.71 | 19.38 | 17.84 | 21.79 | 51.42 |
| MNCN.20797 | *dickhilleni* | 39.38 | 16.69 | 16.47 | 22.38 | 18.55 | 18.35 | 26.40 | 56.14 |
| MNCN.26116 | *dickhilleni* | 41.89 | 15.86 | 16.95 | 23.06 | 17.43 | 18.02 | 25.49 | 58.25 |
| MNCN.26117 | *dickhilleni* | 40.67 | 15.05 | 16.78 | 23.38 | 16.78 | 17.27 | 25.49 | 55.17 |
| MNCN.26119 | *dickhilleni* | 37.80 | 14.70 | 15.12 | 19.68 | 18.47 | 17.77 | 26.39 | 50.60 |
| MNCN.26126 | *dickhilleni* | 42.11 | 15.55 | 16.47 | 22.43 | 18.15 | 19.90 | 25.81 | 53.04 |
| MNCN.26129 | *dickhilleni* | 43.71 | 16.38 | 17.19 | 24.15 | 20.65 | 19.04 | 28.15 | 59.58 |
| MNCN.26130 | *dickhilleni* | 38.67 | 14.12 | 14.87 | 20.27 | 16.65 | 16.27 | 23.92 | 51.68 |
| MNCN.26131 | *dickhilleni* | 42.24 | 15.68 | 17.02 | 21.20 | 15.75 | 17.78 | 25.90 | 51.91 |
| MNCN.26141 | *dickhilleni* | 36.18 | 13.32 | 14.79 | 16.16 | 15.98 | 14.48 | 23.35 | 45.80 |
| MNCN.26142 | *dickhilleni* | 39.84 | 14.76 | 15.71 | 21.25 | 16.53 | 16.66 | 24.90 | 51.70 |
| MNCN.26143 | *dickhilleni* | 38.44 | 14.46 | 16.33 | 19.72 | 17.03 | 16.06 | 24.33 | 49.61 |
| MNCN.26144 | *dickhilleni* | 37.21 | 13.96 | 16.35 | 19.29 | 16.81 | 16.08 | 23.28 | 47.04 |
| MNCN.26145 | *dickhilleni* | 40.82 | 14.33 | 16.06 | 20.17 | 17.75 | 16.60 | 23.79 | 51.77 |
| MNCN.42245 | *dickhilleni* | 43.43 | 14.68 | 17.60 | 24.64 | 18.51 | 18.73 | 27.34 | 57.68 |
| MNCN.211 | *cisternasii* | 32.27 | 10.11 | 14.91 | 14.12 | 13.25 | 11.90 | 19.13 | 39.41 |
| MNCN.212 | *cisternasii* | 33.12 | 10.44 | 14.92 | 15.31 | 14.01 | 11.74 | 20.08 | 40.49 |
| MNCN.213 | *cisternasii* | 31.63 | 10.91 | 14.56 | 14.95 | 13.78 | 12.60 | 19.11 | 40.09 |
| MNCN.214 | *cisternasii* | 33.07 | 10.75 | 15.10 | 12.42 | 12.33 | 11.42 | 18.61 | 37.57 |
| MNCN.215 | *cisternasii* | 30.40 | 11.14 | 15.19 | 13.98 | 13.10 | 12.14 | 17.86 | 38.32 |
| MNCN.216 | *cisternasii* | 24.76 | 10.30 | 12.15 | 13.47 | 12.01 | 10.69 | 13.56 | 34.22 |
| MNCN.26013 | *cisternasii* | 40.18 | 12.96 | 15.04 | 16.02 | 15.42 | 13.05 | 21.98 | 45.22 |
| MNCN.26034 | *cisternasii* | 32.66 | 11.45 | 14.78 | 18.25 | 17.06 | 14.28 | 19.18 | 45.46 |
| MNCN.26035 | *cisternasii* | 32.77 | 11.19 | 13.94 | 13.26 | 13.39 | 12.86 | 18.20 | 40.11 |
| MNCN.26036 | *cisternasii* | 35.37 | 11.99 | 15.44 | 15.88 | 11.97 | 14.27 | 20.77 | 39.29 |
| MNCN.26037 | *cisternasii* | 26.54 | 9.98 | 13.34 | 13.21 | 12.11 | 11.50 | 16.02 | 32.28 |
| MNCN.26038 | *cisternasii* | 35.94 | 12.43 | 16.13 | 15.20 | 16.39 | 14.09 | 21.11 | 46.30 |
| MNCN.26039 | *cisternasii* | 31.60 | 10.59 | 13.37 | 14.90 | 13.59 | 12.73 | 19.56 | 36.85 |

| | | | | | | | | | |
|---|---|---|---|---|---|---|---|---|---|
| MNCN.26040 | *cisternasii* | 35.32 | 11.78 | 14.52 | 16.60 | 15.24 | 14.11 | 20.59 | 44.30 |
| MNCN.26042 | *cisternasii* | 36.78 | 12.20 | 15.47 | 16.75 | 15.17 | 13.40 | 21.63 | 42.83 |
| MNCN.26044 | *cisternasii* | 30.34 | 11.51 | 13.92 | 13.19 | 13.05 | 11.60 | 19.92 | 37.10 |
| MNCN.26045 | *cisternasii* | 28.07 | 10.29 | 12.05 | 12.09 | 11.53 | 11.31 | 16.91 | 34.74 |
| MNCN.26046 | *cisternasii* | 33.41 | 11.80 | 14.66 | 14.56 | 14.55 | 13.30 | 19.40 | 41.44 |
| MNCN.26047 | *cisternasii* | 31.28 | 11.95 | 14.46 | 13.36 | 13.47 | 12.50 | 19.46 | 38.79 |
| MNCN.26048 | *cisternasii* | 32.39 | 11.43 | 14.33 | 16.13 | 12.91 | 12.78 | 19.44 | 38.32 |
| MNCN.26049 | *cisternasii* | 24.14 | 9.10 | 10.54 | 10.80 | 10.16 | 9.85 | 12.58 | 28.36 |
| MNCN.26052 | *cisternasii* | 36.33 | 12.48 | 16.43 | 15.55 | 14.55 | 13.88 | 22.13 | 45.10 |
| MNCN.26053 | *cisternasii* | 39.27 | 13.11 | 16.90 | 16.08 | 16.15 | 14.59 | 22.96 | 46.27 |
| MNCN.26054 | *cisternasii* | 33.56 | 11.20 | 14.91 | 14.18 | 12.79 | 12.36 | 19.35 | 38.45 |
| MNCN.26055 | *cisternasii* | 30.78 | 10.95 | 12.24 | 14.84 | 14.08 | 11.51 | 16.92 | 37.44 |
| MNCN.30762 | *cisternasii* | 28.62 | 9.43 | 11.61 | 12.50 | 11.67 | 11.44 | 14.91 | 33.22 |
| MNCN.30763 | *cisternasii* | 31.67 | 8.96 | 12.73 | 12.49 | 12.46 | 11.33 | 16.95 | 36.00 |
| MNCN.30764 | *cisternasii* | 27.65 | 8.64 | 11.64 | 12.72 | 11.67 | 11.16 | 15.89 | 34.08 |
| MNCN.46559 | *cisternasii* | 33.00 | 11.80 | 13.97 | 15.85 | 15.39 | 13.61 | 19.29 | 44.65 |
| MNCN.46560 | *cisternasii* | 34.30 | 11.37 | 13.72 | 15.03 | 12.66 | 13.09 | 19.58 | 39.27 |
| MNCN.46561 | *cisternasii* | 32.21 | 11.55 | 13.10 | 15.64 | 11.79 | 11.71 | 20.00 | 40.03 |
| MNCN.46562 | *cisternasii* | 31.85 | 10.94 | 13.70 | 14.30 | 11.46 | 13.05 | 18.36 | 37.60 |
| MNCN.46563 | *cisternasii* | 30.41 | 10.94 | 13.34 | 15.07 | 12.04 | 12.76 | 18.25 | 40.04 |
| MNCN.46564 | *cisternasii* | 36.97 | 13.41 | 14.28 | 15.77 | 11.41 | 13.31 | 18.71 | 38.75 |
| MNCN.46565 | *cisternasii* | 35.45 | 11.66 | 14.73 | 15.41 | 13.32 | 12.33 | 19.30 | 38.80 |
| NHM.1980.252 | *cisternasii* | 29.97 | 11.90 | 13.34 | 16.54 | 14.14 | 12.13 | 19.09 | 40.76 |

**File S4**: Characteristics and variables of the species distribution models.

| | ***almogavarii*** | ***inigoi*** | ***boscai*** | ***lusitanicus*** | ***obstetricans*** | ***pertinax*** | ***dickhilleni*** | ***maurus*** | ***muletensis*** | ***cisternasii*** |
|---|---|---|---|---|---|---|---|---|---|---|
| **Model characteristics** | | | | | | | | | | |
| Total number of localities | 828 | 15 | 241 | 261 | 3757 | 763 | 376 | 108 | 22 | 715 |
| Number of localities without duplicates | 759 | 14 | 186 | 228 | 2170 | 706 | 302 | 90 | 12 | 564 |
| Partial ROC | 0 | 0 | 0 | 0 | 0 | 0 | 0 | 0 | 0 | 0 |
| Omission rate 5% | 0.048 | 0.125 | 0.066 | 0.038 | 0.056 | 0.05 | 0.048 | 0.037 | 0.182 | 0.047 |
| AICc | 18,729 | 457 | 5,868 | 6,051 | 113,718 | 18,514 | 8,224 | 2,312 | 504 | 17,971 |
| Delta AICc | 0 | 0 | 0 | 0 | 0 | 0 | 0 | 0 | 0 | 0 |
| Significant models (omission rate criterion) | 40 | 0 | 0 | 192 | 0 | 80 | 215 | 22 | 0 | 277 |
| Significant models (AICc criterion) | 1 | 2 | 1 | 2 | 2 | 1 | 4 | 1 | 3 | 1 |
| Significant models (both criteria) | 1 | 0 | 0 | 2 | 0 | 1 | 1 | 1 | 0 | 1 |
| Regularization multiplier | 2.5 | 6.0* | 3.5 | 0.5 | 4 | 1.5 | 2.5 | 0.5 | 2.5 | 1.5 |
| Response type of feature classes | lqh | th* | q | lq | lqp | pt | pth | lq | lth | lqth |
| Average AUC | 0.985 | 1 | 0.99 | 0.992 | 0.87 | 0.982 | 0.993 | 0.998 | 1 | 0.981 |
| Standard deviation AUC | 0 | 0 | 0 | 0 | 0.002 | 0 | 0 | 0 | 0 | 0.001 |
| Average TSS | 0.972 | 0.989 | 0.952 | 0.953 | 0.501 | 0.931 | 0.973 | 0.963 | 0.986 | 0.916 |
| Standard deviation TSS | 0.006 | 0.023 | 0.005 | 0.011 | 0.008 | 0.007 | 0.008 | 0.014 | 0.026 | 0.005 |
| **Model variables** | | | | | | | | | | |
| Annual mean temperature (Bio1) | 0.1 | 0.1 | 0 | 0 | 1.2 | 0.2 | 0 | 0 | 0 | 0.1 |
| Mean diurnal range (Bio2) | 0.2 | 0.1 | 0 | 0.1 | 2.2 | 2.9 | 28.1 | 2.7 | 3.7 | 0.8 |
| Isothermality (Bio3) | 13.8 | 9.8 | 15.1 | 0.1 | 4.1 | 13.6 | 7.4 | 0.6 | 0 | 1.4 |
| Temperature seasonality (Bio4) | 18.3 | 7.6 | 1 | 0.5 | 6.3 | 3.1 | 0.9 | 0.1 | 0 | 0.1 |
| Maximum temperature of warmest month (Bio5) | 0.1 | 0 | 0 | 0 | 2.3 | 0 | 0.6 | 0 | 0 | 0 |
| Minimum temperature of coldest month (Bio6) | 19.1 | 0.5 | 0 | 0.8 | 1.7 | 0.2 | 0 | 0.1 | 6.6 | 0.3 |
| Temperature annual range (Bio7) | 5.9 | 0.1 | 0.1 | 0.3 | 1 | 2.1 | 4.8 | 0.7 | 3.1 | 0.7 |

| | | | | | | | | | | |
|---|---|---|---|---|---|---|---|---|---|---|
| Mean temperature of wettest quarter (Bio8) | 3 | 2.3 | 5.5 | 2.8 | 0.4 | 1.5 | 3 | 0.3 | 0 | 5.4 |
| Mean temperature of driest quarter (Bio9) | 0 | 10.4 | 0.2 | 0.1 | 2.7 | 0.2 | 0 | 0.1 | 1.1 | 0.1 |
| Mean temperature of warmest quarter (Bio10) | 0 | 0 | 0 | 0 | 1 | 0.4 | 1.2 | 0 | 0 | 0 |
| Mean temperature of coldest quarter (Bio11) | 1.3 | 15.1 | 0.1 | 0.3 | 0.2 | 0.8 | 0 | 0.1 | 0 | 0.1 |
| Annual precipitation (Bio12) | 0 | 0 | 0 | 0.1 | 0.2 | 0.1 | 0 | 0.1 | 0 | 0.6 |
| Precipitation of wettest month (Bio13) | 0 | 0 | 1.5 | 2.5 | 0.1 | 0 | 0.2 | 0.3 | 0.8 | 0.8 |
| Precipitation of driest month (Bio14) | 19 | 0 | 6.1 | 12.9 | 1.1 | 3.6 | 0 | 0.7 | 9.8 | 19.6 |
| Precipitation seasonality (Bio15) | 4.4 | 16.2 | 0.7 | 6.9 | 45.4 | 8.6 | 4.1 | 37.3 | 0.4 | 15.6 |
| Precipitation of wettest quarter (Bio16) | 0.5 | 0 | 0.1 | 1.7 | 0.4 | 1.4 | 1.2 | 1.2 | 0 | 0.1 |
| Precipitation of driest quarter (Bio17) | 0.2 | 0 | 0.1 | 0.1 | 3.3 | 2.9 | 0 | 1.3 | 0 | 0.9 |
| Precipitation of warmest quarter (Bio18) | 0.6 | 0 | 1.6 | 2.8 | 0.9 | 24.7 | 0 | 2.7 | 5.9 | 9.5 |
| Precipitation of coldest quarter (Bio19) | 0.2 | 4 | 51.4 | 43.2 | 0 | 10.3 | 7.6 | 8 | 0.8 | 14.5 |
| Broadleaf forest | 0 | 2.6 | 0.2 | 0.2 | 0.1 | 0.2 | 0 | 1.2 | 4.2 | 0 |
| Needleleaf forest | 0 | 0.9 | 0.8 | 0.2 | 0.2 | 0 | 0.1 | 1.4 | 0.9 | 0.1 |
| Mixed forest | 3.2 | 1.5 | 0 | 1 | 0.7 | 0.1 | 0.1 | 0.1 | 4.8 | 14.2 |
| Shrubs | 0 | 0.6 | 0.6 | 1 | 3.9 | 10.1 | 11.7 | 0.6 | 16.7 | 0.1 |
| Barren | 0 | 1.4 | 3.7 | 4.4 | 1.3 | 0.1 | 0.1 | 13.9 | 4.3 | 1.6 |
| Herbaceous vegetation | 0 | 3.9 | 3.3 | 0.9 | 1.8 | 0.1 | 0 | 0.4 | 3.5 | 0.3 |
| Cultivated vegetation | 0.6 | 0.1 | 5 | 5.1 | 1.6 | 0.4 | 0.2 | 4.6 | 0.3 | 0.5 |
| Altitude | 3.6 | 10.7 | 1.8 | 0.8 | 0.3 | 1.1 | 15.4 | 10.5 | 0.1 | 0.3 |
| Aridity index | 0 | 0 | 0 | 0 | 0.6 | 5.1 | 0 | 0 | 0 | 9 |
| Global land cover | 0.1 | 0.9 | 0.6 | 7.8 | 12.6 | 0.2 | 0.9 | 6.4 | 3.4 | 0.4 |
| Habitat homogeneity | 0 | 0.1 | 0 | 0.2 | 0.1 | 0.1 | 0.1 | 0.1 | 0 | 0.7 |
| Hillshade | 0 | 0.2 | 0.1 | 0.1 | 0.2 | 0.1 | 0 | 0.9 | 3 | 0.2 |
| Slope | 0 | 0.1 | 0 | 0 | 1.5 | 0.2 | 0.1 | 0 | 0 | 0.5 |
| Terrain ruggedness index | 5.7 | 0.6 | 0 | 0.8 | 0 | 5.4 | 9.8 | 1.8 | 26.5 | 0.2 |
| Topographic position index | 0 | 5.1 | 0.1 | 2.2 | 0 | 0.1 | 0 | 1.6 | 0 | 1.1 |
| Tree coverage percent | 0 | 5.2 | 0 | 0.2 | 0.7 | 0 | 2.1 | 0.4 | 0 | 0.3 |

* parameter settings proposed by *kuenm* did not allow us to build an acceptable model of the taxon and the model was built with default settings (regularization multiplier is 1.0 and response type of feature classes is lqph).

**File S5**: STRUCTURE analyses in the *A. obstetricans* complex for K = 2–6.

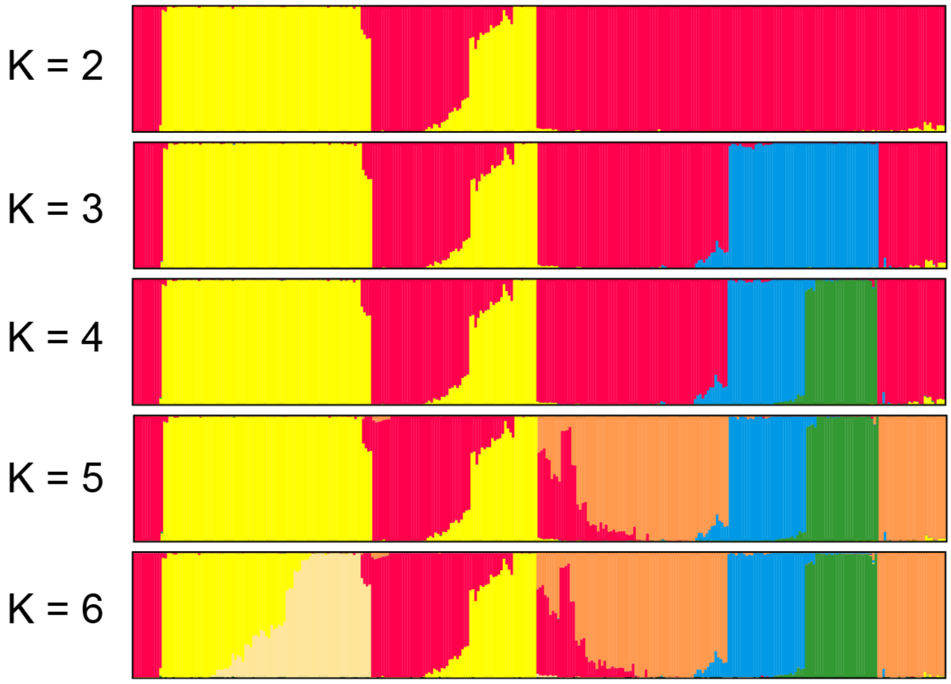

**File S6:** Relationship between genome average cline width and the median of cline widths obtained from individual loci. The link is significant (linear regression, $R^2 = 0.99$, $P < 0.05$; dashed line).

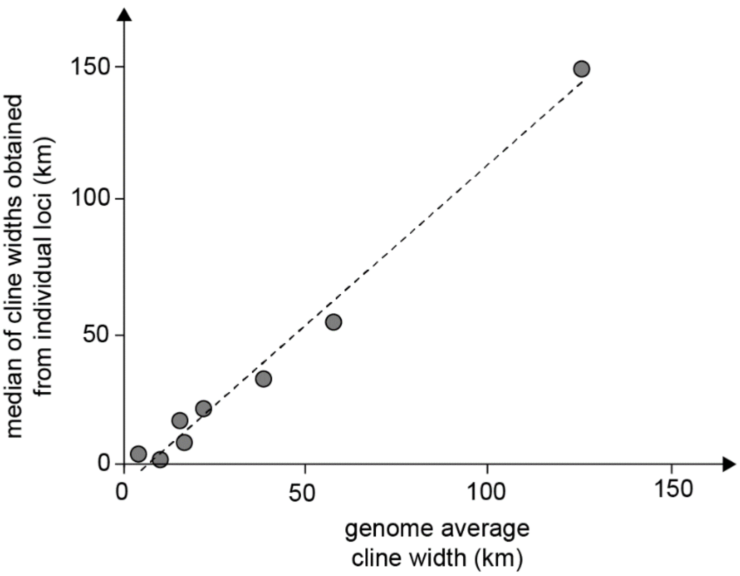

**File S7:** Pairwise genomic divergence (net sequence divergence) based on ddRAD-seq loci used in phylogenomic analyses.

| | ***boscai*** | ***lusitanicus*** | ***almogavarii*** | ***inigoi*** | ***obstetricans*** | ***pertinax*** | ***cisternasii*** | ***dickhilleni*** | ***maurus*** | ***muletensis*** |
|---|---|---|---|---|---|---|---|---|---|---|
| ***boscai*** | - | | | | | | | | | |
| ***lusitanicus*** | 0.19% | - | | | | | | | | |
| ***almogavarii*** | 0.38% | 0.33% | - | | | | | | | |
| ***inigoi*** | 0.41% | 0.36% | 0.18% | - | | | | | | |
| ***obstetricans*** | 0.30% | 0.29% | 0.33% | 0.36% | - | | | | | |
| ***pertinax*** | 0.25% | 0.22% | 0.24% | 0.28% | 0.14% | - | | | | |
| ***cisternasii*** | 0.58% | 0.49% | 0.61% | 0.62% | 0.61% | 0.54% | - | | | |
| ***dickhilleni*** | 0.88% | 0.73% | 0.88% | 0.90% | 0.91% | 0.82% | 0.58% | - | | |
| ***maurus*** | 0.83% | 0.69% | 0.84% | 0.86% | 0.86% | 0.78% | 0.55% | 0.57% | - | |
| ***muletensis*** | 0.58% | 0.49% | 0.58% | 0.61% | 0.61% | 0.54% | 0.39% | 0.37% | 0.38% | - |

**File S8:** Pairwise bioacoustic differentiation (Euclidian distances).

| | ***boscai*** | ***lusitanicus*** | ***almogavarii*** | ***inigoi*** | ***obstetricans*** | ***pertinax*** | ***cisternasii*** | ***dickhilleni*** | ***maurus*** | ***muletensis*** |
|---|---|---|---|---|---|---|---|---|---|---|
| ***boscai*** | - | | | | | | | | | |
| ***lusitanicus*** | 0.23 | - | | | | | | | | |
| ***almogavarii*** | 1.06 | 0.84 | - | | | | | | | |
| ***inigoi*** | 1.28 | 1.36 | 2.06 | - | | | | | | |
| ***obstetricans*** | 0.60 | 0.52 | 0.85 | 1.46 | - | | | | | |
| ***pertinax*** | 1.01 | 0.85 | 0.46 | 2.19 | 1.01 | - | | | | |
| ***cisternasii*** | 2.49 | 2.35 | 2.21 | 2.24 | 2.07 | 2.66 | - | | | |
| ***dickhilleni*** | 5.10 | 4.99 | 4.89 | 4.54 | 4.65 | 5.33 | 2.72 | - | | |
| ***maurus*** | NA | NA | NA | NA | NA | NA | NA | NA | - | |
| ***muletensis*** | 3.48 | 3.29 | 2.51 | 4.54 | 3.31 | 2.47 | 3.99 | 6.27 | NA | - |

**File S9:** Pairwise morphological differentiation after allometric size correction (Euclidian distances).

| | ***boscai*** | ***lusitanicus*** | ***almogavarii*** | ***inigoi*** | ***obstetricans*** | ***pertinax*** | ***cisternasii*** | ***dickhilleni*** | ***maurus*** | ***muletensis*** |
|---|---|---|---|---|---|---|---|---|---|---|
| ***boscai*** | - | | | | | | | | | |
| ***lusitanicus*** | 1.23 | - | | | | | | | | |
| ***almogavarii*** | 1.16 | 2.23 | - | | | | | | | |
| ***inigoi*** | 0.70 | 1.81 | 0.61 | - | | | | | | |
| ***obstetricans*** | 1.14 | 0.98 | 1.79 | 1.50 | - | | | | | |
| ***pertinax*** | 0.51 | 1.14 | 1.18 | 0.92 | 0.87 | - | | | | |
| ***cisternasii*** | 4.37 | 3.30 | 5.31 | 4.94 | 3.55 | 4.18 | - | | | |
| ***dickhilleni*** | 0.58 | 1.67 | 1.12 | 0.76 | 1.40 | 0.89 | 4.62 | - | | |
| ***maurus*** | 2.02 | 1.26 | 2.77 | 2.47 | 1.02 | 1.76 | 2.62 | 2.31 | - | |
| ***muletensis*** | 4.87 | 3.82 | 5.58 | 5.32 | 3.87 | 4.58 | 1.71 | 5.20 | 2.91 | - |

**File S10**: Niche overlap between reconstructed distribution models (Schoener's *D*).

| | ***boscai*** | ***lusitanicus*** | ***almogavarii*** | ***inigoi*** | ***obstetricans*** | ***pertinax*** | ***cisternasii*** | ***dickhilleni*** | ***maurus*** | ***muletensis*** |
|---|---|---|---|---|---|---|---|---|---|---|
| ***boscai*** | - | | | | | | | | | |
| ***lusitanicus*** | 0.46 | - | | | | | | | | |
| ***almogavarii*** | 0.06 | 0.02 | - | | | | | | | |
| ***inigoi*** | 0.07 | 0.00 | 0.08 | - | | | | | | |
| ***obstetricans*** | 0.18 | 0.03 | 0.25 | 0.22 | - | | | | | |
| ***pertinax*** | 0.10 | 0.05 | 0.16 | 0.12 | 0.17 | - | | | | |
| ***cisternasii*** | 0.35 | 0.47 | 0.02 | 0.01 | 0.04 | 0.07 | - | | | |
| ***dickhilleni*** | 0.13 | 0.13 | 0.03 | 0.02 | 0.04 | 0.11 | 0.21 | - | | |
| ***maurus*** | 0.10 | 0.08 | 0.00 | 0.00 | 0.01 | 0.02 | 0.10 | 0.12 | - | |
| ***muletensis*** | 0.13 | 0.14 | 0.13 | 0.01 | 0.08 | 0.16 | 0.15 | 0.07 | 0.06 | - |

**File S11:** Pairwise environmental differentiation based on the raw geo-climatic data (Euclidian distances).

| | ***boscai*** | ***lusitanicus*** | ***almogavarii*** | ***inigoi*** | ***obstetricans*** | ***pertinax*** | ***cisternasii*** | ***dickhilleni*** | ***maurus*** | ***muletensis*** |
|---|---|---|---|---|---|---|---|---|---|---|
| ***boscai*** | - | | | | | | | | | |
| ***lusitanicus*** | 2.00 | - | | | | | | | | |
| ***almogavarii*** | 6.54 | 6.51 | - | | | | | | | |
| ***inigoi*** | 7.27 | 7.71 | 5.55 | - | | | | | | |
| ***obstetricans*** | 6.86 | 7.31 | 3.78 | 3.66 | - | | | | | |
| ***pertinax*** | 6.82 | 6.54 | 3.98 | 5.56 | 4.79 | - | | | | |
| ***cisternasii*** | 5.37 | 4.39 | 6.05 | 8.20 | 7.22 | 4.34 | - | | | |
| ***dickhilleni*** | 8.55 | 7.69 | 7.35 | 7.19 | 7.48 | 4.33 | 5.12 | - | | |
| ***maurus*** | 5.81 | 4.92 | 8.12 | 8.62 | 8.66 | 6.16 | 3.77 | 5.16 | - | |
| ***muletensis*** | 6.16 | 5.60 | 4.45 | 7.63 | 6.75 | 4.17 | 4.56 | 6.79 | 6.51 | - |

**File S12:** Relationship (log-transformed) between pairwise niche dissimilarity (1-Schoener's *D*) of the reconstructed models and environmental differentiation (Euclidian distances). The link is not significant (Mantel test, $P = 0.98$).

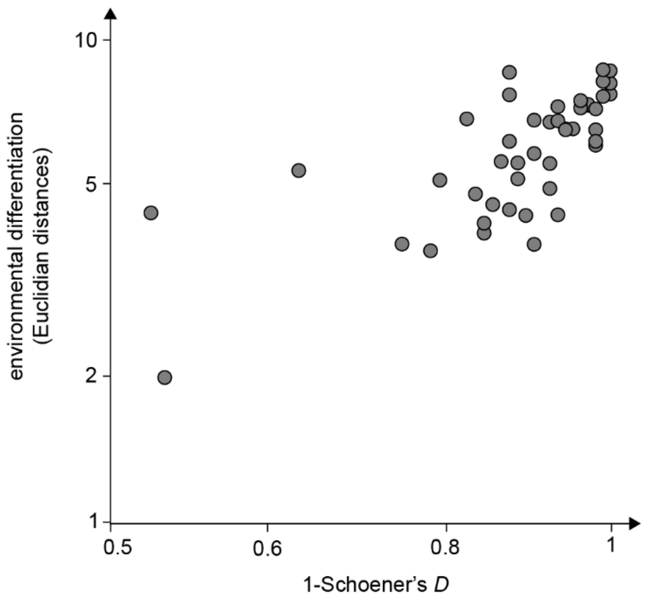

**File S13**: Relationship (log-transformed) between pairwise environmental differentiation and spatial distances (Euclidian distances). The link is significant (Mantel test, $P < 0.001$; dashed line).

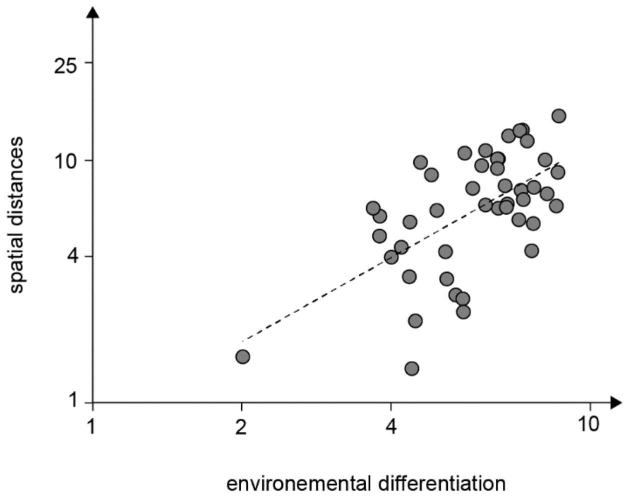

**File S14**: Genome average clines (sigmoid lines) and their confidence interval (colored areas) overlayed by the occurrence probabilities of the sampled populations obtained by the ecological niche models (dots) for the two lineages of each contact zone; vertical lines: genome average cline centers *c*.

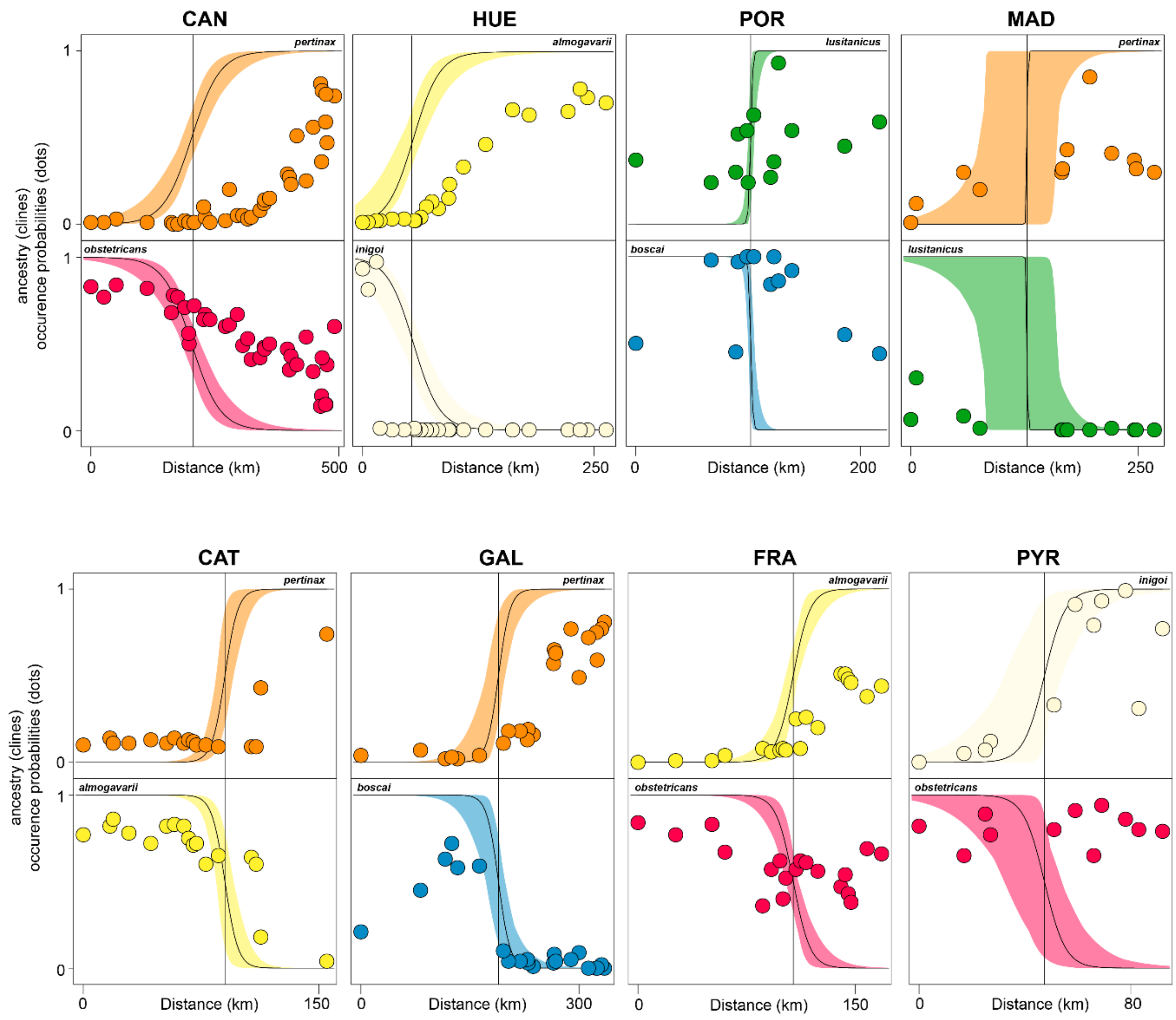
